# Candidate methylation sites associated with endocrine therapy resistance in the TCGA ER+/HER2- breast cancer cohort

**DOI:** 10.1101/826198

**Authors:** Maryam Soleimani, Simone Borgoni, Emre Sofyalı, Pernette J. Verschure, Stefan Wiemann, Perry D. Moerland, Antoine H.C. van Kampen

**Author notes:** These authors contributed equally to this work (Perry D. Moerland, Antoine H.C. van Kampen). Corresponding author: Antoine H.C. van Kampen, PhD Meibergdreef 9, 1105 AZ Amsterdam, the Netherlands.

## Abstract

**Background:** Estrogen receptor (ER) positive breast cancer is often effectively treated with drugs that inhibit ER signaling, i.e., tamoxifen (TAM) and aromatase inhibitors (AIs). However, about 30% of ER+ breast cancer patients develop resistance to therapy leading to tumour recurrence. Changes in the methylation profile have been implicated as one of the mechanisms through which therapy resistance develops. Therefore, we aimed to identify methylation loci associated with endocrine therapy resistance.

**Methods:** We used genome-wide DNA methylation profiles of primary ER+ tumors from The Cancer Genome Atlas in combination with curated data on survival and treatment to predict development of endocrine resistance. Association of individual DNA methylation markers with survival was assessed using Cox proportional hazards models in a cohort of ER+/HER2- tumours (*N*=552) and two sub-cohorts corresponding to the endocrine treatment (AIs or TAM) that patients received (*N*=210 and *N*=172, respectively). Models were adjusted for clinical variables tumour stage, age, AI treatment and luminal subtype. We also identified signatures of multiple methylation loci associated with survival using Cox proportional hazards models with elastic net regularization. Individual markers and multivariable signatures were compared with DNA methylation profiles generated in a time course experiment using the T47D ER+ breast cancer cell line treated with tamoxifen or deprived from estrogen.

**Results:** We identified 132, 9 and 1 CpGs for which DNA methylation is significantly associated with survival in the ER+/HER2-, TAM and AI cohorts respectively. Corresponding multi-locus signatures consisted of 171, 50 and 160 CpGs and showed a large overlap with the corresponding single-locus signatures. Single-locus signatures for the ER+/HER2- and TAM cohorts were conserved among the loci that were differentially methylated in endocrine-resistant T47D cells. Similarly, multi-locus signatures for the ER+/HER2- and AI cohorts were conserved in endocrine-resistant T47D cells.

**Conclusions:** We identified individual and multivariable DNA methylation markers associated with therapy resistance independently of luminal status. Our results suggest that these markers identified from primary tumours and prior to any endocrine treatment are associated with development of endocrine resistance.

## Introduction

Breast cancer is among the most common cancers diagnosed in women in Europe where it also is the third cause of cancer death after lung and colorectal cancer (1). Approximately 75% of breast cancers is characterized by the expression of estrogen receptor alpha (ERα), encoded by the estrogen receptor 1 (*ESR1*) gene. These tumours require estrogen signals for continued growth and, consequently, patients generally receive endocrine treatment to inhibit ER signalling (2) Endocrine treatment comprises selective estrogen receptor modulators, including tamoxifen, selective estrogen receptor down-regulators including fulvestrant, and aromatase inhibitors (e.g., anastrozole, letrozole and exemestane) that inhibit the production of estrogen from androgen. Unfortunately, resistance to endocrine therapy develops in approximately 30% of ER+ breast cancer patients resulting in recurrence of the tumour (3). Despite many efforts the precise mechanisms leading to acquired treatment resistance remain mostly unknown and, hence, therapies to prevent or revert resistance are currently lacking. Therefore, the identification of biomarkers, including epigenetic markers, that can predict endocrine resistance are considered of great value for patient stratification prior to endocrine therapy (4).

In general, breast cancer development, progression and (endocrine) drug resistance result from the cumulative burden of genetic and epigenetic changes. Moreover, post-transcriptional and post-translational modifications are likely to contribute as well (5–7).

The association of epigenetic changes with tumour characteristics, subtypes, prognosis, and treatment outcome is not well characterized (8). Epigenetic changes have been shown to drive resistance acquisition through their effect on gene expression and/or chromosomal stability (9). For example, using RNA-seq and ChIP-seq analysis of the acetylation of lysine 27 on histone 3 (H3K27ac), an established active enhancer marker, revealed that epigenetic activation of the cholesterol biosynthesis pathway causes activation of ERα resulting in resistance(10). DNA methylation has also been shown to be perturbed during breast cancer development and may largely affect gene expression (4, 11) Since DNA methylation has also been shown to be altered in endocrine resistant tumours (12) the identification of methylation markers for disease diagnosis, prognosis, and treatment outcome is receiving increased attention. Moreover, breast cancer treatment might benefit from the regulation of methylation activity by using DNA methyltransferase inhibitors (4). Treatment with the DNA methyltransferase inhibitor 5-aza-2’ deoxycytidine caused a significant reduction in promoter methylation and a concurrent increase in expression of the gene *ZNF350* that encodes a DNA damage response protein, and of *MAGED1* which is a tumour antigen and putative regulator of P53, suggesting that a methylation-targeted therapy might be beneficial (13). However, current inhibitors have weak stability, lack specificity for cancer cells and are inactivated by cytidine deaminase thus limiting their use in the treatment of breast cancer (14).

Several studies investigated DNA methylation in relation to disease outcome and therapy resistance. Lin et al. observed significant differences in DNA methylation profiles between tamoxifen sensitive and tamoxifen resistant cell lines (15). There, a large number of genes, several of which have been previously implicated in breast cancer pathogenesis, were shown to have increased DNA methylation of their promoter CpG islands in the resistant cell lines. Similarly, Williams et al. observed a large number of hypermethylated genes in a tamoxifen-resistant cell line (13). In a meta-analysis of two human breast cancer gene expression datasets, 144 abnormally methylated genes were shortlisted as putative epigenetic biomarkers of survival. Kaplan-Meier survival analysis on the expression of these genes further reduced this list to 48 genes, and a subsequent correlation analysis of gene expression and DNA methylation provided evidence for the potential association of DNA methylation with survival in different breast cancer subtypes including ER+/HER2- (16). Another study compared ductal carcinoma in situ to invasive breast cancer and suggested that methylation changes indicate an early event in the progression of cancer and, therefore, might be of relevance for clinical decision making (17). In contrast to studies that showed the impact of promoter methylation, it has also been demonstrated that endocrine response in cell lines is mainly modulated by methylation of estrogen-responsive enhancers (18). There, increased *ESR1*-responsive enhancer methylation in primary tumours was found to be associated with endocrine resistance and disease relapse in ER-positive (luminal A) human breast cancer, suggesting that methylation levels can be used to identify patients that positively respond to endocrine therapy. Note that, although limited ER-responsive enhancer methylation may already be present in the primary tumour, the analysis of methylation profiles of matched relapse samples showed that enhancer DNA methylation increased during treatment. Therefore, a combination of pre-existing and acquired differences in enhancer DNA methylation could be associated with the development of endocrine therapy resistance.

In the current work we investigated if DNA methylation profiles of primary ER+/HER2- tumours provide information to predict endocrine resistance. We selected methylation profiles provided by The Cancer Genome Atlas (TCGA) (19) from patients treated with tamoxifen or aromatase inhibitors, and assumed that patient survival is a proxy for absence of therapy resistance. To identify specific DNA methylation markers we tested the association with survival using a Cox proportional hazards model. We were able to identify DNA methylation markers associated with patient outcome. We validated these markers using DNA methylation profiles generated in a time course experiment using the T47D cell line treated with tamoxifen or deprived from estrogen.

## Materials and methods

### Data

We used clinical, biospecimen, gene expression (RNAseq V2) and DNA methylation (Illumina Human Methylation 450K) data of 1,098 patients with breast invasive carcinoma (BRCA) from TCGA (cancergenome.nih.gov). Samples represented in TCGA were all collected prior to adjuvant therapy (20). TCGA also recorded patient follow-up information describing clinical events such as type of treatment, the number of days from the date of initial pathological diagnosis to a new tumour event, death, and date of last contact. Since clinical and biospecimen data are scattered over multiple files in the TCGA repository, we first merged all information in a single table with one row per patient using the patient identifiers provided in the clinical and biospecimen data. Subsequently, we corrected drug names for tamoxifen and aromatase inhibitors (AIs; anastrozole, exemestane and letrozole) for spelling variants and mapped synonyms to their generic drug names (Additional File 1). We selected the subset of patients (samples) that were treated with AI or tamoxifen.

### Patient cohorts

For all patients with DNA methylation data available we selected data from primary tumours (indicated with “01” in the patient barcode) of female ER+/HER2- BRCA patients (Figure 1). The molecular subtype was determined using TCGA gene expression data for these samples (see below). The ER+/HER2- cohort was further subdivided according to the endocrine treatment (AI or tamoxifen) patients received during follow-up. Patients who received both drugs were included in both sub-cohorts. Consequently, we considered three patient cohorts, i.e., ER+/HER2-, AI, and tamoxifen (TAM).

**Figure 1.**
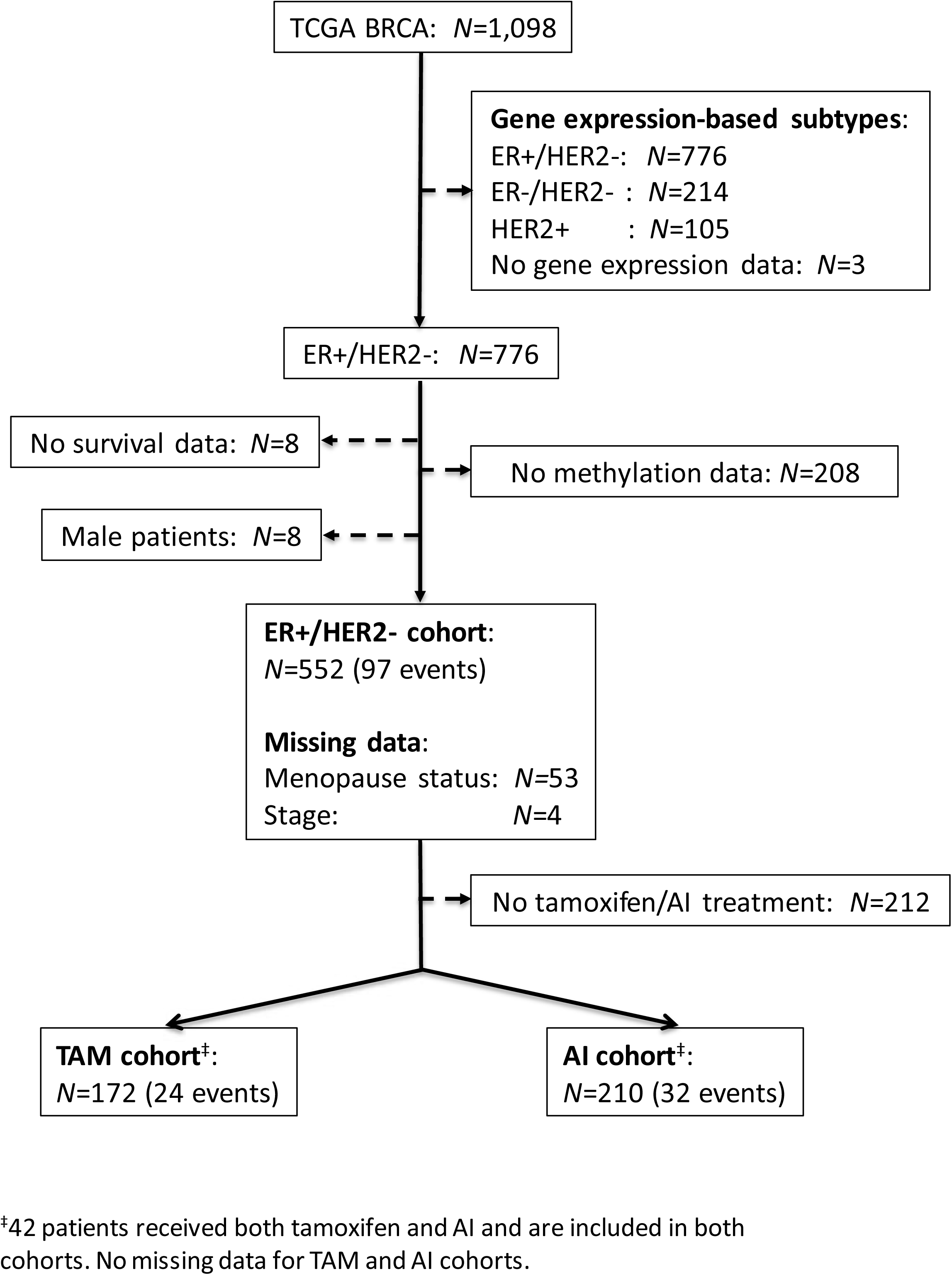
Study flow chart and cohort definition. This figure shows the steps taken to define each of the three cohorts. First the molecular subtype was determined and ER+/HER2- patients were selected. Next, patients without follow-up data and patients for whom no methylation profiles were measured were removed. Finally, male patients were removed leading to the study cohort of ER+/HER2- patients. Patients who received tamoxifen form the TAM sub-cohort and patients who received AI form the AI sub-cohort, 42 patients are included in both the TAM and AI sub-cohort. Dashed arrows indicate filter steps. AI, aromatase inhibitor; BRCA, breast invasive carcinoma; ER, estrogen receptor; HER2, human epidermal growth factor receptor 2; TAM, tamoxifen; TCGA, The Cancer Genome Atlas.

### Subtype determination

Information for BRCA subtyping by immunohistochemistry of ER or HER2 is missing for 192 out of 1,098 patients. Therefore, we used TCGA BRCA RNAseq V2 gene expression data to determine molecular subtypes (Additional File 2). To this end, gene expression data from primary tumours were retrieved from the Genomic Data Commons legacy archive (https://portal.gdc.cancer.gov/legacy-archive) using the R package *TCGAbiolinks* (21). RSEM estimated abundances were normalised using the upper quartile method from the R package *edgeR* (22) and subsequently log2-transformed with an offset of one. Breast cancer subtypes ER-/HER2-, HER2+, and the lowly proliferative ER+/HER2- (luminal A) and highly proliferative ER+/HER2- (luminal B) subtypes were determined using the SCMOD2 model from the R package *genefu* (23).

### DNA methylation data and pre-processing

Illumina Human Methylation 450K raw data (IDAT files) for the patients in the cohorts defined above were retrieved from TCGA. Pre-processing was performed using the R package *minfi* (24). Detection p-values were calculated for each methylation probe. 82,150 probes showed an unreliable signal (p>0.01) in one or more samples and were removed. Data were normalized using functional normalization (25). Probes corresponding to loci that contain a SNP in the CpG site or in the single-base extension site were removed. We also removed probes that have been shown to cross-hybridize to multiple genomic positions (26). Finally, M-values were calculated and probes with low variation across samples (standard deviation of M-values ≤0.4) were removed. The final data set comprised 322,426 CpG loci. Probes were annotated to genes and enhancer regions using the R package *IlluminaHumanMethylation450kanno.ilmn12.hg19*.

### Survival analysis

#### Clinical variables

Based on literature (27–29) we selected menopause status (pre/post, after merging pre- and peri-menopausal; values ‘[Unknown]’ and ‘Indeterminate’ were considered missing), AI treatment (yes, no), tamoxifen treatment (yes, no), tumour stage (I-IV, after merging subcategories; stage X was considered missing), and age at diagnosis as candidate variables predictive of survival. We tested association with survival using the Cox proportional hazards model (R package *survival*). We defined an event as the first occurrence of a new tumour event or death. For patients without an event we used the latest contact date as provided by the clinical data (right censoring). To account for missing values for the variables menopause status and stage we used multiple imputation (R package *mice*) to generate 50 datasets and perform survival analysis on each dataset separately (30). The input data used for multiple imputation is available in Additional File 3. Rubin’s rule was applied to combine individual estimates and standard errors (SEs) of the model coefficients from each of the imputed datasets into an overall estimate and SE resulting in a single p-value for each variable. Clinical variables with a p-value <0.10 in a univariable survival model were selected for inclusion in the multivariable survival model. Variables in the final multivariable model were determined using backward selection by iteratively removing variables with the highest p-value until all variables had a p-value <0.05.

#### Single-locus survival analysis

Next we performed survival analysis to identify single methylation loci associated with patient survival using the methylation M-values in a Cox proportional hazards model. The models for each locus were adjusted for significant clinical variables from the multivariable model. To account for missing values for clinical variables, multiple imputation was used as described above. Resulting p-values were corrected for multiple testing using the Benjamini-Hochberg false discovery rate (FDR). Corrected p-values <0.05 were considered statistically significant. Subsequently, single-locus survival models were also adjusted for ER+/HER2- subtypes (luminal A/luminal B) in addition to the clinical variables selected above. Kaplan-Meier curves for individual loci were determined by calculating the median of the methylation levels over all patients in a cohort and then assigning a patient to a low (methylation level < median) and a high (methylation level ≥ median) group.

#### Multi-locus survival analysis

We used the Cox proportional hazards model with elastic net regularization (function cv.glmnet, R package *glmnet*) (31) to identify a signature of multiple methylation loci associated with survival. We followed a two-stage approach. First, the CpG signature was determined without including clinical variables using Cox regression with elastic net penalty. Secondly, from the resulting model the risk score (see below) was calculated and used in a new model that includes the clinical variables selected above in order to establish whether the methylation signature provided additional information compared to merely using clinical variables. Optimal values, minimizing the partial likelihood deviance, for the elastic net mixing parameter (α) and tuning parameter (λ) were determined by stratified (for event status) 10-fold cross-validation using a grid search varying α from 0 to 1 in steps of 0.1 and using 100 values for λ that were automatically generated for each α. We constructed one model for each of the three cohorts (ER+/HER2-, AI, TAM). Subsequently, for each cohort we used the identified signature to calculate a risk score for each patient:

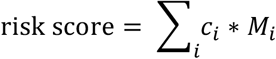

where for CpG locus *i, c*_*i*_ denotes the corresponding coefficient in the Cox model and *M*_*i*_ the methylation M-value. Next, multivariable Cox proportional hazards regression was performed using the risk score as a variable and adjusting for significant clinical variables from the multivariable model. Missing values for the clinical variables were imputed as described above. Finally, the risk-score-based models were also adjusted for ER+/HER2- subtypes (luminal A/luminal B) in addition to the selected clinical variables. Kaplan-Meier curves were determined for two groups of patients by calculating the median of the risk scores over all patients in a cohort and then assigning a patient to a good (risk score < median) and a bad prognosis group (risk score ≥ median).

#### Stability of multi-locus signatures

To assess the stability of the multi-locus signatures 50 regularized Cox models were fitted using a stratified (for event status) selection of 90% of the samples for each cohort. We counted the number of times each CpG locus was included in the 50 signatures and then selected those CpGs that occurred in at least 10 or at least 35 signatures. We refer to the resulting signatures as stability signatures. Fisher’s exact test was used to determine the significance of the overlap between the original multi-locus signature and the stability signatures.

### Methylation profiling of resistance acquisition in an ER+ breast cancer cell line

T47D cells were either treated with 100 nM 4-hydroxytamoxifen (TMX), long-term estrogen deprived (LTED; modelling AI treatment) (32) or not treated (wild type (WT)) in two biological replicates cultured for 7 and 5 months, respectively. DNA was extracted after 0, 1, 2, 5 and 7 (only one replicate) months. Methylation profiling was performed using the Illumina MethylationEPIC BeadChip platform at the Genomic and Proteomic Core Facility (DKFZ, Germany). For each sample two technical replicates were measured. Pre-processing was performed as described above. 8,682 probes showed an unreliable signal (detection p-value >0.01) in one or more samples and were removed. Probes that cross-hybridized to multiple genomic positions as listed by Pidsley et al. (33) were removed. No filtering based on M-values was performed. The final data set contains 786,500 CpG loci. Using the resulting M-values CpG-wise linear models were fitted with coefficients for each treatment (TMX, LTED, WT) and time point combination. In addition, we included a coefficient to correct for systematic differences between the two biological replicates (R package *limma*). For both LTED and TMX treated samples, contrasts were made between each individual time point *t* and the WT cell line at baseline, that is, LTED_*t*_ – WT_0_ and TMX_*t*_ – WT_0_, respectively. Differential methylation was assessed using empirical Bayes moderated statistics while also including the consensus correlation within pairs of technical replicates in the linear model fit (function duplicateCorrelation, *limma* package).

### Enrichment analysis

We tested whether the methylation loci identified from the TCGA BRCA single-locus and multi-locus signatures (based on Illumina 450k arrays) and also represented on the Illumina EPIC array were enriched in the T47D resistance acquisition experiment using ROAST rotation-based gene set tests (*limma* package) (34). Enrichment of TAM and AI survival signatures was assessed using the comparison of TMX and LTED treated cells to WT baseline via the linear model and contrasts described above. Enrichment of the ER+/HER2- survival signature was assessed using the comparison of pooled TMX and LTED treated cells versus WT baseline via the contrast (LTED_*t*_ + TMX_*t*_)/2 – WT_0_. ROAST p-values were calculated, for two alternative hypotheses denoted as ‘up’ and ‘down’ using 9999 rotations. In the ROAST analyses directional contribution weights of 1 or −1 were used depending on whether a CpG of the signature under consideration had a positive (corresponding to increased risk of an event) or negative (corresponding to decreased risk of an event) coefficient in the corresponding Cox model. In this case, the alternative hypothesis ‘up’ corresponds to methylation levels changing in the same direction in the TCGA BRCA survival signature and in the resistance acquisition experiment, whereas the alternative hypothesis ‘down’ corresponds to a change in the opposite direction (Figure 2). The two-sided directional p-value is reported.

**Figure 2.**
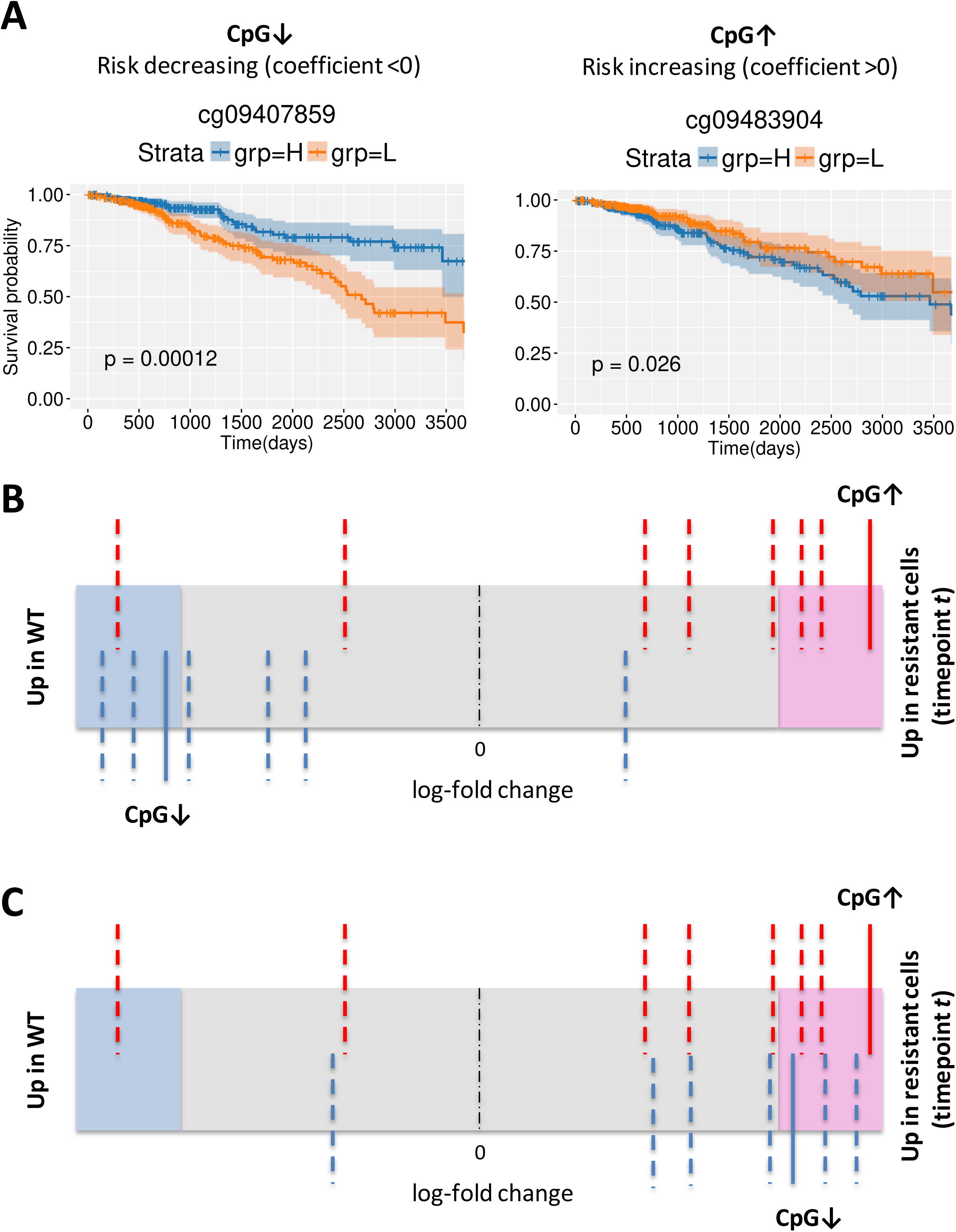
Validation of survival signatures in resistance acquisition experiment. (A) Kaplan-Meier plots for two selected CpGs significantly associated with survival in the ER+/HER2- cohort. Patients were stratified based on the methylation levels of risk decreasing locus CpG↓ (left; higher methylation is associated with longer survival) and a risk increasing locus CpG↑ (right; higher methylation is associated with shorter survival). H, methylation levels above median; L, methylation levels below median. Shaded areas denote the 95% confidence interval in the H and L strata. P-values are based on a log-rank test. (B) Example of a barcode enrichment plot for a TCGA BRCA survival signature in the cell line comparison of treated (LTED or TMX) samples at time point *t* versus WT baseline. All methylation loci are ranked from left to right by increasing log-fold change in the cell line comparison under consideration and represented by a shaded bar. Loci within the survival signature are represented by vertical bars. Red bars correspond to risk increasing loci (for example, CpG↑), blue bars correspond to risk decreasing loci (for example, CpG↓). In this example, the risk increasing loci tend to be hypermethylated in the treated cell line and the risk decreasing loci tend to be hypomethylated. That is, most loci change in the same direction in the survival signature and the resistance acquisition experiment. (C) When using directional weights of 1 and −1 for risk increasing and risk decreasing loci respectively, the blue bars are reflected across the dashed line at a log-fold-change of 0. In this case for a ROAST gene set test, the alternative hypothesis ‘up’ corresponds to methylation levels changing in the same direction whereas the alternative hypothesis ‘down’ corresponds to a change in the opposite direction.

## Results

### Clinical variables are associated with survival in ER+/HER2- cohort

For the TCGA BRCA ER+/HER2- cohort (N=552, Figure 1) we assessed whether the clinical variables menopause status, AI treatment, tamoxifen treatment, tumour stage and age at diagnosis were associated with survival, with an event defined as first occurrence of a new tumour event or death. In a univariable Cox proportional hazards model tumour stage (HR 1.92, 95% CI 1.43-2.59; p=1.63E-05) and age at diagnosis (HR 1.03, 95% CI 1.01-1.05; p=2.40E-04) are significantly associated with survival (Table 1A). This is in agreement with previous findings that a more advanced tumour stage and increased age are associated with poorer outcome (35).Tamoxifen treatment, AI treatment and menopause status are not significantly associated with survival in our cohort. When we included the clinical variables in a multivariable Cox proportional hazards model, tumour stage, age and AI treatment were selected for inclusion in the final multivariable model using backward selection (Table 1B).

**Table 1A.**
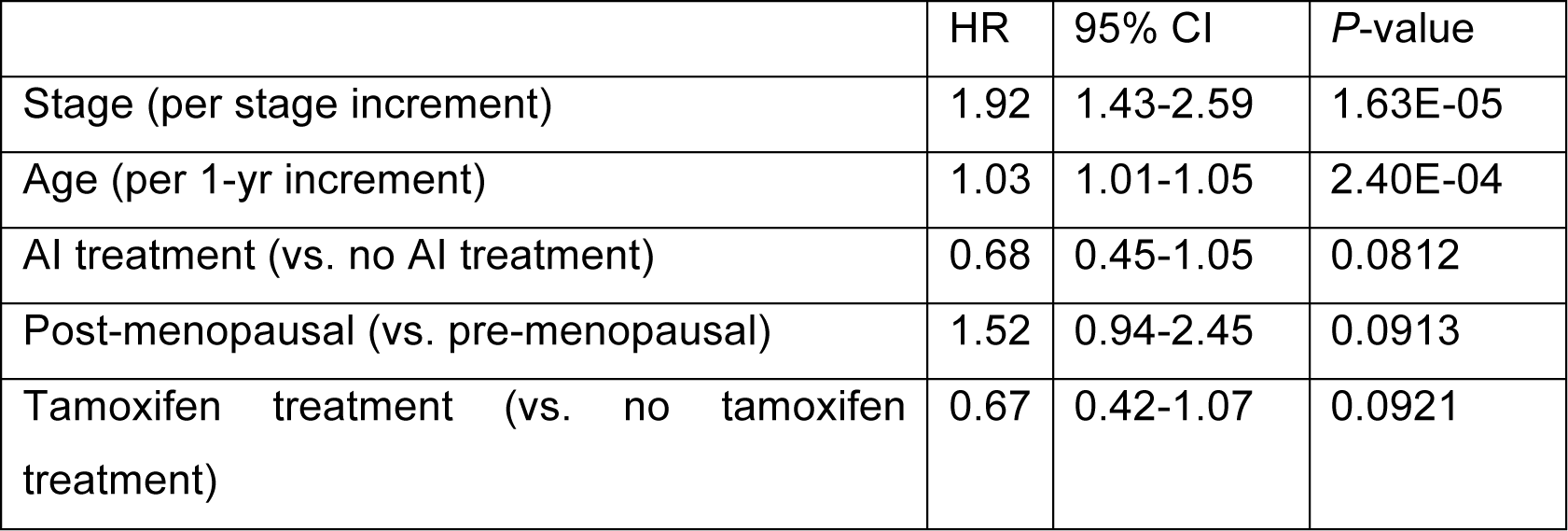
Univariable Cox proportional hazards model for clinical variables (ER+/HER2- cohort). HR, hazard ratio; CI, confidence interval; AI: aromatase inhibitor.

**Table 1B.**
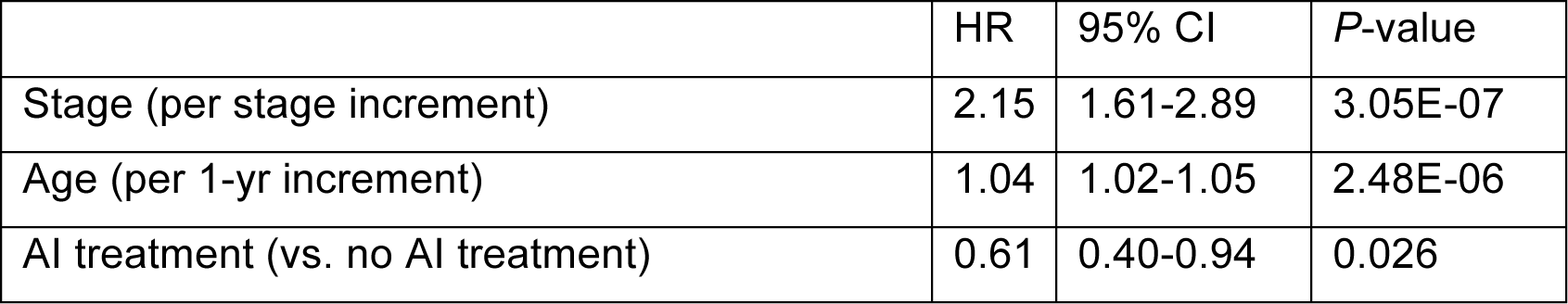
Multivariable Cox proportional hazards model for clinical variables (ER+/HER2- cohort). HR, hazard ratio; CI, confidence interval; AI: aromatase inhibitor.

|  | HR | 95% CI | *P*-value |
| --- | --- | --- | --- |
| Stage (per stage increment) | 2.15 | 1.61-2.89 | 3.05E-07 |
| Age (per 1-yr increment) | 1.04 | 1.02-1.05 | 2.48E-06 |
| AI treatment (vs. no AI treatment) | 0.61 | 0.40-0.94 | 0.026 |

### Single methylation loci associated with survival

To identify individual methylation loci associated with survival we fitted 322,426 Cox proportional hazard models using the M-value of each CpG while adjusting for the clinical variables selected in the multivariable model above (tumour stage, age and AI treatment (ER+/HER2- cohort only)). This resulted in 132, 9 and 1 CpGs for which DNA methylation is significantly (adjusted p-value <0.05) associated with survival in the ER+/HER2-, TAM, and AI cohort respectively (Additional File 4). The Kaplan-Meier curves show a significant difference in survival between the two groups stratified on median methylation level for nearly all selected loci (Additional File 5). Interestingly, apart from one CpG in the ER+/HER2- signature, for all of the CpGs increased methylation is associated with decreased risk of an event. Additional File 6 shows the overlap of the signatures for the three cohorts. Four out of nine methylation loci from the TAM signature are also found in the ER+/HER2- signature and, consequently, the other five loci are specific for tamoxifen treated patients.. Since all patients in the TAM cohort are also included in the ER+/HER2- cohort, overlap between the signatures is expected. TAM and AI signatures do not share methylation loci. ER+/HER2- and TAM signatures are enriched for enhancer CpGs (ER+/HER2-: 37%, p=0.0006; TAM: 55%, p=0.039; Fisher’s exact test).

### Multi-locus methylation signature associated with survival

Next we performed a multivariable analysis with elastic net penalty to find combinations of methylation loci associated with survival in a Cox proportional hazards model, This resulted in 171, 50 and 160 CpGs that are included in the survival signatures of the ER+/HER2-, TAM, and AI cohort respectively (Additional File 7). The ER+/HER2- and TAM signatures are enriched for enhancer loci (ER+/HER2-: 42%, p=1.60E-09; TAM: 38%, p=0.008; Fisher’s exact test). The risk score calculated from the multi-locus signature and adjusted for tumour stage, age and AI treatment (ER+/HER2- cohort only) is significantly associated with survival (p<10E-5) for all three cohorts (Additional File 8) indicating that DNA methylation is an independent factor in predicting survival. The risk scores calculated from the multi-locus signatures stratify the patients in two groups for each cohort (Figure 3A).

**Figure 3.**
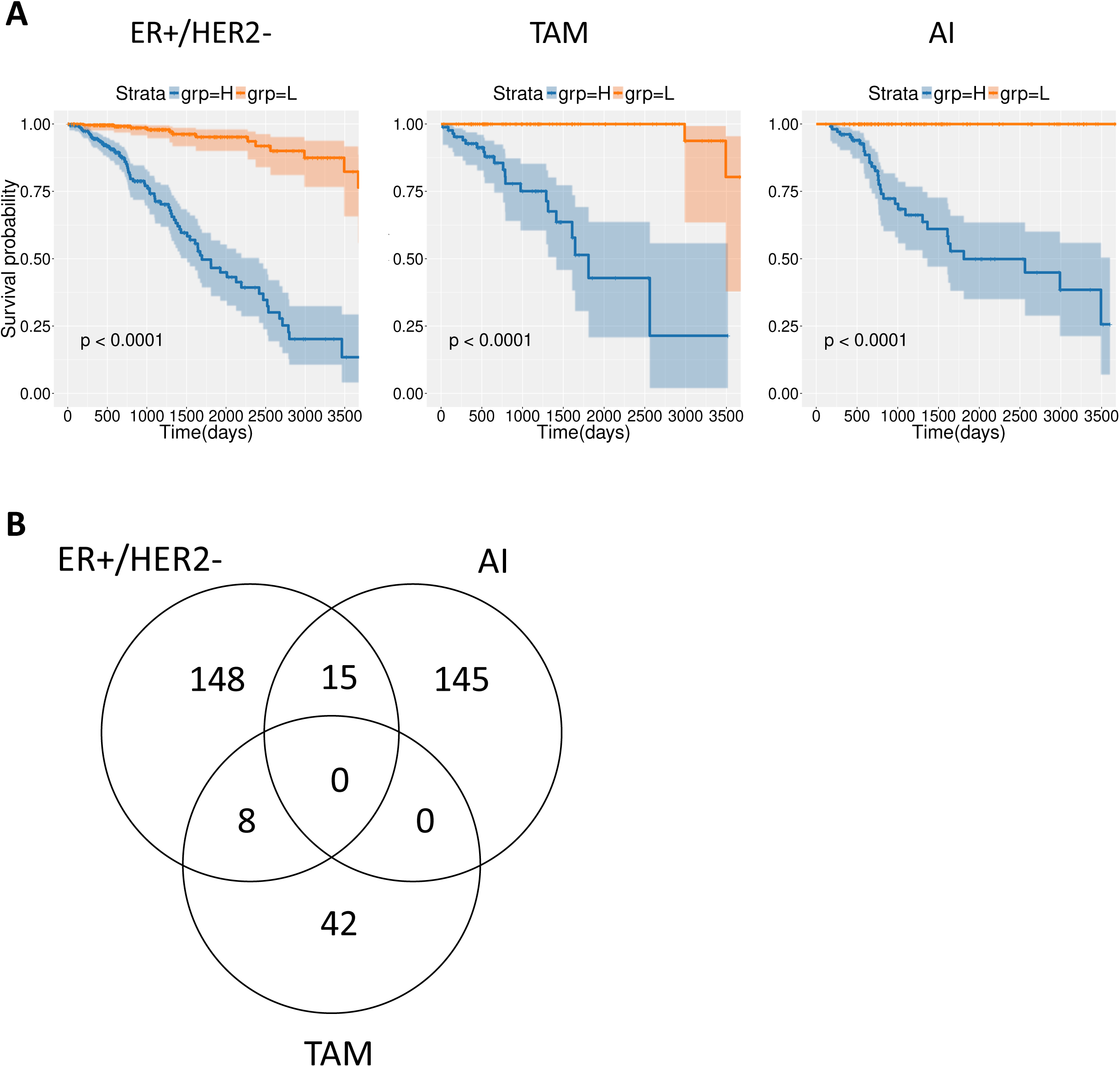
Multi-locus survival analysis. (A) Kaplan-Meier plots of the patients stratified based on the risk scores of the multi-locus signature in ER+/HER2, TAM and AI cohorts. H, risk score above median; L, risk score below median. Shaded areas denote the 95% confidence interval in the H and L strata. P-values are based on a log-rank test. (B) Venn diagram denoting the number of methylation loci in the multi-locus signatures for the ER+/HER2-, TAM, and AI cohorts.

There is no overlap between the signatures of TAM and AI cohorts. However, the ER+/HER2- signature partly overlaps with the TAM and AI signatures (Figure 3B). The coefficients in the Cox models corresponding to the overlapping loci have an identical sign in both cohorts. The multi-locus signatures include a large number of methylation loci that were also identified in the corresponding single-locus survival analysis. 41 out of 171 methylation loci in the ER+/HER2- multi-locus signature were also found in the single-locus signature (Additional File 9). Moreover, all methylation loci in the TAM and AI single-locus signatures, nine and one respectively, are part of the corresponding multi-locus signature.

We assessed the stability of the multi-locus signatures using a 10% leave-out test. The stability signature was enriched in the original multi-locus signature for each corresponding cohort (p<1E-3; Additional File 10).

### Validation of survival signatures in T47D resistance acquisition experiment

The single-locus survival signatures for ER+/HER2- and TAM are significantly enriched in the comparison of the last time point (7 months) versus WT baseline in a resistance acquisition experiment using T47D cells (ER+/HER2-: p=2.5E-3, TAM: p=2.9E-3; direction: ‘up’; Table 2). The signatures are not enriched at earlier time points. However, the proportion of CpGs contributing to enrichment in the same direction (‘up’) increases over time until it becomes significant for the last time point. The single-locus AI signature consists of only one CpG and an enrichment analysis is therefore not possible. However, for this locus the change in methylation level when comparing LTED treated cells with WT baseline is not concordant with the log-hazard ratio for that locus (data not shown).

**Table 2.** ROAST test results for the single-locus signatures. Direction indicates the direction of change. Methylation loci were weighted by their direction of change in the survival signature. ‘Up’ therefore corresponds to changes in the same direction in the survival signature and in the resistance acquisition experiment. That is, if a locus is risk in/decreasing in the survival signature than it is hyper/hypomethylated in the cell line signature for the indicated time point as compared to WT baseline. ‘Down’ corresponds to changes in the opposite direction. Prop., proportion of loci in the signature contributing to the estimated p-value and direction. Significant p-values (<0.05) are indicated in bold.

| Time point | ER+/HER2- (126 CpG sites) |  |  |  | TAM (8 CpG sites) |  |  |  |
| --- | --- | --- | --- | --- | --- | --- | --- | --- |
|  | Direction | P-value | Prop. (down) | Prop. (up) | Direction | P-value | Prop. (down) | Prop. (up) |
| 1 | Down | 0.9371 | 0.0952 | 0.071 | Down | 0.7520 | 0 | 0 |
| 2 | Up | 0.1685 | 0.1270 | 0.206 | Up | 0.8089 | 0 | 0 |
| 5 | Down | 0.2796 | 0.2778 | 0.270 | Up | 0.1184 | 0.125 | 0.25 |
| 7 | Up | **0.0025** | 0.2222 | 0.357 | Up | **0.0029** | 0 | 0.5 |

The multi-locus survival signatures for ER+/HER2 and AI are also significantly enriched at the 7-month time point in the resistance acquisition experiment (Table 3). The multi-locus TAM signature is not significantly enriched at any time point.

**Table 3.**
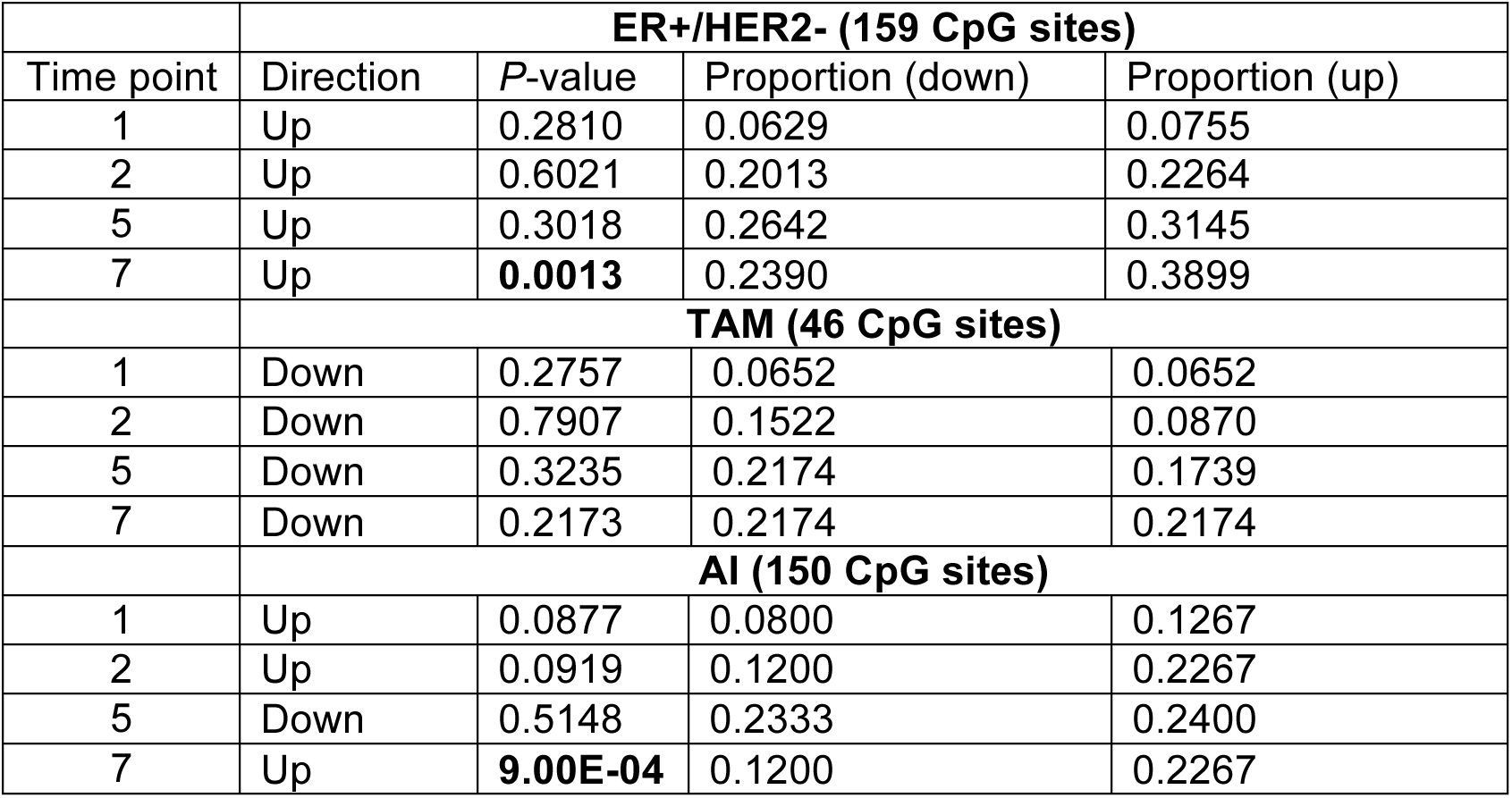
ROAST test results for the multi-locus signatures. Direction indicates the direction of change. Methylation loci were weighted by their direction of change in the survival signature. ‘Up’ therefore corresponds to changes in the same direction in the survival signature and in the resistance acquisition experiment. That is, if a locus is risk in/decreasing in the survival signature than it is hyper/hypomethylated in the cell line signature for the indicated time point as compared to WT baseline. ‘Down’ corresponds to changes in the opposite direction. Prop., proportion of loci in the signature contributing to the estimated p-value and direction. Significant p-values (<0.05) are indicated in bold.

| **ER+/HER2- (159 CpG sites)** |  |  |  |  |
| --- | --- | --- | --- | --- |
| Time point | Direction | P-value | Proportion (down) | Proportion (up) |
| 1 | Up | 0.2810 | 0.0629 | 0.0755 |
| 2 | Up | 0.6021 | 0.2013 | 0.2264 |
| 5 | Up | 0.3018 | 0.2642 | 0.3145 |
| 7 | Up | **0.0013** | 0.2390 | 0.3899 |
| **TAM (46 CpG sites)** |  |  |  |  |
| 1 | Down | 0.2757 | 0.0652 | 0.0652 |
| 2 | Down | 0.7907 | 0.1522 | 0.0870 |
| 5 | Down | 0.3235 | 0.2174 | 0.1739 |
| 7 | Down | 0.2173 | 0.2174 | 0.2174 |
| **AI (150 CpG sites)** |  |  |  |  |
| 1 | Up | 0.0877 | 0.0800 | 0.1267 |
| 2 | Up | 0.0919 | 0.1200 | 0.2267 |
| 5 | Down | 0.5148 | 0.2333 | 0.2400 |
| 7 | Up | **9.00E-04** | 0.1200 | 0.2267 |

## Discussion

We investigated whether TCGA DNA methylation profiles measured in primary ER+/HER2- tumours can be used to predict development of resistance to endocrine therapy in two sub-cohorts of patients treated with tamoxifen or AI. Using a single-locus Cox proportional hazard model we were able to identify 132, 9 and 1 CpGs for which DNA methylation is significantly associated with survival in the ER+/HER2-, TAM and AI cohorts respectively, while the corresponding multi-locus signatures consisted of 171, 50 and 160 CpGs. The multi-locus signatures showed a large overlap of 31%, 100%, and 100% with the ER+/HER2-, TAM and AI single-locus signatures respectively. The risk scores of the multi-locus signatures were significantly associated with survival. Moreover, we found that the ER+/HER2- and TAM single-locus and multi-locus signatures were significantly enriched for CpGs in enhancer regions suggesting a functional effect (on gene expression) (18). For both the single-locus signatures (Additional File 6) and the multi-locus signatures (Figure 3A) we observed no overlap of loci associated with survival between the AI and TAM cohorts. This could be indicative of a difference in development of resistance against tamoxifen or AI. This is in line with earlier observations in endocrine-resistant cells compared with wild type MCF7 cells, which also showed limited overlap in their response to tamoxifen and estrogen deprivation in terms of their gene expression (10) and DNA methylation profiles (18).

In our analyses we adjusted for clinical variables associated with survival (tumour stage, age and AI treatment (ER+/HER2- cohort only)) in order to estimate the independent effect of methylation on survival. It has been shown that methylation profiles can discriminate between the ER+/HER2- subtypes luminal A and B (36). Moreover, patients with a luminal B tumour have worse prognosis compared to patients with a luminal A tumour (37), which is also the case in our ER+/HER2- cohort (HR 2.04, 95%CI 1.11-3.74, p=0.020). We, therefore, also performed survival analyses adjusted for luminal status in addition to the clinical variables mentioned earlier. The single-locus signatures with correction for luminal status showed a considerable overlap of 80%, 44%, and 100% with the original (that is, without correction for luminal status) ER+/HER2-, TAM and AI single-locus signatures respectively (Additional File 11). Notably, all except two CpGs in the ER+/HER2- signature included in the original single-locus signatures still have an FDR<0.1 after correction for luminal status. The risk scores of the original multi-locus signatures were also significantly associated with survival after correction for luminal status (Additional File 11). In summary, the methylation signatures we identified are associated with survival independently of luminal status.

We note that although the methylation profiles provided by TCGA are measured in untreated primary tumour samples, treatment regimens after initial diagnosis are heterogeneous. Some patients received adjuvant chemotherapy and/or radiotherapy next to endocrine therapy and 42 patients in the TAM and AI cohorts received both types of endocrine treatments. Moreover, the duration of (endocrine) treatment varied among patients. Furthermore, treatment information may not be complete (20). These aspects were not taken into account in our analyses and might have biased the results. We also acknowledge that this study is limited by the relatively modest number of events (i.e., new tumour event, death) for the different cohorts (ER+/HER2-: 97 events in 552 patients; TAM: 24 events in 172 patients; AI: 32 events in 210 patients) due to the relatively short follow-up time. This affects statistical power to identify methylation loci associated with survival.

In this study we assumed that the methylation events in the primary tumour, rather than acquired methylation during tumour progression, are associated with patient survival as a proxy for development of therapy resistance. To validate our results we aimed to use methylation profiles from the International Cancer Genome Consortium (ICGC; https://icgc.org). However, the number of patients in the ICGC breast cancer cohort with reliable information on endocrine treatment was too small to make such a comparison meaningful. Instead, we used DNA methylation measurements obtained from T47D cells as a model system for resistance acquisition in ER+ luminal A breast cancer. We showed that our single-locus signatures for the ER+/HER2- and TAM cohorts were conserved among the loci that are differentially methylated in endocrine-resistant T47D cells. Similarly, our multi-locus signatures for the ER+/HER2- and AI cohorts were also significantly enriched in the T47D experiment. Although this is not a final validation of our results, it strongly suggests that the loci we identified from primary tumours, that is prior to any endocrine treatment, are also associated with endocrine resistance.

Stone et al. (18) recently demonstrated in a small cohort of patients who received endocrine treatment for at least five years that methylation levels in selected ESR1-enhancer loci were significantly increased in primary tumours of patients who relapsed within six years as compared to patients with 14-year relapse free survival. Moreover, these differences were even more pronounced in matched local relapse samples. DNA methylation data measured in a large number of pre- and post-treatment samples obtained from patients who received endocrine therapy that either relapsed due to endocrine therapy resistance or remained relapse-free will enable validation of the signatures identified in this and other studies. Moreover, such a cohort enables comparison of methylation levels in paired primary and local relapse samples providing the opportunity to identify epigenetic drivers of endocrine therapy resistance (38).

## Supporting information

Supplemental Table 1 _ Drugs

Supplemental table 2 _ Subtypes

Supplemental table 3 _ sample annotations

Supplemental table 4 _ single locus

Supplemental file 5 _ KM plots

Supplemental file 6 _ overlap

Supplemental file 7 _ multi locus signature

Supplemental file 8 _ risk score

Supplemental file 9 _ Venn Diagrams

Supplemental file 10 _ stability

Supplemental file 11 _ luminal

## Declarations

### Ethics approval and consent to participate

Not applicable.

### Consent for publication

Not applicable.

### Availability of data and materials

All data used in this study are publicly available on the Genomics Data Commons Legacy Archive (https://portal.gdc.cancer.gov/legacy-archive). Data supporting the findings described in this manuscript are included in the Additional Files.

### Authors’ contributions

MS performed the data analysis, interpreted the results, and drafted the manuscript. SB and ES performed the cell line experiment under supervision of SW. SB, ES, PJV and SW critically reviewed the manuscript. MS, PJV, SW, PDM and AHCvK interpreted the results. PDM and AHCvK conceived the study, supervised the project and helped draft the manuscript. All authors read and approved the final manuscript.

## Acknowledgements

We acknowledge Dr. Alex Michie and Dr. Age K. Smilde for their helpful suggestions while carrying out this research. The results shown here are in part based upon data generated by the TCGA Research Network: https://www.cancer.gov/tcga.

## List of abbreviations

AI: aromatase inhibitor
BRCA: breast invasive carcinoma
CI: confidence interval
ESR1: estrogen receptor 1
ER: estrogen receptor
FDR: false discovery rate
H3K27ac: acetylation of lysine 27 on histone 3
HR: hazard ratio
LTED: long-term estrogen deprived
SE: standard error
TAM: tamoxifen (patient data)
TCGA: The Cancer Genome Atlas
TMX: tamoxifen (cell line experiment)

## Additional files

### Additional file 1: Mapping to generic drug names

Overview of synonyms and spelling variants for drug names used in TCGA BRCA and their mapping to a generic drug name used in our study.

### Additional file 2: Molecular subtypes

Overview of the molecular subtype frequency as determined by immunohistochemistry of ER and HER2 and as predicted by the SCMOD2 model (R package genefu) using TGCA BRCA primary tumour gene expression data. Subtypes are listed for the 1,095 patients for whom gene expression data is available (Figure 1).

### Additional file 3: Sample annotation

Sample annotation for the 552 patients in the ER+/HER2- cohort. The first sheet provides a short definition of the variables included in the second sheet.

### Additional file 4: Single-locus survival analysis

Results of single-locus survival analysis on ER+/HER2-, TAM and AI cohorts.

### Additional file 5: Single-locus Kaplan-Meier plots

Kaplan-Meier plots for each CpG site from the single-locus signatures. Patients were stratified based on the methylation levels of the indicated locus in ER+/HER2, TAM and AI cohorts.

H, methylation level above median; L, methylation level below median. Shaded areas denote the 95% confidence interval in the H and L strata. P-values are based on a log-rank test.

### Additional file 6: Single-locus Venn diagram

Venn diagram of the single-locus signatures in the ER+/HER2-, TAM and AI cohorts.

### Additional file 7: Multi-locus survival analysis

Results of multi-locus survival analysis on ER+/HER2-, TAM and AI cohorts.

### Additional file 8: Survival analysis using risk score

Results of survival analysis of the multi-locus signature using the risk score corrected for selected clinical variables in ER+/HER2-, TAM and AI cohorts.

### Additional file 9: Overlap between single-locus and multi-locus signatures

Venn diagrams of the overlap between single-locus and multi-locus signatures in the three cohorts ER+/HER2-, TAM and AI.

### Additional file 10: Stability of multi-locus signatures

Results of Fisher’s exact test to determine the significance of the overlap between the original multi-locus signature and the stability signature.

### Additional file 11: Survival analyses including luminal status

Reanalysis when also including in luminal status in the (i) multivariable survival analysis, (ii) single-locus survival analysis, and (iii) the risk score for the multi-locus signature.

## References

1. Ferlay J, Colombet M, Soerjomataram I, Dyba T, Randi G, Bettio M, et al. Cancer incidence and mortality patterns in Europe: Estimates for 40 countries and 25 major cancers in 2018. Eur J Cancer. 2018;103:356–87.

2. Johnston SJ, Cheung KL. Endocrine Therapy for Breast Cancer: A Model of Hormonal Manipulation. Oncol Ther. 2018;6(2):141–56.

3. Early Breast Cancer Trialists’ Collaborative G. Effects of chemotherapy and hormonal therapy for early breast cancer on recurrence and 15-year survival: an overview of the randomised trials. Lancet. 2005;365(9472):1687–717.

4. Pouliot MC, Labrie Y, Diorio C, Durocher F. The Role of Methylation in Breast Cancer Susceptibility and Treatment. Anticancer Res. 2015;35(9):4569–74.

5. Abdel-Hafiz H. Epigenetic Mechanisms of Tamoxifen Resistance in Luminal Breast Cancer. Diseases. 2017;5(3):16.

6. Clarke R, Tyson JJ, Dixon JM. Endocrine resistance in breast cancer--An overview and update. Mol Cell Endocrinol. 2015;418 Pt 3:220–34.

7. Bianco S, Gevry N. Endocrine resistance in breast cancer: from cellular signaling pathways to epigenetic mechanisms. Transcription. 2012;3(4):165–70.

8. O’Sullivan DE, Johnson KC, Skinner L, Koestler DC, Christensen BC. Epigenetic and genetic burden measures are associated with tumor characteristics in invasive breast carcinoma. Epigenetics. 2016;11(5):344–53.

9. Hervouet E, Cartron PF, Jouvenot M, Delage-Mourroux R. Epigenetic regulation of estrogen signaling in breast cancer. Epigenetics. 2013;8(3):237–45.

10. Nguyen VTM, Barozzi I, Faronato M, Lombardo Y, Steel JH, Patel N, et al. Differential epigenetic reprogramming in response to specific endocrine therapies promotes cholesterol biosynthesis and cellular invasion. Nat Commun. 2015;6:10044.

11. Fleischer T, Tekpli X, Mathelier A, Wang S, Nebdal D, Dhakal HP, et al. DNA methylation at enhancers identifies distinct breast cancer lineages. Nat Commun. 2017;8(1):1379.

12. Pathiraja TN, Nayak SR, Xi Y, Jiang S, Garee JP, Edwards DP, et al. Epigenetic reprogramming of HOXC10 in endocrine-resistant breast cancer. Sci Transl Med. 2014;6(229):229ra41.

13. Williams KE, Anderton DL, Lee MP, Pentecost BT, Arcaro KF. High-density array analysis of DNA methylation in Tamoxifen-resistant breast cancer cell lines. Epigenetics. 2014;9(2):297–307.

14. Gnyszka A, Jastrzebski Z, Flis S. DNA Methyltransferase Inhibitors and Their Emerging Role in Epigenetic Therapy of Cancer. Anticancer Research. 2013;33(8):2989–96.

15. Lin X, Li J, Yin G, Zhao Q, Elias D, Lykkesfeldt AE, et al. Integrative analyses of gene expression and DNA methylation profiles in breast cancer cell line models of tamoxifen-resistance indicate a potential role of cells with stem-like properties. Breast Cancer Res. 2013;15(6):R119.

16. Gyorffy B, Bottai G, Fleischer T, Munkacsy G, Budczies J, Paladini L, et al. Aberrant DNA methylation impacts gene expression and prognosis in breast cancer subtypes. Int J Cancer. 2016;138(1):87–97.

17. Fleischer T, Frigessi A, Johnson KC, Edvardsen H, Touleimat N, Klajic J, et al. Genome-wide DNA methylation profiles in progression to in situ and invasive carcinoma of the breast with impact on gene transcription and prognosis. Genome Biol. 2014;15(8):435.

18. Stone A, Zotenko E, Locke WJ, Korbie D, Millar EK, Pidsley R, et al. DNA methylation of oestrogen-regulated enhancers defines endocrine sensitivity in breast cancer. Nat Commun. 2015;6:7758.

19. Cancer Genome Atlas N. Comprehensive molecular portraits of human breast tumours. Nature. 2012;490(7418):61–70.

20. Liu J, Lichtenberg T, Hoadley KA, Poisson LM, Lazar AJ, Cherniack AD, et al. An Integrated TCGA Pan-Cancer Clinical Data Resource to Drive High-Quality Survival Outcome Analytics. Cell. 2018;173(2):400–16 e11.

21. Colaprico A, Silva TC, Olsen C, Garofano L, Cava C, Garolini D, et al. TCGAbiolinks: an R/Bioconductor package for integrative analysis of TCGA data. Nucleic Acids Res. 2016;44(8):e71.

22. Robinson MD, McCarthy DJ, Smyth GK. edgeR: a Bioconductor package for differential expression analysis of digital gene expression data. Bioinformatics. 2010;26(1):139–40.

23. Gendoo DM, Ratanasirigulchai N, Schroder MS, Pare L, Parker JS, Prat A, et al. Genefu: an R/Bioconductor package for computation of gene expression-based signatures in breast cancer. Bioinformatics. 2016;32(7):1097–9.

24. Aryee MJ, Jaffe AE, Corrada-Bravo H, Ladd-Acosta C, Feinberg AP, Hansen KD, et al. Minfi: a flexible and comprehensive Bioconductor package for the analysis of Infinium DNA methylation microarrays. Bioinformatics. 2014;30(10):1363–9.

25. Fortin JP, Labbe A, Lemire M, Zanke BW, Hudson TJ, Fertig EJ, et al. Functional normalization of 450k methylation array data improves replication in large cancer studies. Genome Biol. 2014;15(12):503.

26. Chen YA, Lemire M, Choufani S, Butcher DT, Grafodatskaya D, Zanke BW, et al. Discovery of cross-reactive probes and polymorphic CpGs in the Illumina Infinium HumanMethylation450 microarray. Epigenetics. 2013;8(2):203–9.

27. Ronneberg JA, Fleischer T, Solvang HK, Nordgard SH, Edvardsen H, Potapenko I, et al. Methylation profiling with a panel of cancer related genes: association with estrogen receptor, TP53 mutation status and expression subtypes in sporadic breast cancer. Mol Oncol. 2011;5(1):61–76.

28. Lyman GH, Kuderer NM, Lyman SL, Debus M, Minton S, Balducci L, et al. Menopausal Status and the Impact of Early Recurrence on Breast Cancer Survival. Cancer Control. 1997;4(4):335–41.

29. Cianfrocca M, Goldstein LJ. Prognostic and predictive factors in early-stage breast cancer. Oncologist. 2004;9(6):606–16.

30. van Buuren S, Groothuis-Oudshoorn K. mice: Multivariate Imputation by Chained Equations in R. Journal of Statistical Software. 2011;45(3):1–67.

31. Simon N, Friedman J, Hastie T, Tibshirani R. Regularization Paths for Cox’s Proportional Hazards Model via Coordinate Descent. J Stat Softw. 2011;39(5):1–13.

32. Martin LA, Ghazoui Z, Weigel MT, Pancholi S, Dunbier A, Johnston S, et al. An in vitro model showing adaptation to long-term oestrogen deprivation highlights the clinical potential for targeting kinase pathways in combination with aromatase inhibition. Steroids. 2011;76(8):772–6.

33. Pidsley R, Zotenko E, Peters TJ, Lawrence MG, Risbridger GP, Molloy P, et al. Critical evaluation of the Illumina MethylationEPIC BeadChip microarray for whole-genome DNA methylation profiling. Genome Biol. 2016;17(1):208.

34. Wu D, Lim E, Vaillant F, Asselin-Labat ML, Visvader JE, Smyth GK. ROAST: rotation gene set tests for complex microarray experiments. Bioinformatics. 2010;26(17):2176–82.

35. Saadatmand S, Bretveld R, Siesling S, Tilanus-Linthorst MMA. Influence of tumour stage at breast cancer detection on survival in modern times: population based study in 173 797 patients. Bmj-Brit Med J. 2015;351.

36. Stefansson OA, Moran S, Gomez A, Sayols S, Arribas-Jorba C, Sandoval J, et al. A DNA methylation-based definition of biologically distinct breast cancer subtypes. Mol Oncol. 2015;9(3):555–68.

37. Fallahpour S, Navaneelan T, De P, Borgo A. Breast cancer survival by molecular subtype: a population-based analysis of cancer registry data. CMAJ Open. 2017;5(3):E734–E9.

38. Chatterjee A, Rodger EJ, Eccles MR. Epigenetic drivers of tumourigenesis and cancer metastasis. Semin Cancer Biol. 2018;51:149–59.

