## Supplemental file 5 _ KM plots for "Candidate methylation sites associated with endocrine therapy resistance in the TCGA ER+/HER2- breast cancer cohort"

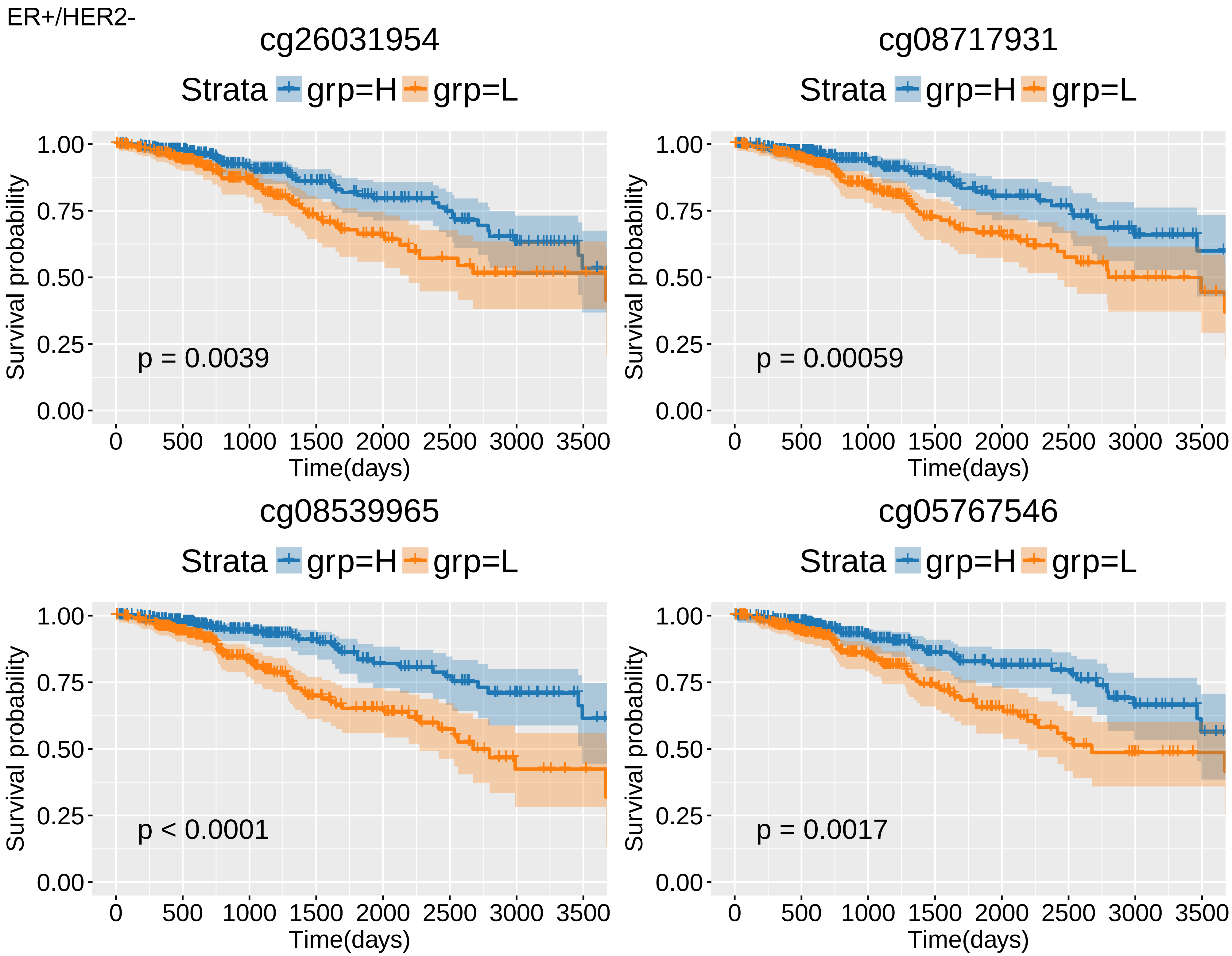

ER+/HER2-
cg26031954
Strata grp=H grp=L
p = 0.0039
cg08717931
Strata grp=H grp=L
p = 0.00059
cg08539965
Strata grp=H grp=L
p < 0.0001
cg05767546
Strata grp=H grp=L
p = 0.0017
Survival probability
1.00
0.75
0.50
0.25
0.00
Time(days)
0
500
1000
1500
2000
2500
3000
3500

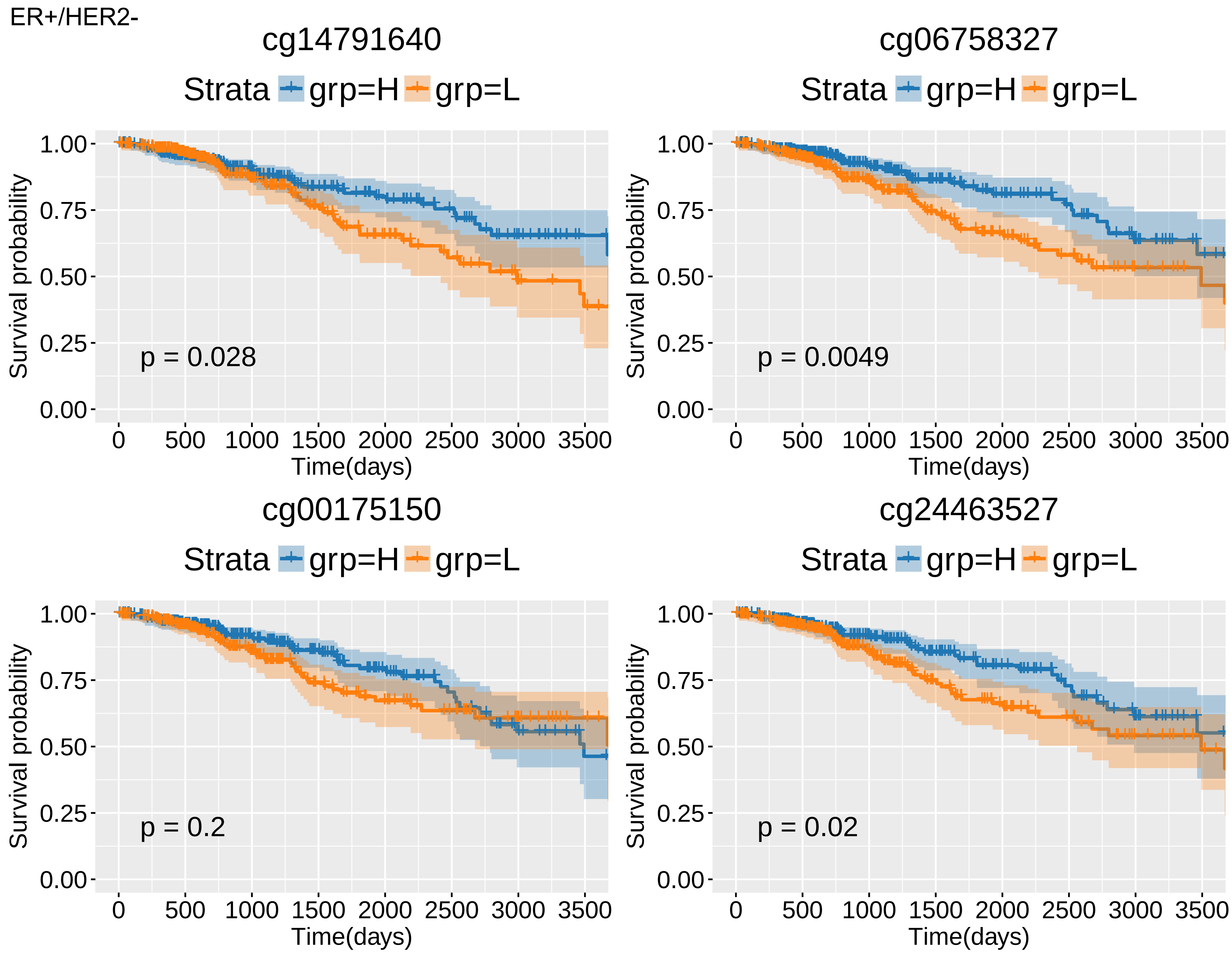
ER+/HER2-
cg14791640
Strata grp=H grp=L
Survival probability
1.00
0.75
0.50
0.25
0.00
p = 0.028
0 500 1000 1500 2000 2500 3000 3500
Time(days)
cg06758327
Strata grp=H grp=L
Survival probability
p = 0.0049
Time(days)
cg00175150
Strata grp=H grp=L
Survival probability
p = 0.2
Time(days)
cg24463527
Strata grp=H grp=L
Survival probability
p = 0.02
Time(days)

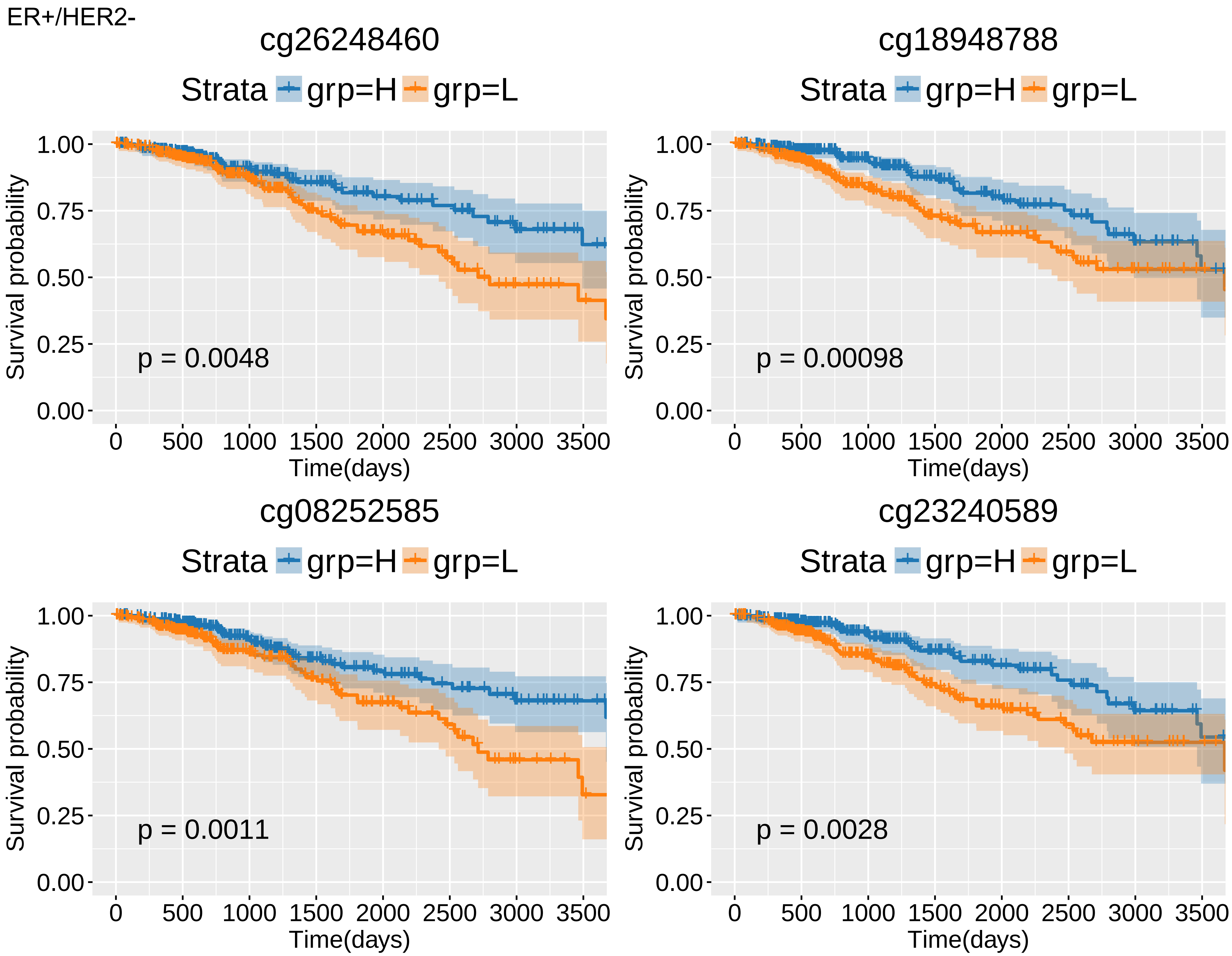

ER+/HER2-
cg26248460
Strata grp=H grp=L
Survival probability
1.00
0.75
0.50
0.25
0.00
p = 0.0048
0 500 1000 1500 2000 2500 3000 3500
Time(days)
cg18948788
Strata grp=H grp=L
Survival probability
p = 0.00098
Time(days)
cg08252585
Strata grp=H grp=L
Survival probability
p = 0.0011
Time(days)
cg23240589
Strata grp=H grp=L
Survival probability
p = 0.0028
Time(days)

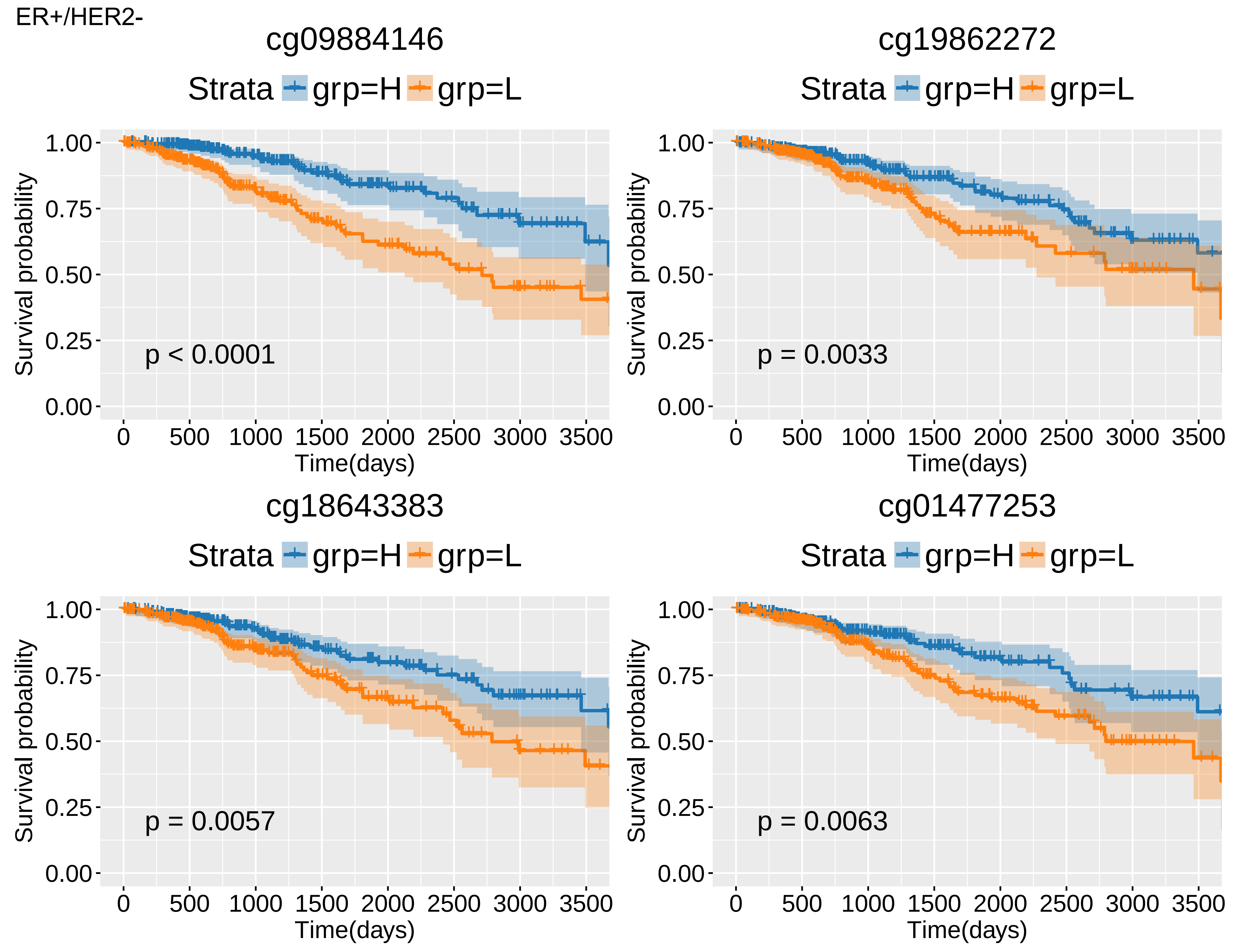
ER+/HER2-
cg09884146
Strata grp=H grp=L
Survival probability
1.00
0.75
0.50
0.25
0.00
p < 0.0001
0 500 1000 1500 2000 2500 3000 3500
Time(days)
cg19862272
Strata grp=H grp=L
Survival probability
p = 0.0033
Time(days)
cg18643383
Strata grp=H grp=L
Survival probability
p = 0.0057
Time(days)
cg01477253
Strata grp=H grp=L
Survival probability
p = 0.0063
Time(days)

ER+/HER2-

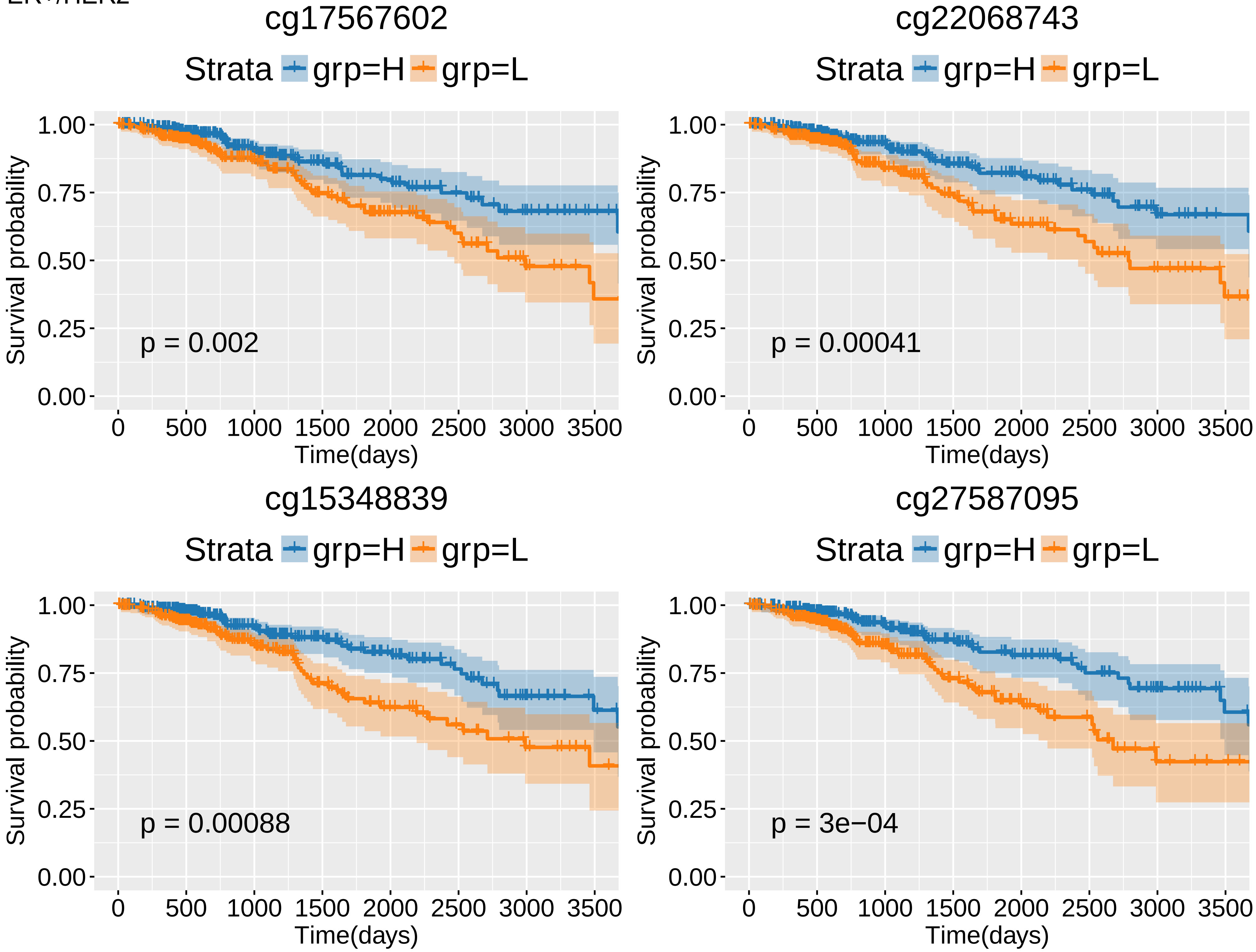
cg17567602
Strata grp=H grp=L
p = 0.002
cg22068743
Strata grp=H grp=L
p = 0.00041
cg15348839
Strata grp=H grp=L
p = 0.00088
cg27587095
Strata grp=H grp=L
p = 3e−04
Survival probability
1.00
0.75
0.50
0.25
0.00
Time(days)
0 500 1000 1500 2000 2500 3000 3500

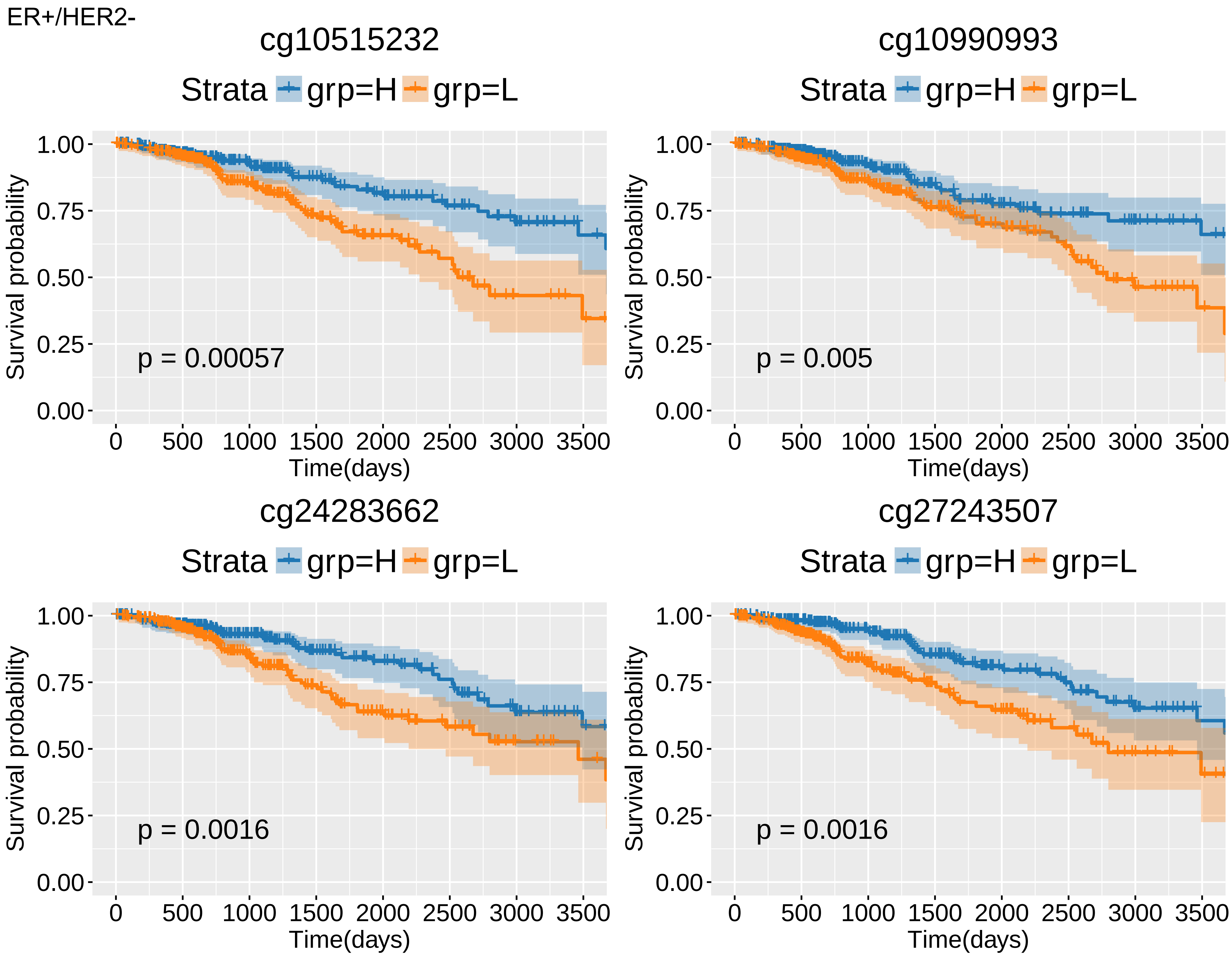
ER+/HER2-
cg10515232
Strata grp=H grp=L
Survival probability
1.00
0.75
0.50
0.25
0.00
p = 0.00057
0 500 1000 1500 2000 2500 3000 3500
Time(days)
cg10990993
Strata grp=H grp=L
Survival probability
p = 0.005
Time(days)
cg24283662
Strata grp=H grp=L
Survival probability
p = 0.0016
Time(days)
cg27243507
Strata grp=H grp=L
Survival probability
p = 0.0016
Time(days)

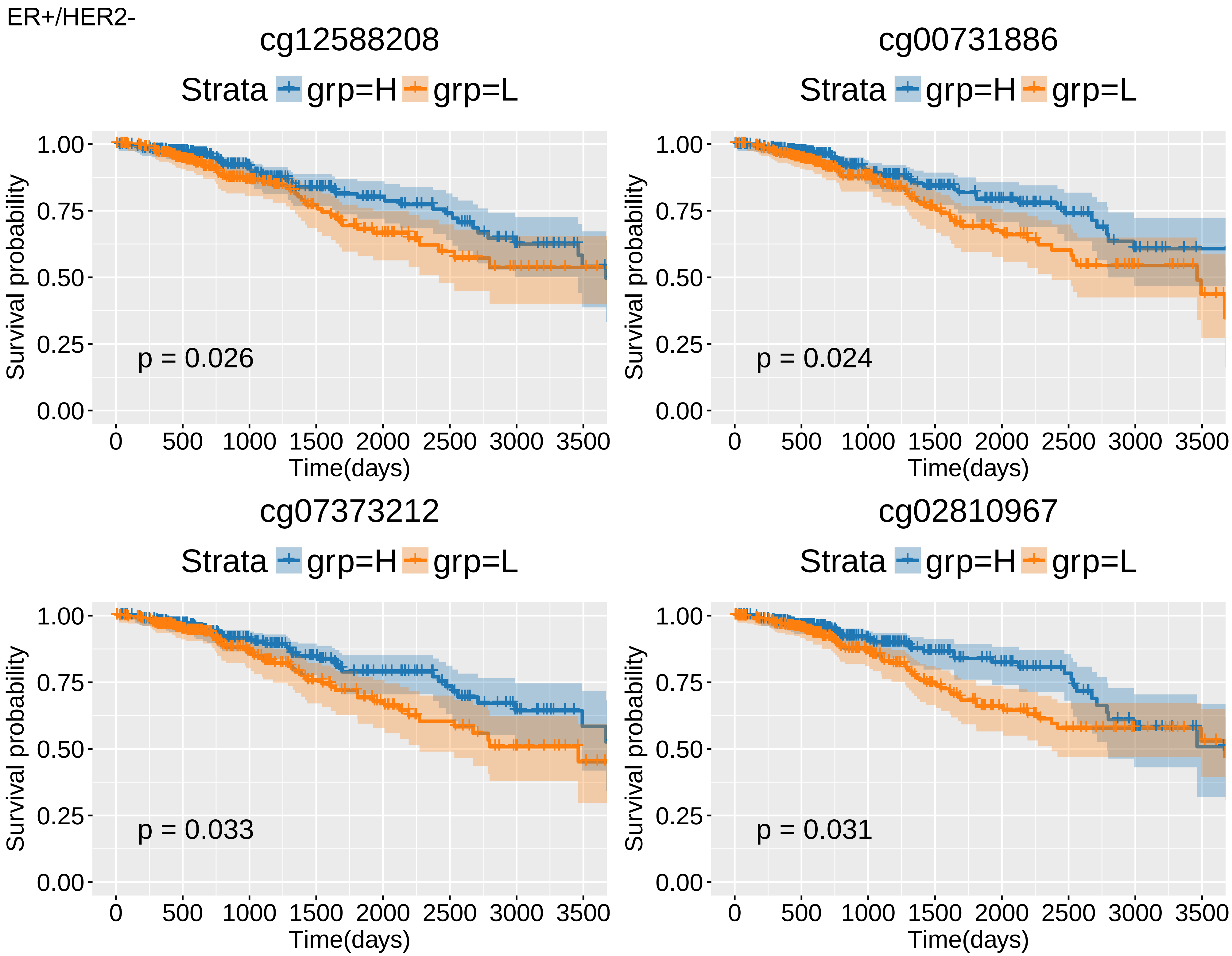
ER+/HER2-
cg12588208
Strata grp=H grp=L
p = 0.026
cg00731886
Strata grp=H grp=L
p = 0.024
cg07373212
Strata grp=H grp=L
p = 0.033
cg02810967
Strata grp=H grp=L
p = 0.031
Survival probability
Time(days)
1.00
0.75
0.50
0.25
0.00
0
500
1000
1500
2000
2500
3000
3500

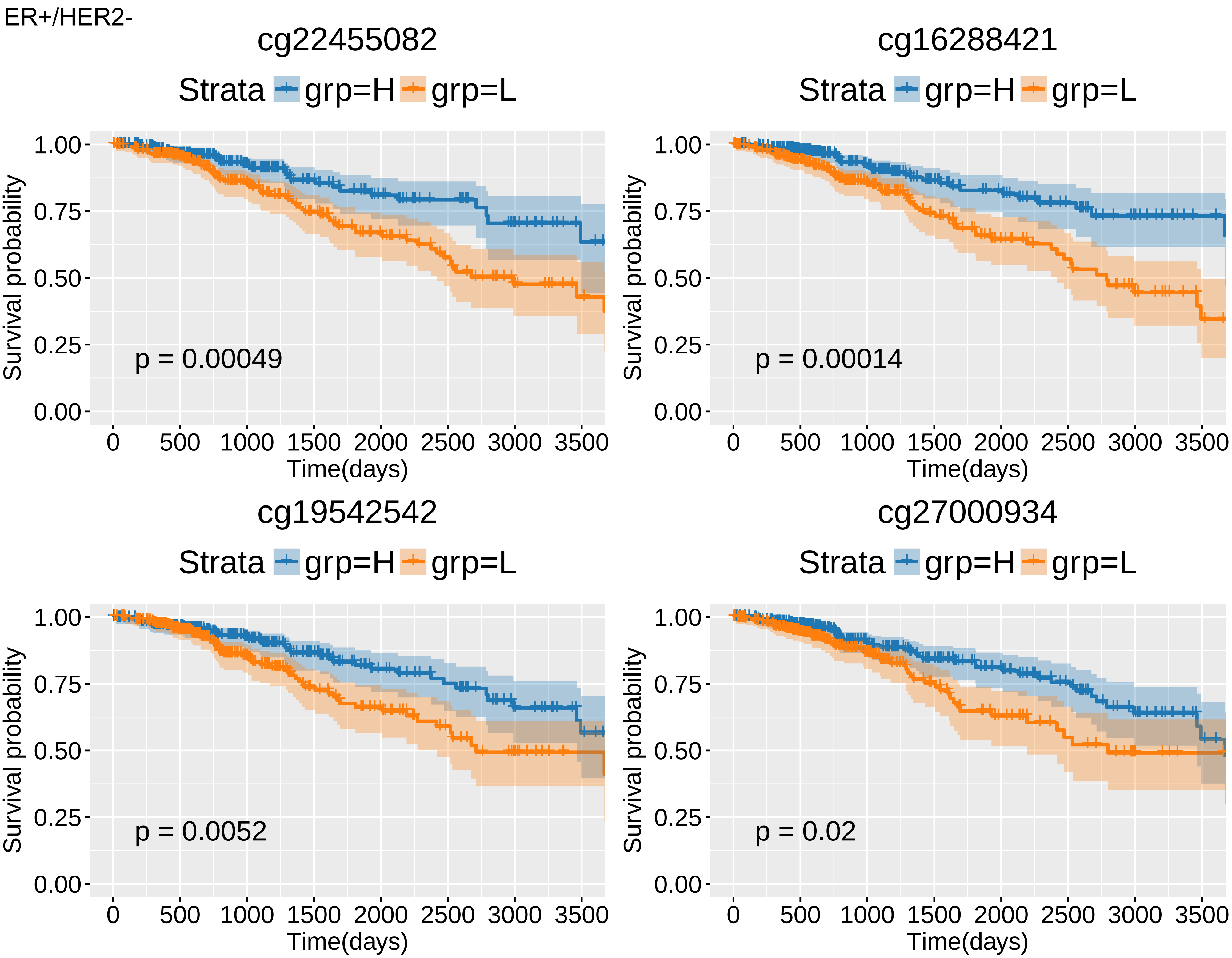
ER+/HER2-
cg22455082
Strata grp=H grp=L
Survival probability
1.00
0.75
0.50
0.25
0.00
p = 0.00049
0 500 1000 1500 2000 2500 3000 3500
Time(days)
cg16288421
Strata grp=H grp=L
Survival probability
p = 0.00014
Time(days)
cg19542542
Strata grp=H grp=L
Survival probability
p = 0.0052
Time(days)
cg27000934
Strata grp=H grp=L
Survival probability
p = 0.02
Time(days)

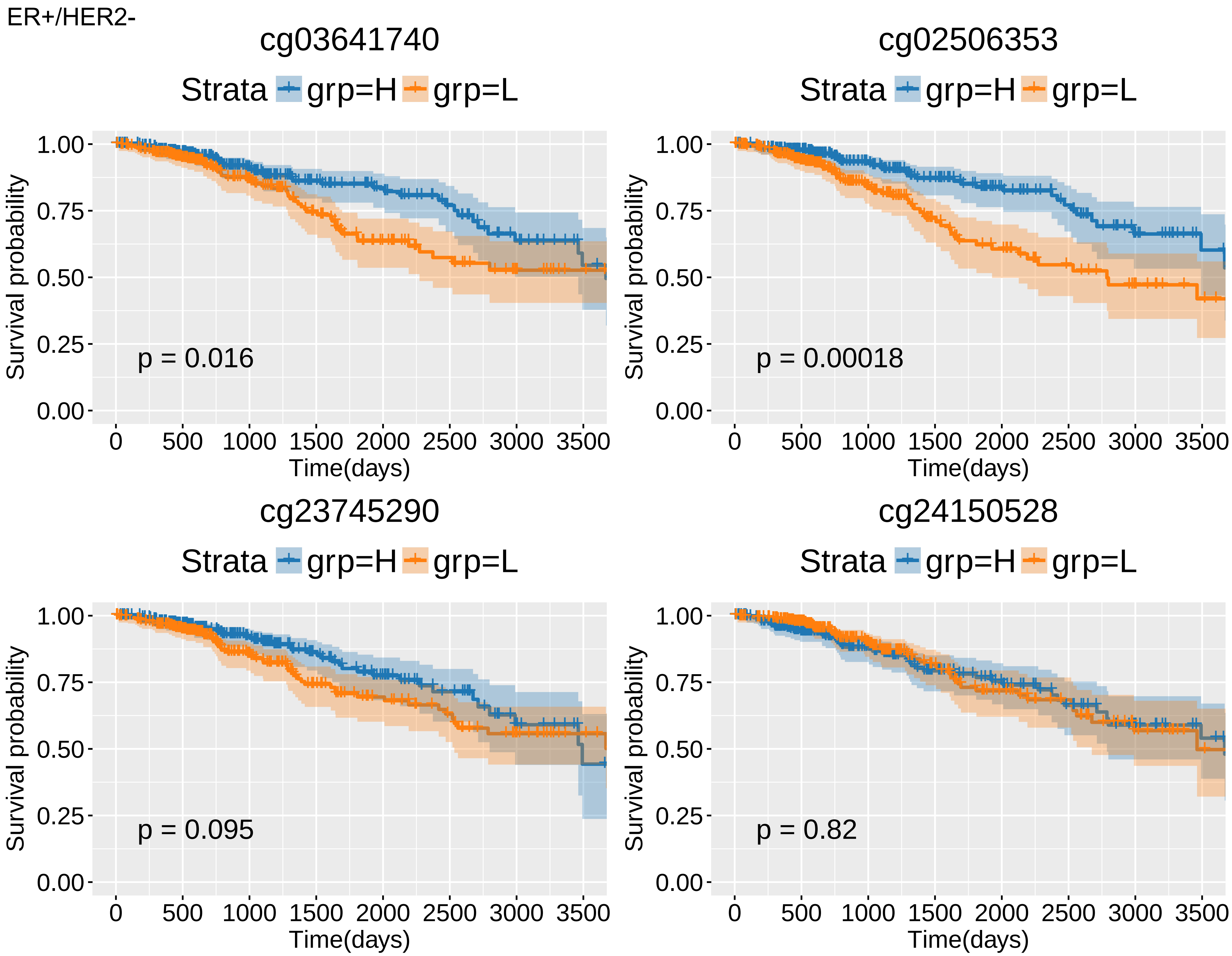
ER+/HER2-
cg03641740
Strata grp=H grp=L
Survival probability
1.00
0.75
0.50
0.25
0.00
0
500
1000
1500
2000
2500
3000
3500
Time(days)
p = 0.016
cg02506353
Strata grp=H grp=L
Survival probability
Time(days)
p = 0.00018
cg23745290
Strata grp=H grp=L
Survival probability
Time(days)
p = 0.095
cg24150528
Strata grp=H grp=L
Survival probability
Time(days)
p = 0.82

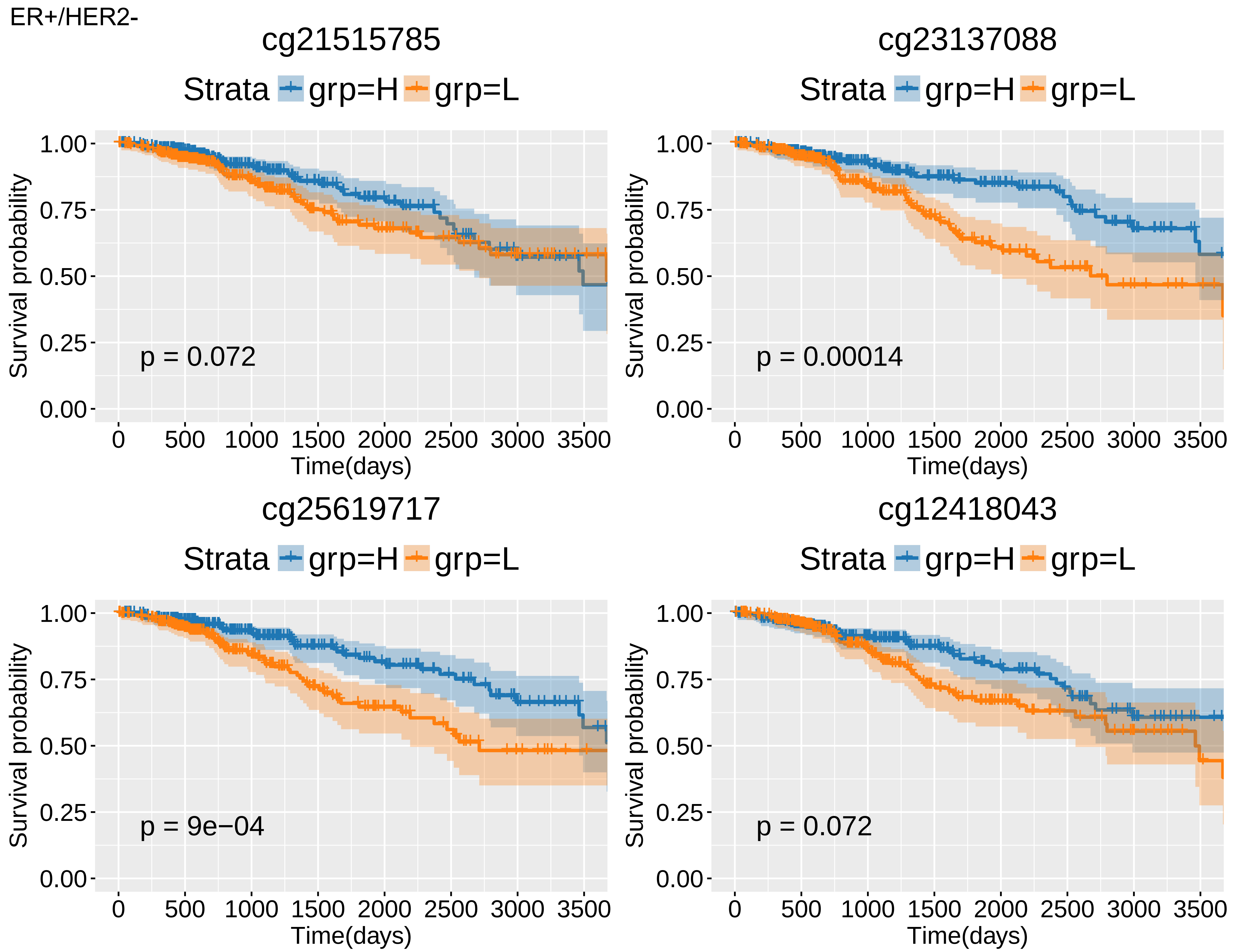
ER+/HER2-
cg21515785
Strata grp=H grp=L
p = 0.072
cg23137088
Strata grp=H grp=L
p = 0.00014
cg25619717
Strata grp=H grp=L
p = 9e−04
cg12418043
Strata grp=H grp=L
p = 0.072
Survival probability
1.00
0.75
0.50
0.25
0.00
0
500
1000
1500
2000
2500
3000
3500
Time(days)

ER+/HER2-

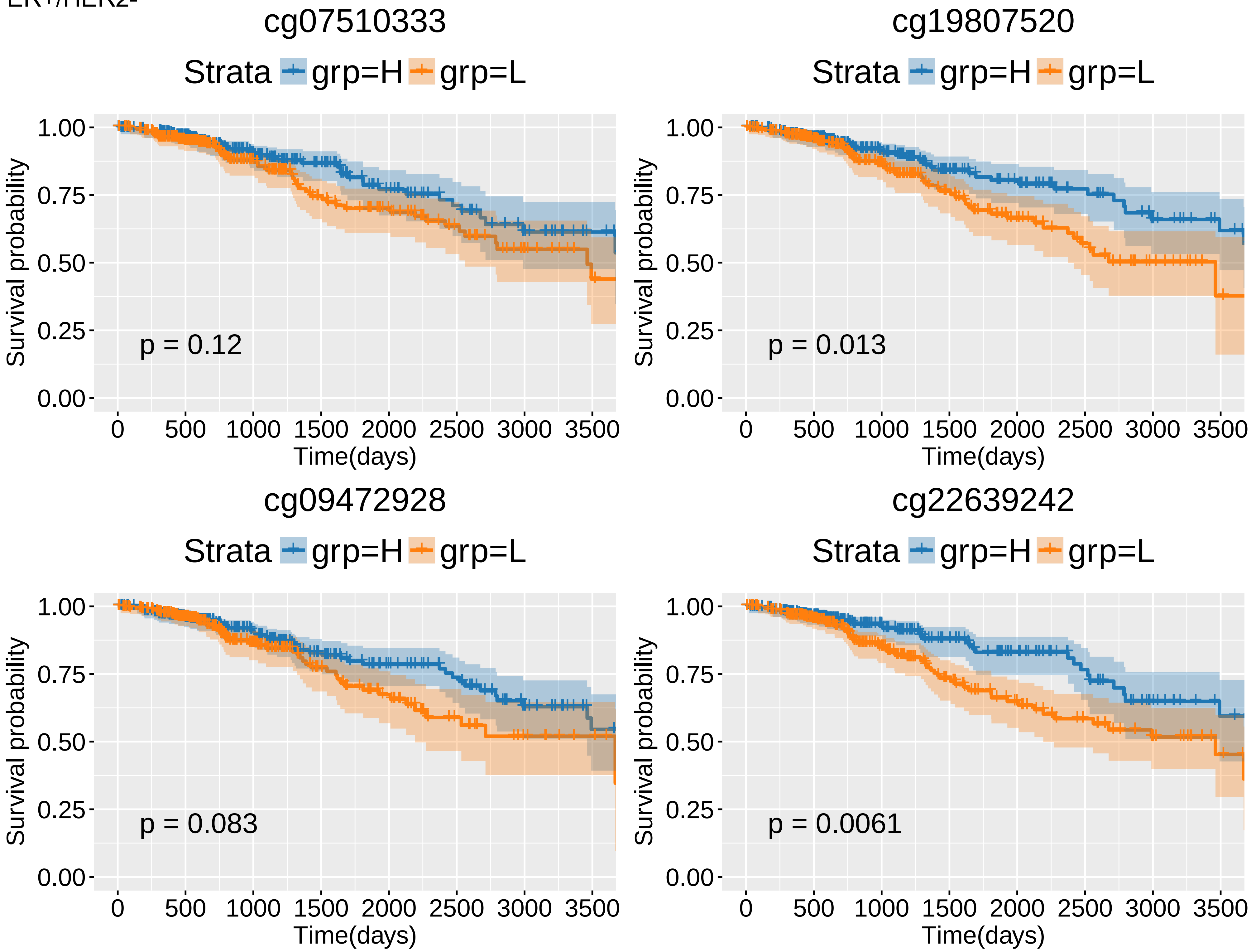

cg07510333
Strata
grp=H
grp=L
Survival probability
1.00
0.75
0.50
0.25
0.00
p = 0.12
0
500
1000
1500
2000
2500
3000
3500
Time(days)
cg19807520
Strata
grp=H
grp=L
Survival probability
p = 0.013
Time(days)
cg09472928
Strata
grp=H
grp=L
Survival probability
p = 0.083
Time(days)
cg22639242
Strata
grp=H
grp=L
Survival probability
p = 0.0061
Time(days)

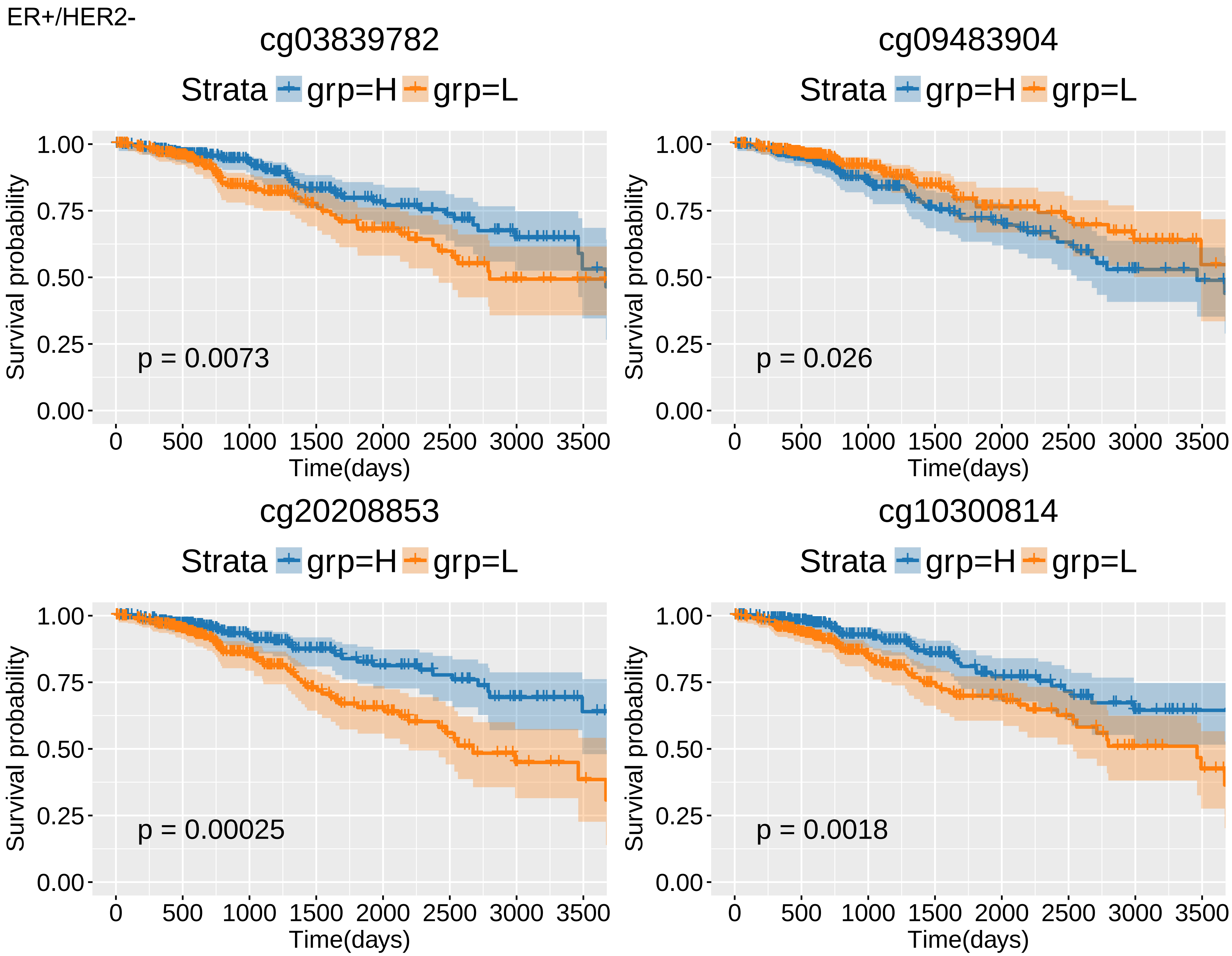
ER+/HER2-
cg03839782
Strata grp=H grp=L
p = 0.0073
cg09483904
Strata grp=H grp=L
p = 0.026
cg20208853
Strata grp=H grp=L
p = 0.00025
cg10300814
Strata grp=H grp=L
p = 0.0018
Survival probability
1.00
0.75
0.50
0.25
0.00
Time(days)
0
500
1000
1500
2000
2500
3000
3500

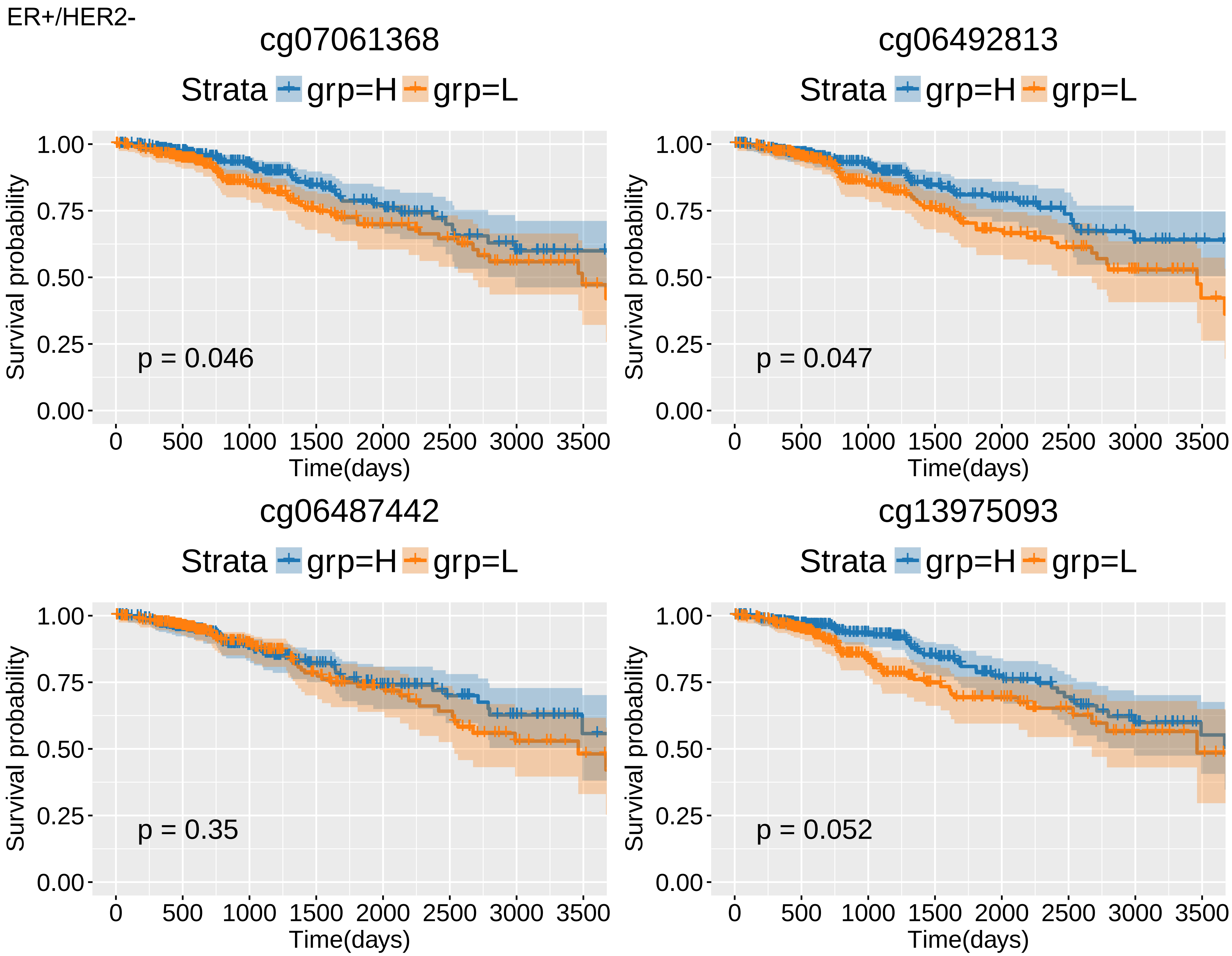
ER+/HER2-
cg07061368
Strata grp=H grp=L
Survival probability
1.00
0.75
0.50
0.25
0.00
p = 0.046
0 500 1000 1500 2000 2500 3000 3500
Time(days)
cg06492813
Strata grp=H grp=L
Survival probability
1.00
0.75
0.50
0.25
0.00
p = 0.047
0 500 1000 1500 2000 2500 3000 3500
Time(days)
cg06487442
Strata grp=H grp=L
Survival probability
1.00
0.75
0.50
0.25
0.00
p = 0.35
0 500 1000 1500 2000 2500 3000 3500
Time(days)
cg13975093
Strata grp=H grp=L
Survival probability
1.00
0.75
0.50
0.25
0.00
p = 0.052
0 500 1000 1500 2000 2500 3000 3500
Time(days)

ER+/HER2-

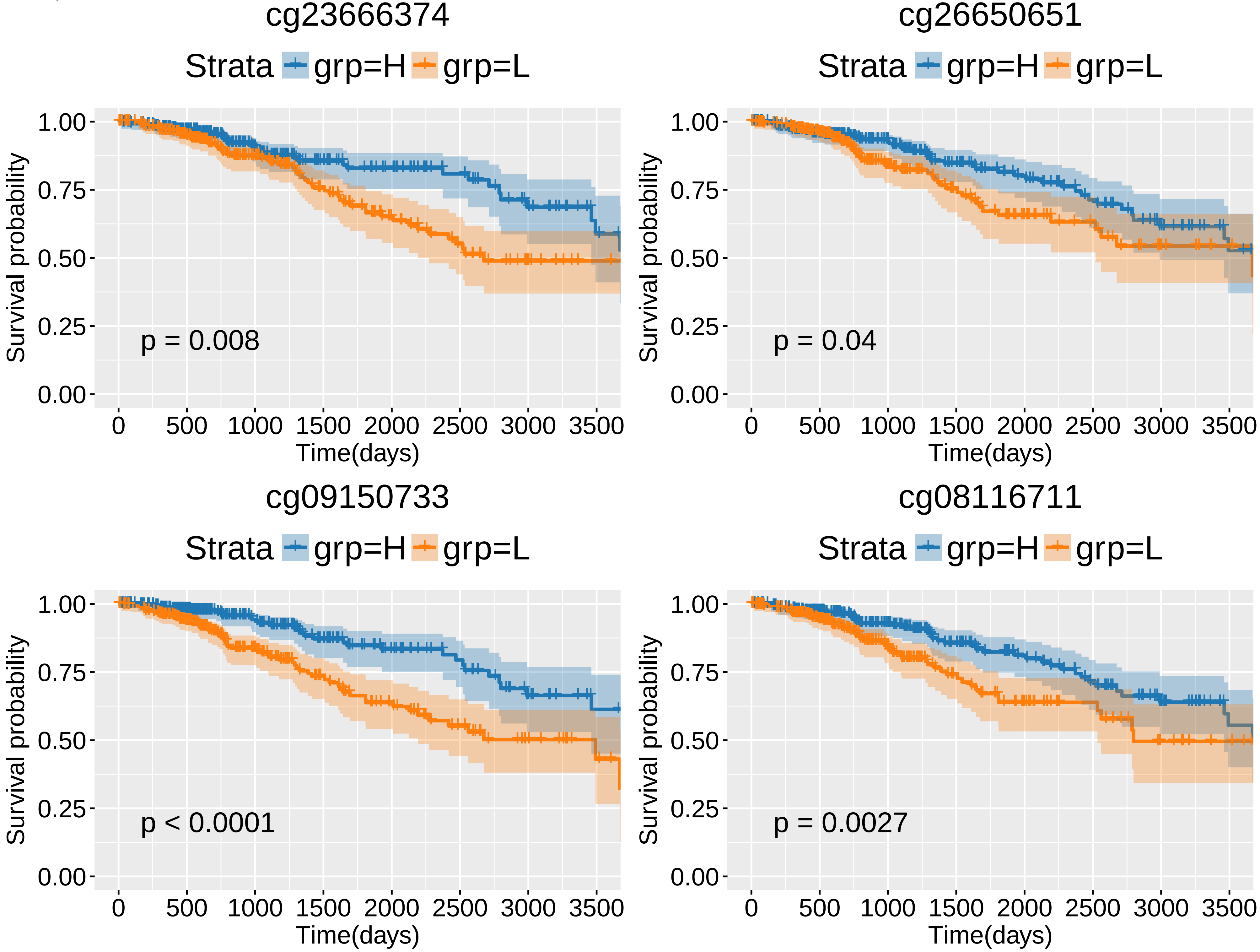
cg23666374
Strata grp=H grp=L
Survival probability
1.00
0.75
0.50
0.25
0.00
p = 0.008
0 500 1000 1500 2000 2500 3000 3500
Time(days)
cg26650651
Strata grp=H grp=L
Survival probability
1.00
0.75
0.50
0.25
0.00
p = 0.04
0 500 1000 1500 2000 2500 3000 3500
Time(days)
cg09150733
Strata grp=H grp=L
Survival probability
1.00
0.75
0.50
0.25
0.00
p < 0.0001
0 500 1000 1500 2000 2500 3000 3500
Time(days)
cg08116711
Strata grp=H grp=L
Survival probability
1.00
0.75
0.50
0.25
0.00
p = 0.0027
0 500 1000 1500 2000 2500 3000 3500
Time(days)

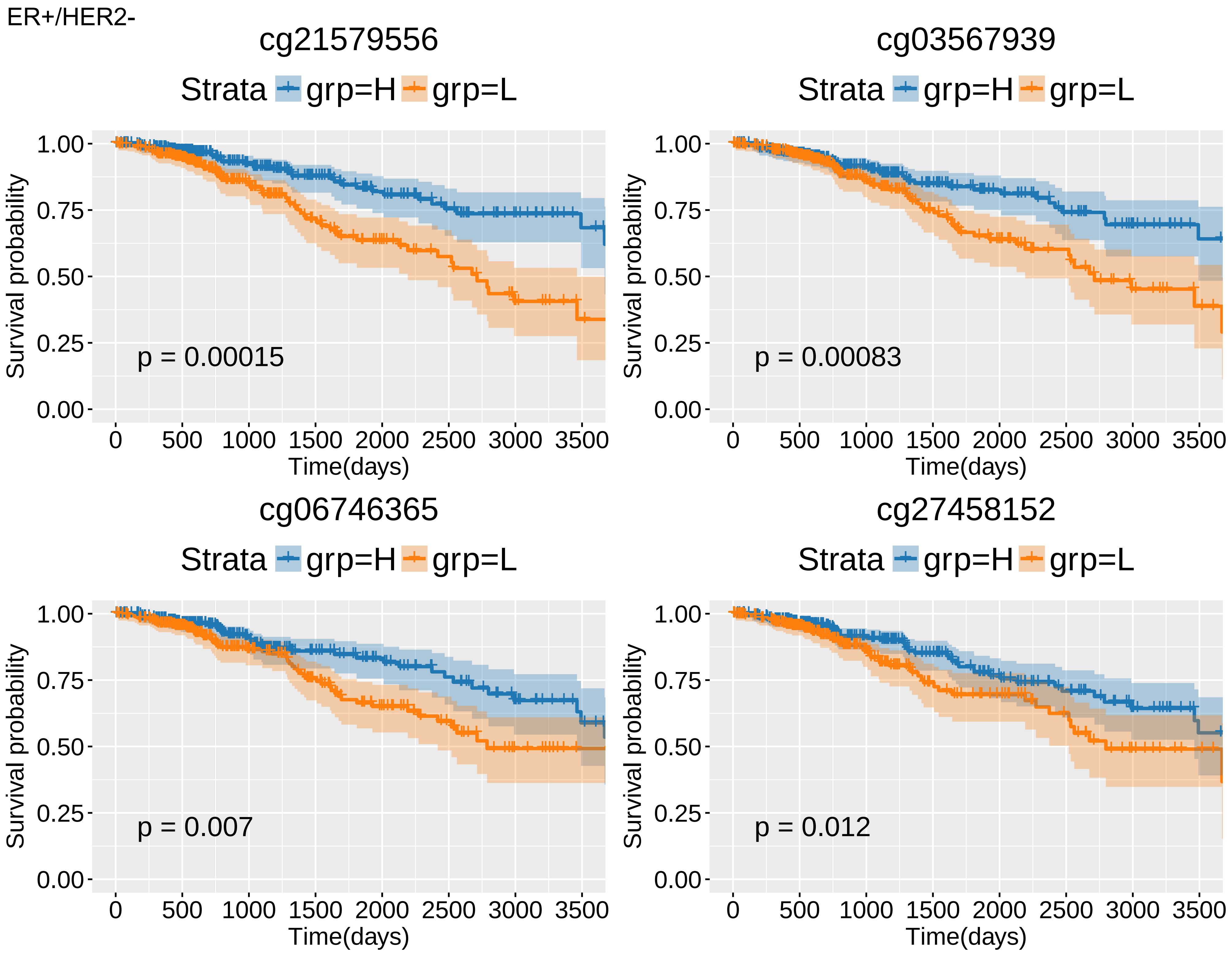
ER+/HER2-
cg21579556
Strata grp=H grp=L
p = 0.00015
cg03567939
Strata grp=H grp=L
p = 0.00083
cg06746365
Strata grp=H grp=L
p = 0.007
cg27458152
Strata grp=H grp=L
p = 0.012
Survival probability
1.00
0.75
0.50
0.25
0.00
Time(days)
0
500
1000
1500
2000
2500
3000
3500

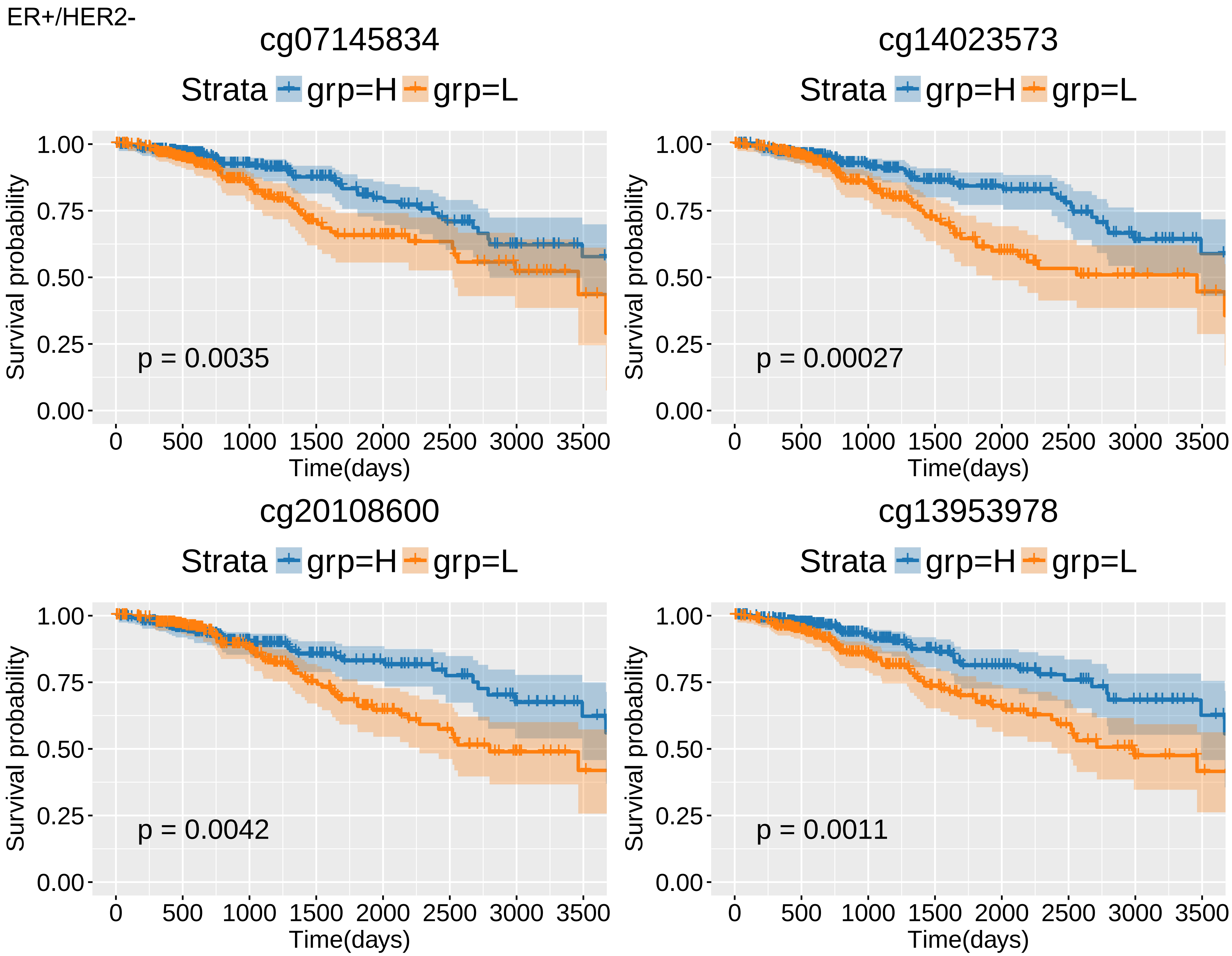
ER+/HER2-
cg07145834
Strata grp=H grp=L
p = 0.0035
Survival probability
Time(days)
cg14023573
Strata grp=H grp=L
p = 0.00027
Survival probability
Time(days)
cg20108600
Strata grp=H grp=L
p = 0.0042
Survival probability
Time(days)
cg13953978
Strata grp=H grp=L
p = 0.0011
Survival probability
Time(days)

ER+/HER2-

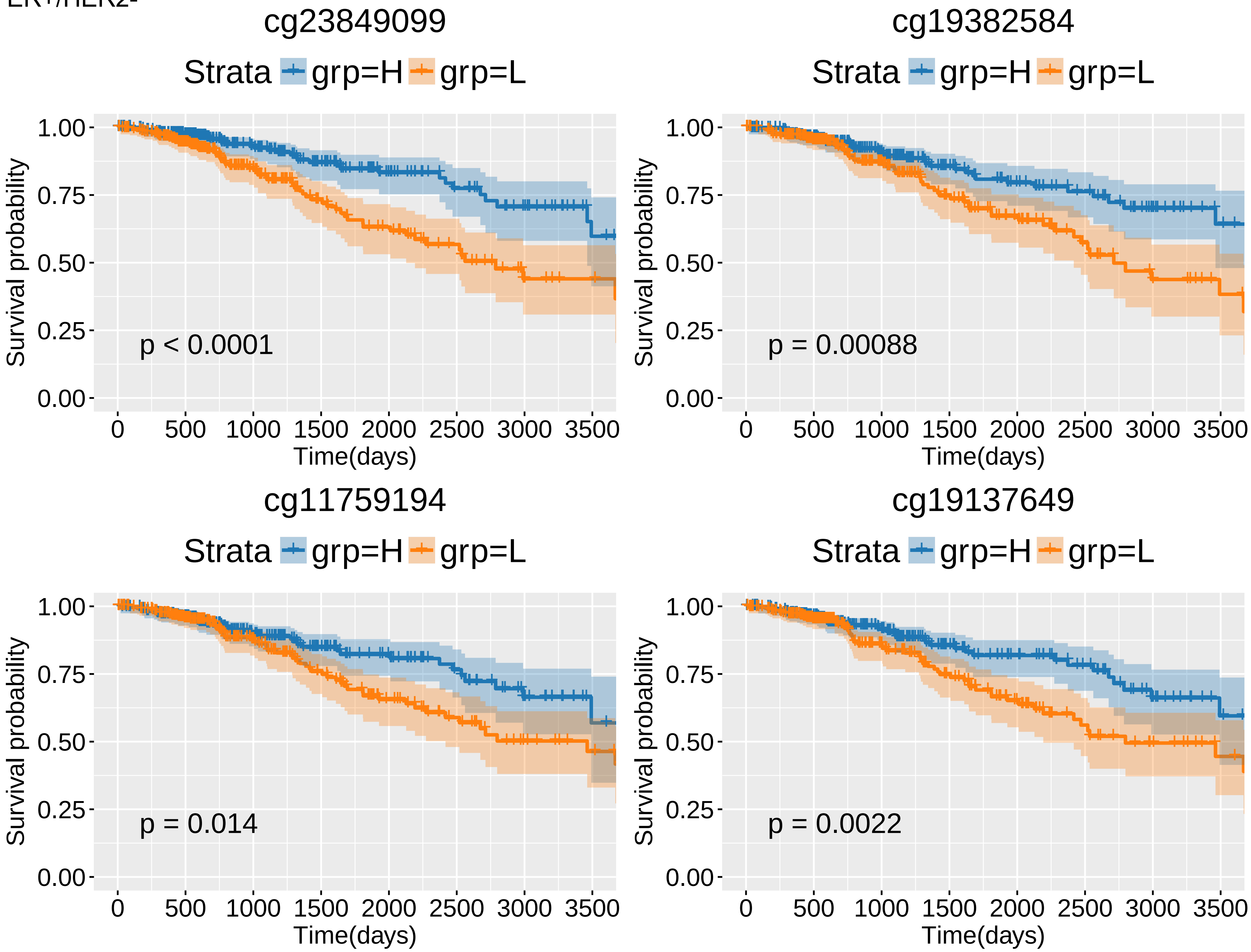
cg23849099
Strata grp=H grp=L
Survival probability
1.00
0.75
0.50
0.25
0.00
p < 0.0001
0 500 1000 1500 2000 2500 3000 3500
Time(days)
cg19382584
Strata grp=H grp=L
Survival probability
p = 0.00088
Time(days)
cg11759194
Strata grp=H grp=L
Survival probability
p = 0.014
Time(days)
cg19137649
Strata grp=H grp=L
Survival probability
p = 0.0022
Time(days)

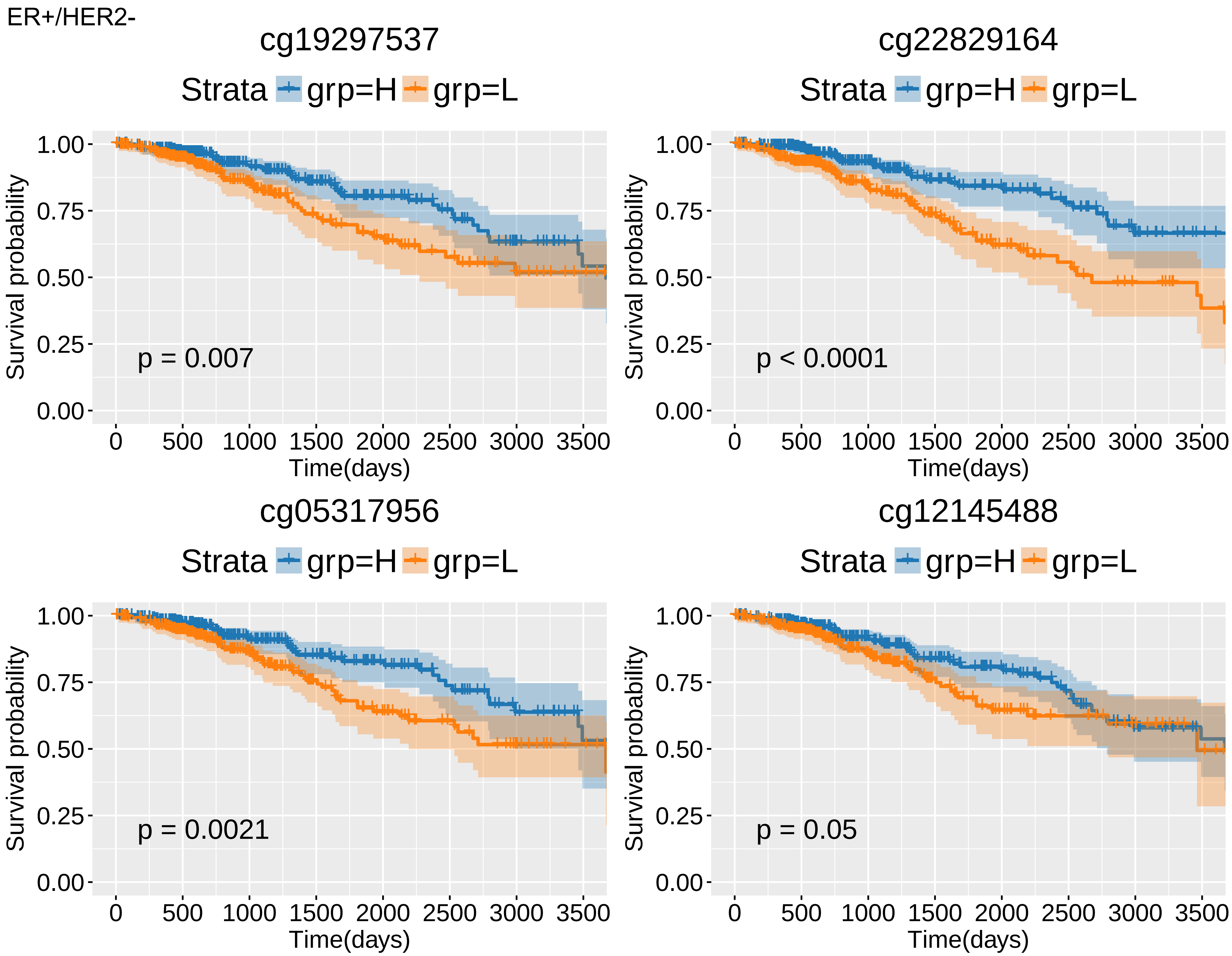
ER+/HER2-
cg19297537
Strata grp=H grp=L
Survival probability
1.00
0.75
0.50
0.25
0.00
p = 0.007
0 500 1000 1500 2000 2500 3000 3500
Time(days)
cg22829164
Strata grp=H grp=L
Survival probability
p < 0.0001
Time(days)
cg05317956
Strata grp=H grp=L
Survival probability
p = 0.0021
Time(days)
cg12145488
Strata grp=H grp=L
Survival probability
p = 0.05
Time(days)

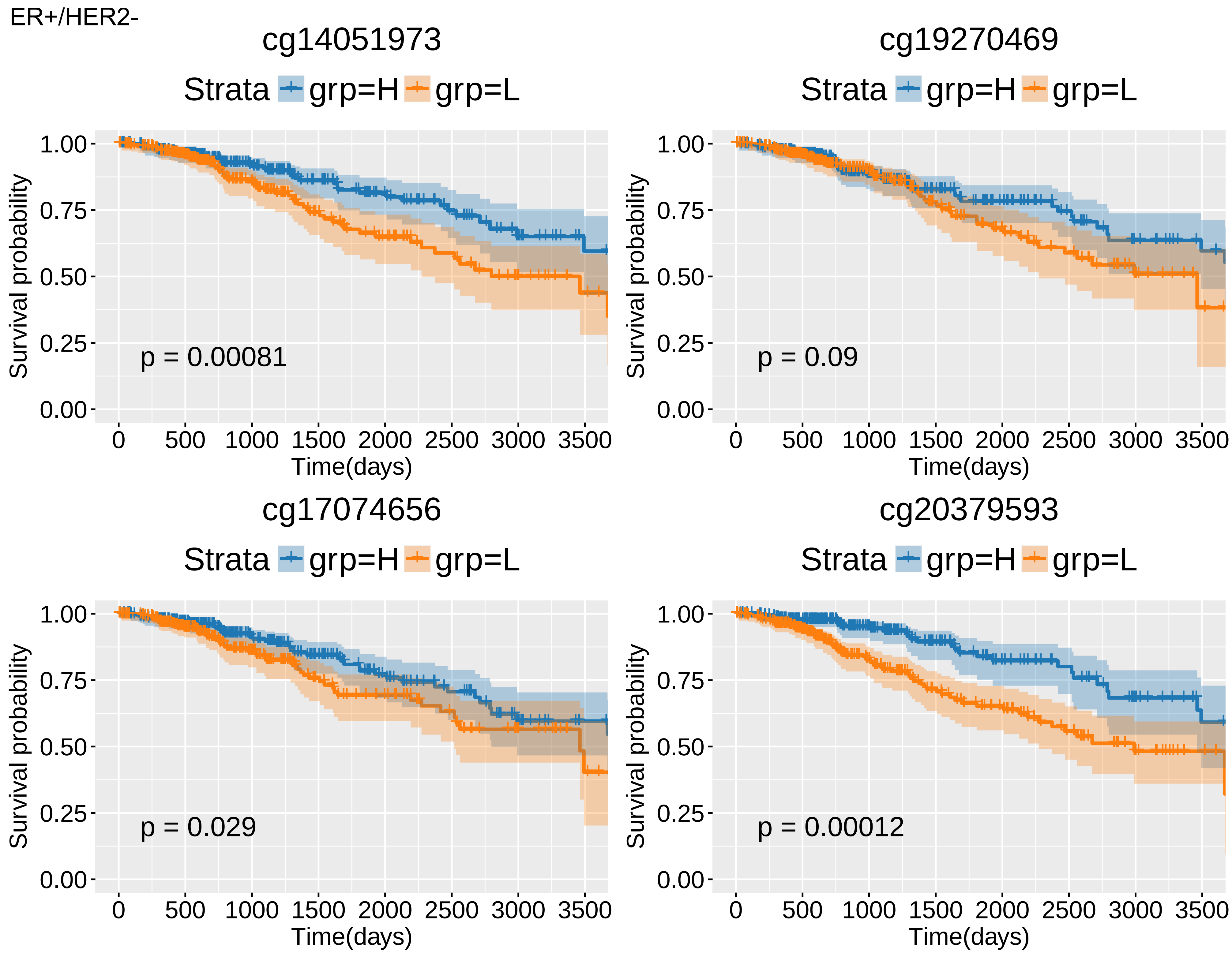

ER+/HER2-
cg14051973
Strata grp=H grp=L
Survival probability
1.00
0.75
0.50
0.25
0.00
p = 0.00081
0 500 1000 1500 2000 2500 3000 3500
Time(days)
cg19270469
Strata grp=H grp=L
Survival probability
p = 0.09
Time(days)
cg17074656
Strata grp=H grp=L
Survival probability
p = 0.029
Time(days)
cg20379593
Strata grp=H grp=L
Survival probability
p = 0.00012
Time(days)

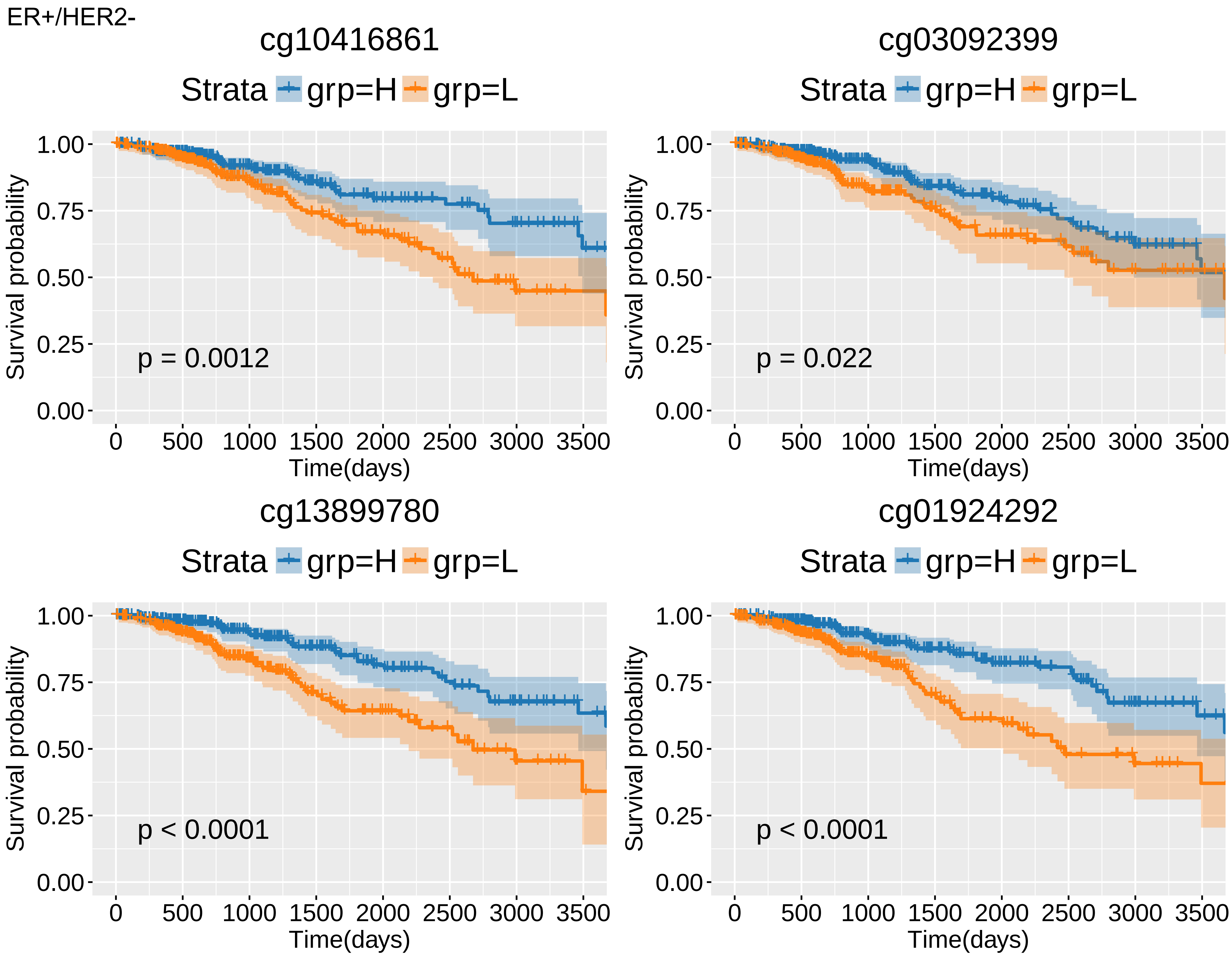
ER+/HER2-
cg10416861
Strata grp=H grp=L
Survival probability
1.00
0.75
0.50
0.25
0.00
p = 0.0012
0 500 1000 1500 2000 2500 3000 3500
Time(days)
cg03092399
Strata grp=H grp=L
Survival probability
p = 0.022
Time(days)
cg13899780
Strata grp=H grp=L
Survival probability
p < 0.0001
Time(days)
cg01924292
Strata grp=H grp=L
Survival probability
p < 0.0001
Time(days)

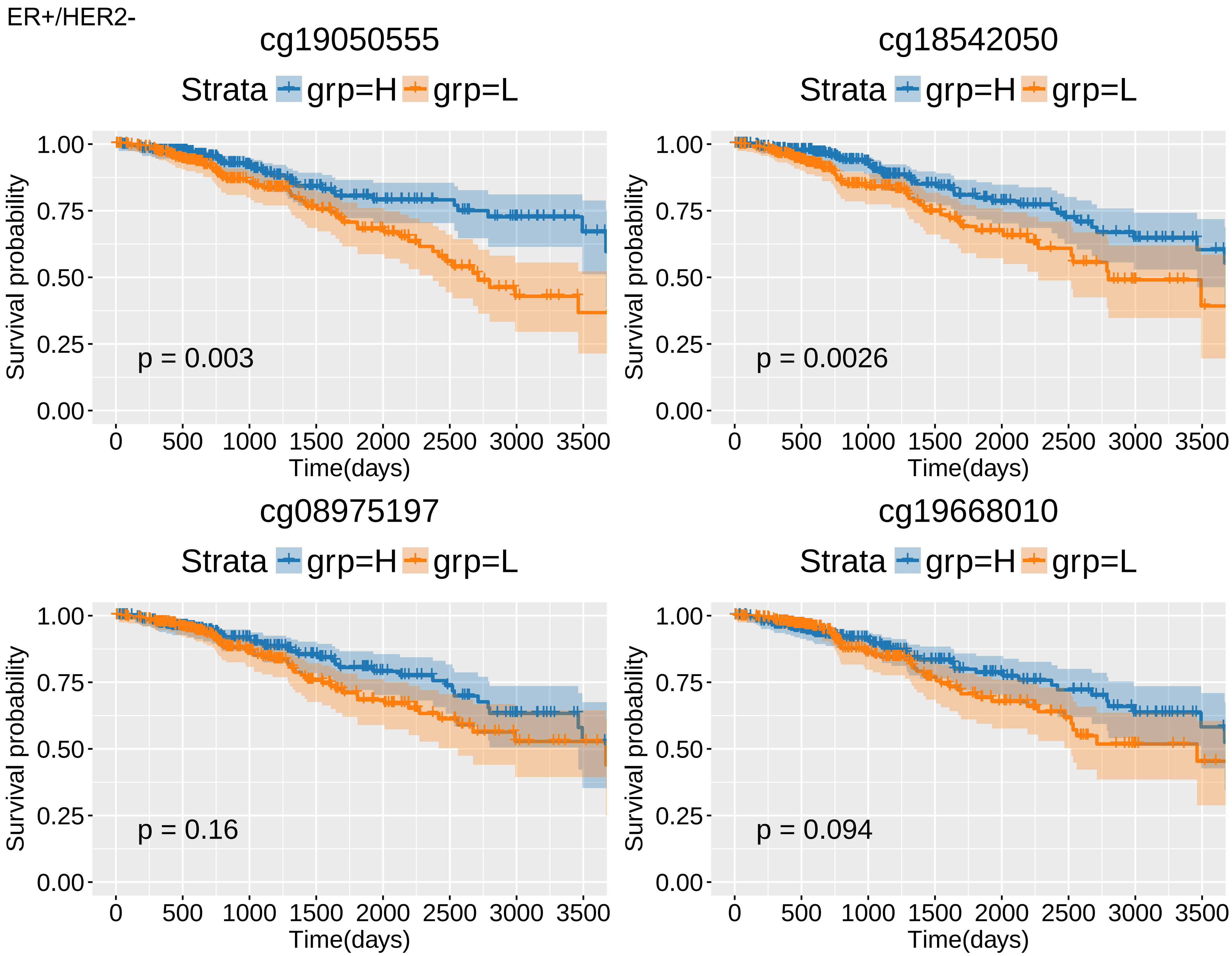
ER+/HER2-
cg19050555
Strata grp=H grp=L
p = 0.003
cg18542050
Strata grp=H grp=L
p = 0.0026
cg08975197
Strata grp=H grp=L
p = 0.16
cg19668010
Strata grp=H grp=L
p = 0.094
Survival probability
Time(days)
1.00
0.75
0.50
0.25
0.00
0
500
1000
1500
2000
2500
3000
3500

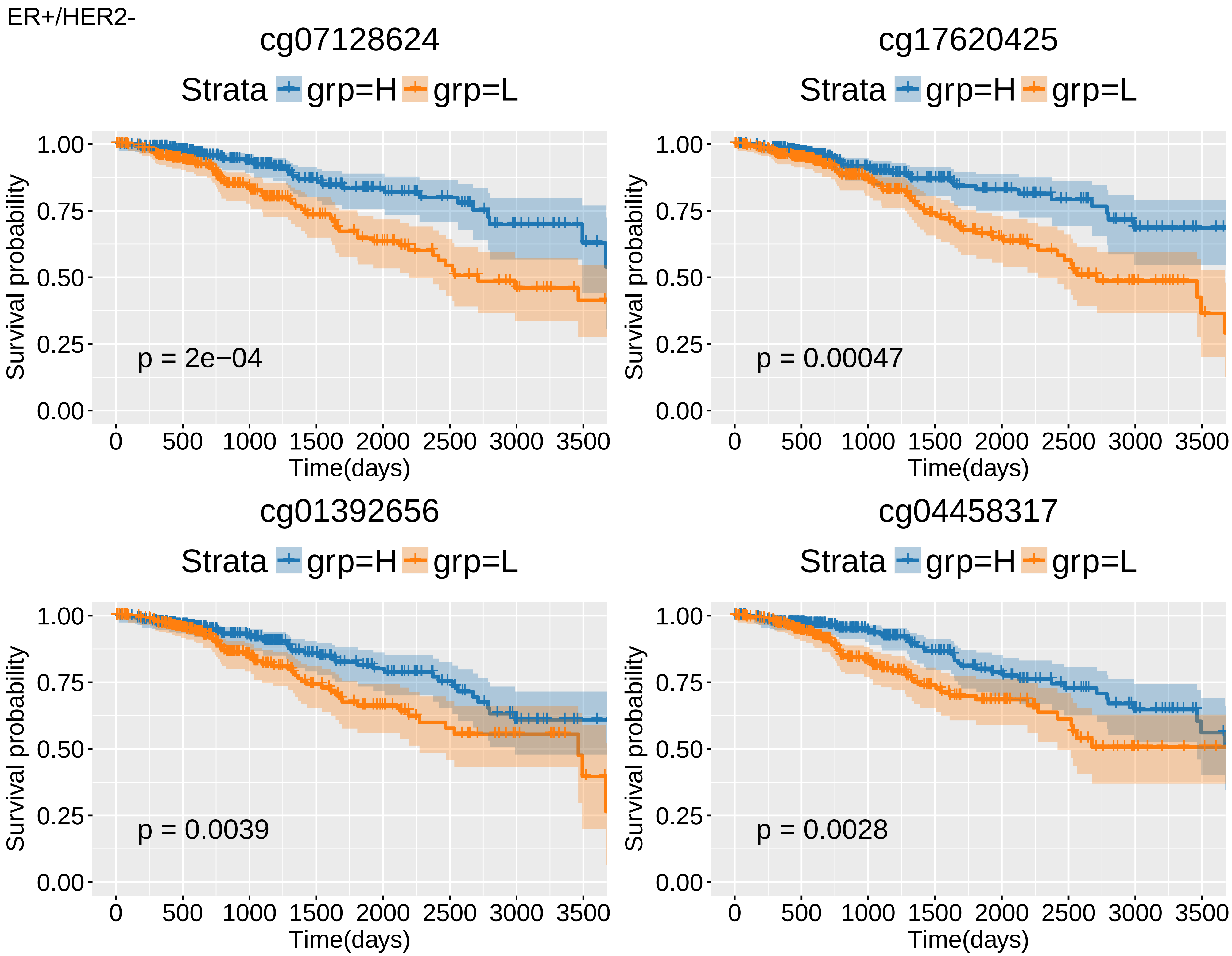
ER+/HER2-
cg07128624
Strata grp=H grp=L
p = 2e−04
cg17620425
Strata grp=H grp=L
p = 0.00047
cg01392656
Strata grp=H grp=L
p = 0.0039
cg04458317
Strata grp=H grp=L
p = 0.0028
Survival probability
1.00
0.75
0.50
0.25
0.00
Time(days)
0 500 1000 1500 2000 2500 3000 3500

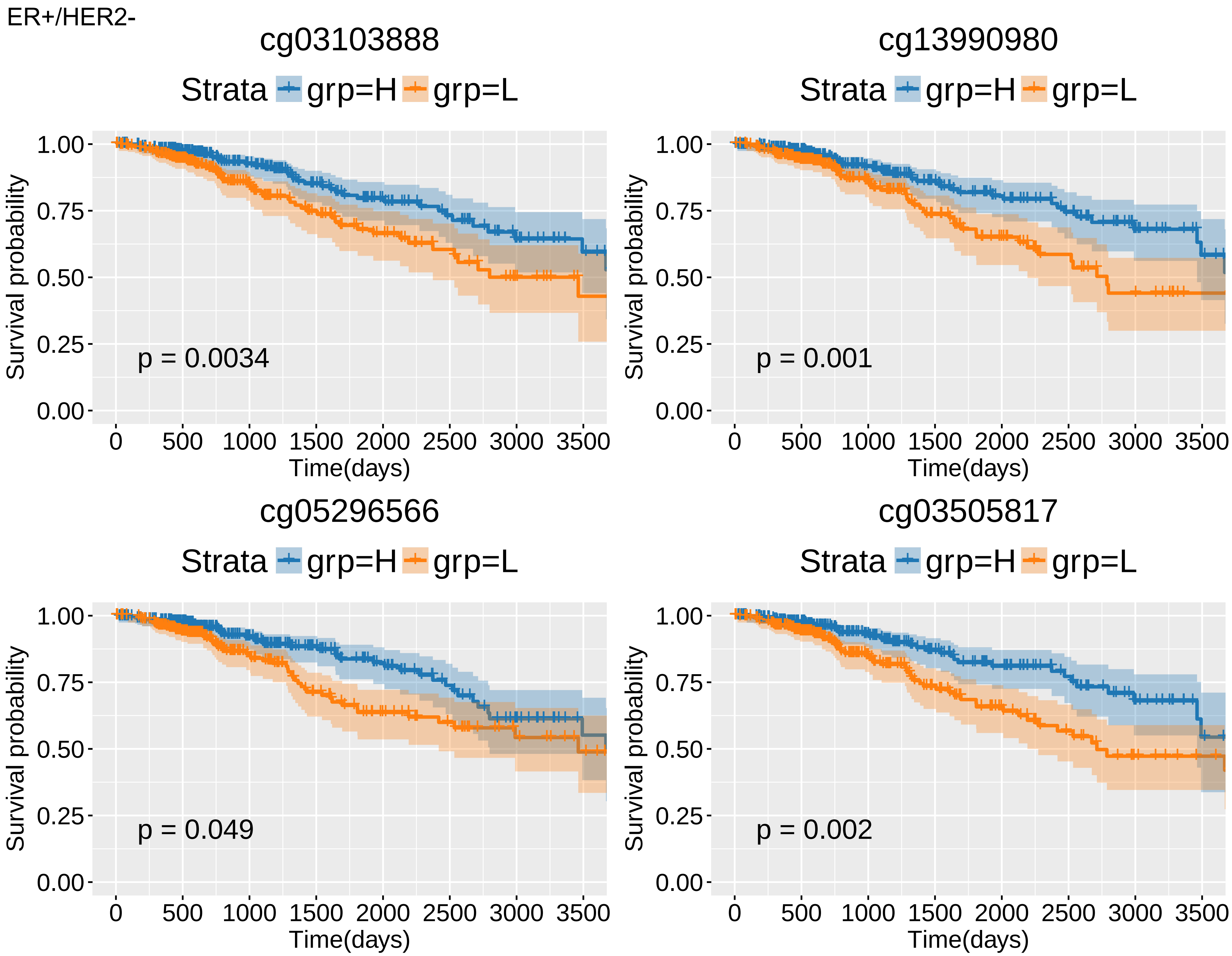
ER+/HER2-
cg03103888
Strata grp=H grp=L
Survival probability
1.00
0.75
0.50
0.25
0.00
p = 0.0034
0 500 1000 1500 2000 2500 3000 3500
Time(days)
cg13990980
Strata grp=H grp=L
Survival probability
1.00
0.75
0.50
0.25
0.00
p = 0.001
0 500 1000 1500 2000 2500 3000 3500
Time(days)
cg05296566
Strata grp=H grp=L
Survival probability
1.00
0.75
0.50
0.25
0.00
p = 0.049
0 500 1000 1500 2000 2500 3000 3500
Time(days)
cg03505817
Strata grp=H grp=L
Survival probability
1.00
0.75
0.50
0.25
0.00
p = 0.002
0 500 1000 1500 2000 2500 3000 3500
Time(days)

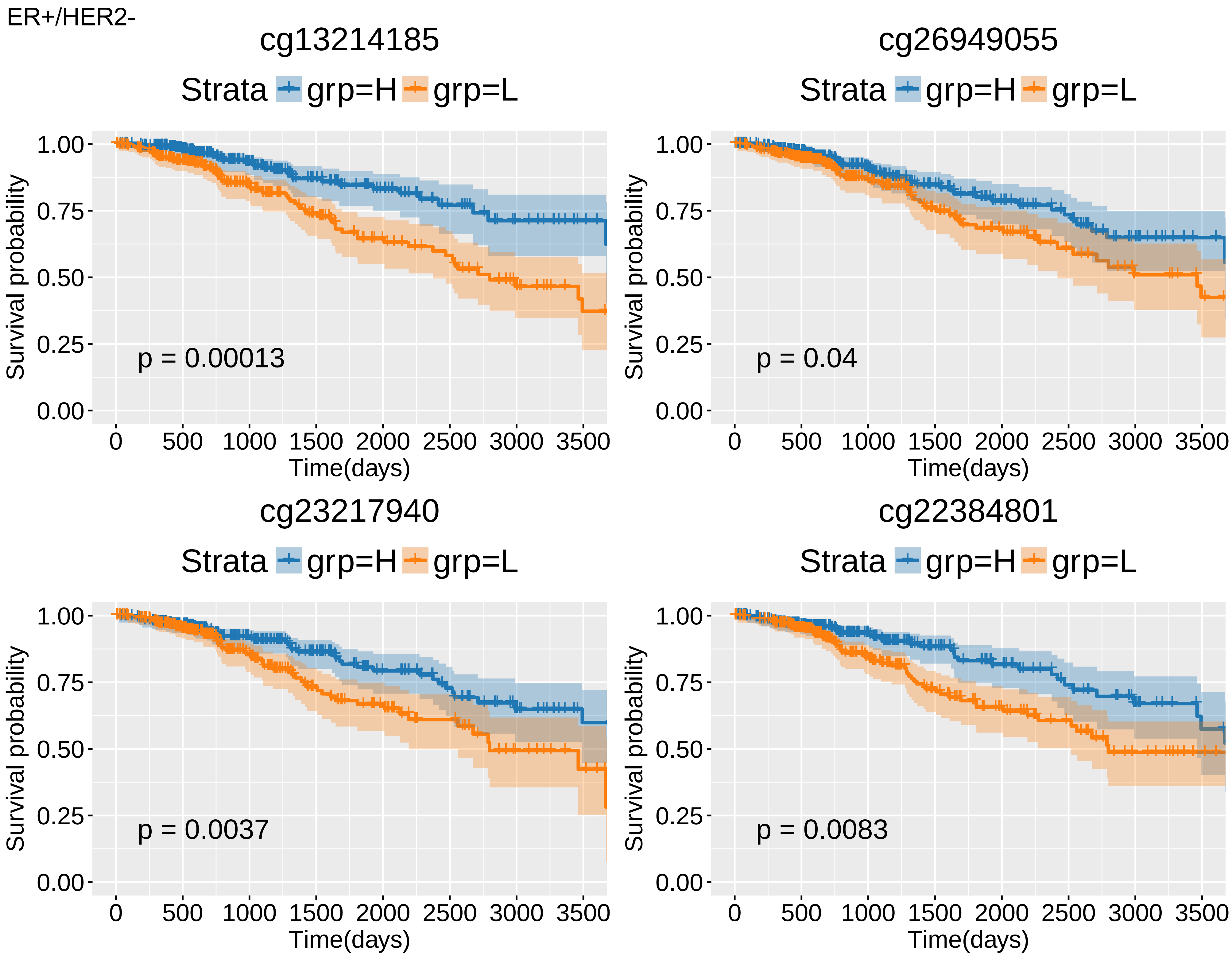

ER+/HER2-
cg13214185
Strata grp=H grp=L
Survival probability
1.00
0.75
0.50
0.25
0.00
p = 0.00013
0 500 1000 1500 2000 2500 3000 3500
Time(days)
cg26949055
Strata grp=H grp=L
p = 0.04
Time(days)
cg23217940
Strata grp=H grp=L
p = 0.0037
Time(days)
cg22384801
Strata grp=H grp=L
p = 0.0083
Time(days)

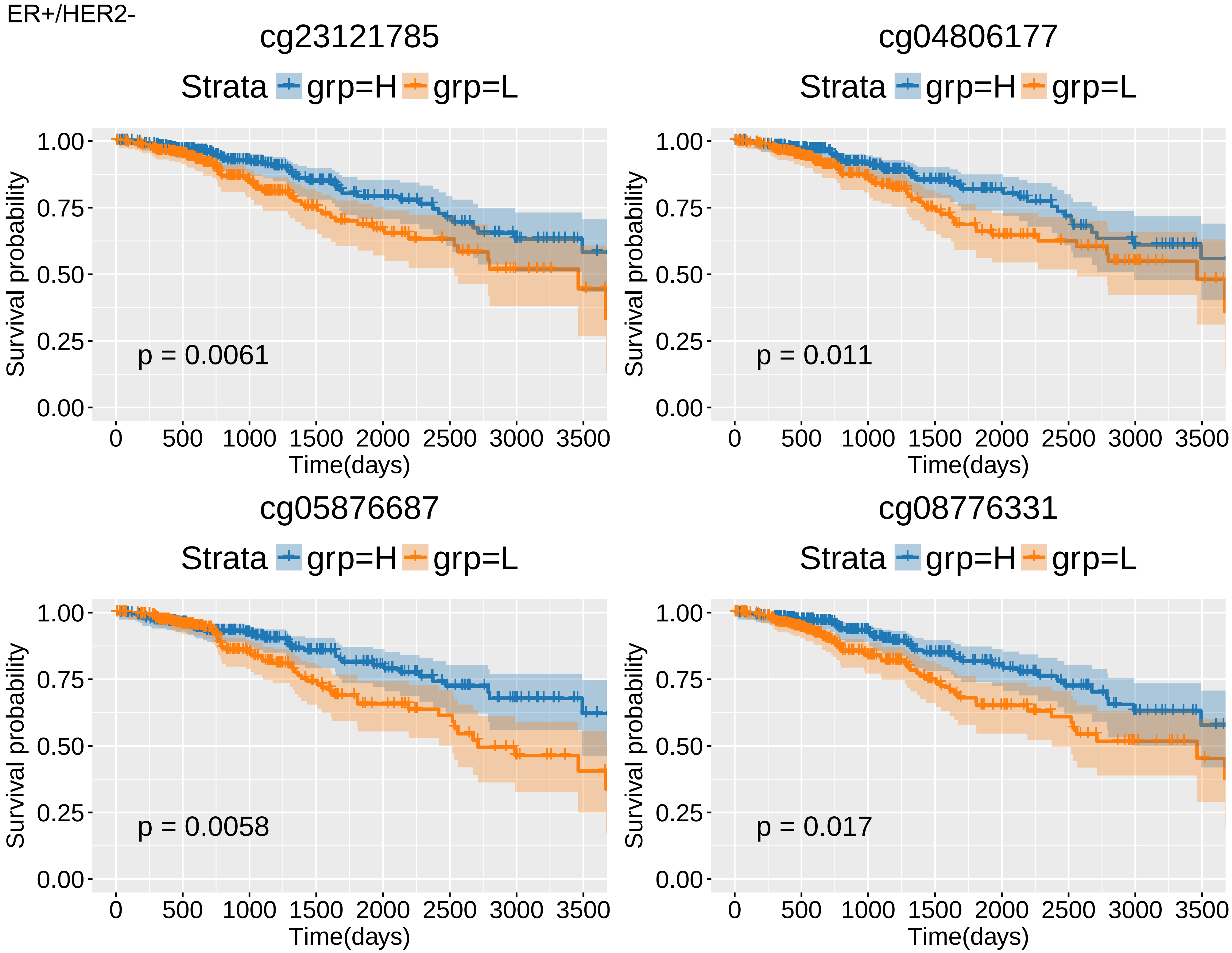
ER+/HER2-
cg23121785
Strata
grp=H
grp=L
Survival probability
1.00
0.75
0.50
0.25
0.00
p = 0.0061
0
500
1000
1500
2000
2500
3000
3500
Time(days)
cg04806177
Strata
grp=H
grp=L
Survival probability
p = 0.011
Time(days)
cg05876687
Strata
grp=H
grp=L
Survival probability
p = 0.0058
Time(days)
cg08776331
Strata
grp=H
grp=L
Survival probability
p = 0.017
Time(days)

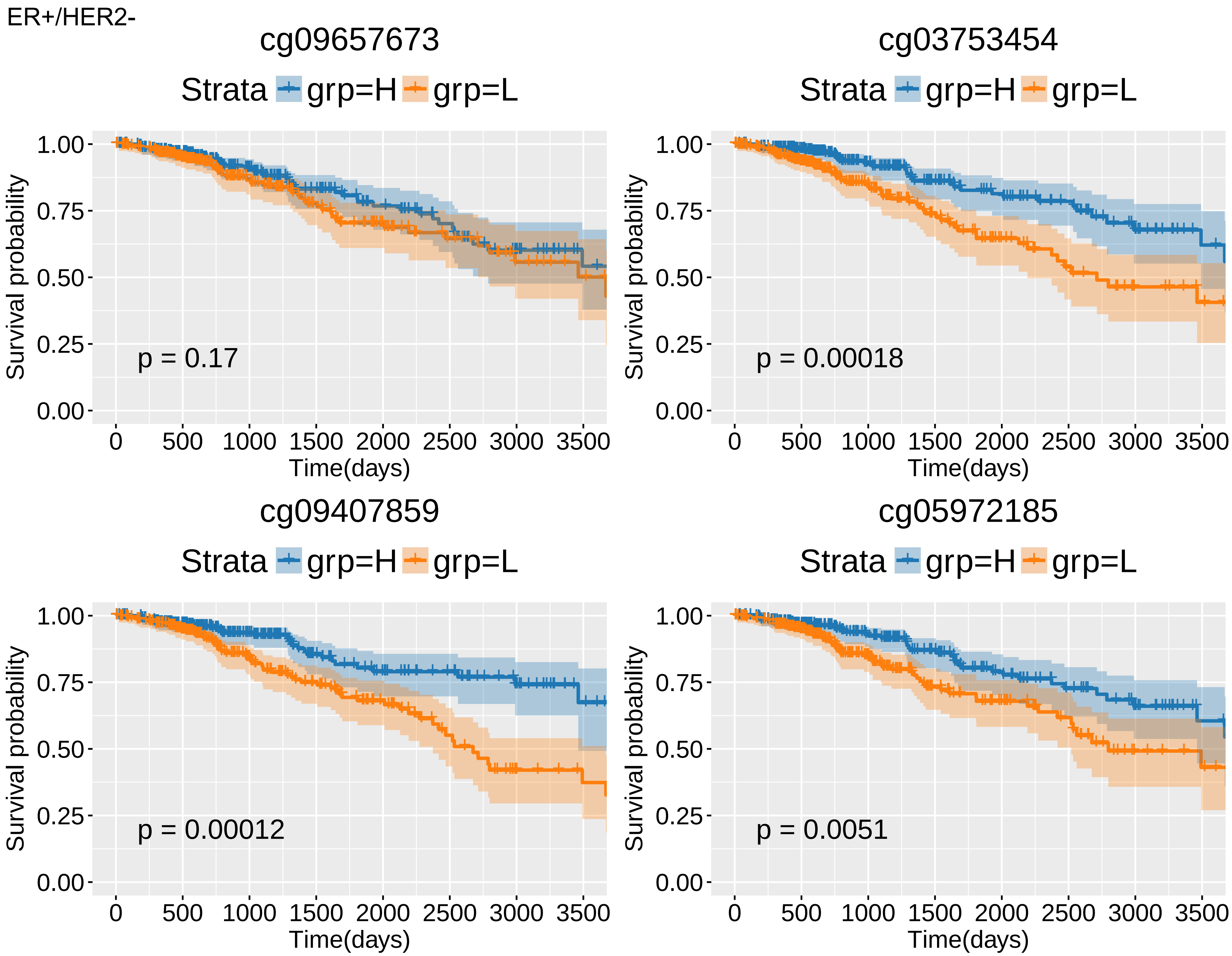
ER+/HER2-
cg09657673
Strata grp=H grp=L
p = 0.17
cg03753454
Strata grp=H grp=L
p = 0.00018
cg09407859
Strata grp=H grp=L
p = 0.00012
cg05972185
Strata grp=H grp=L
p = 0.0051
Survival probability
1.00
0.75
0.50
0.25
0.00
Time(days)
0 500 1000 1500 2000 2500 3000 3500

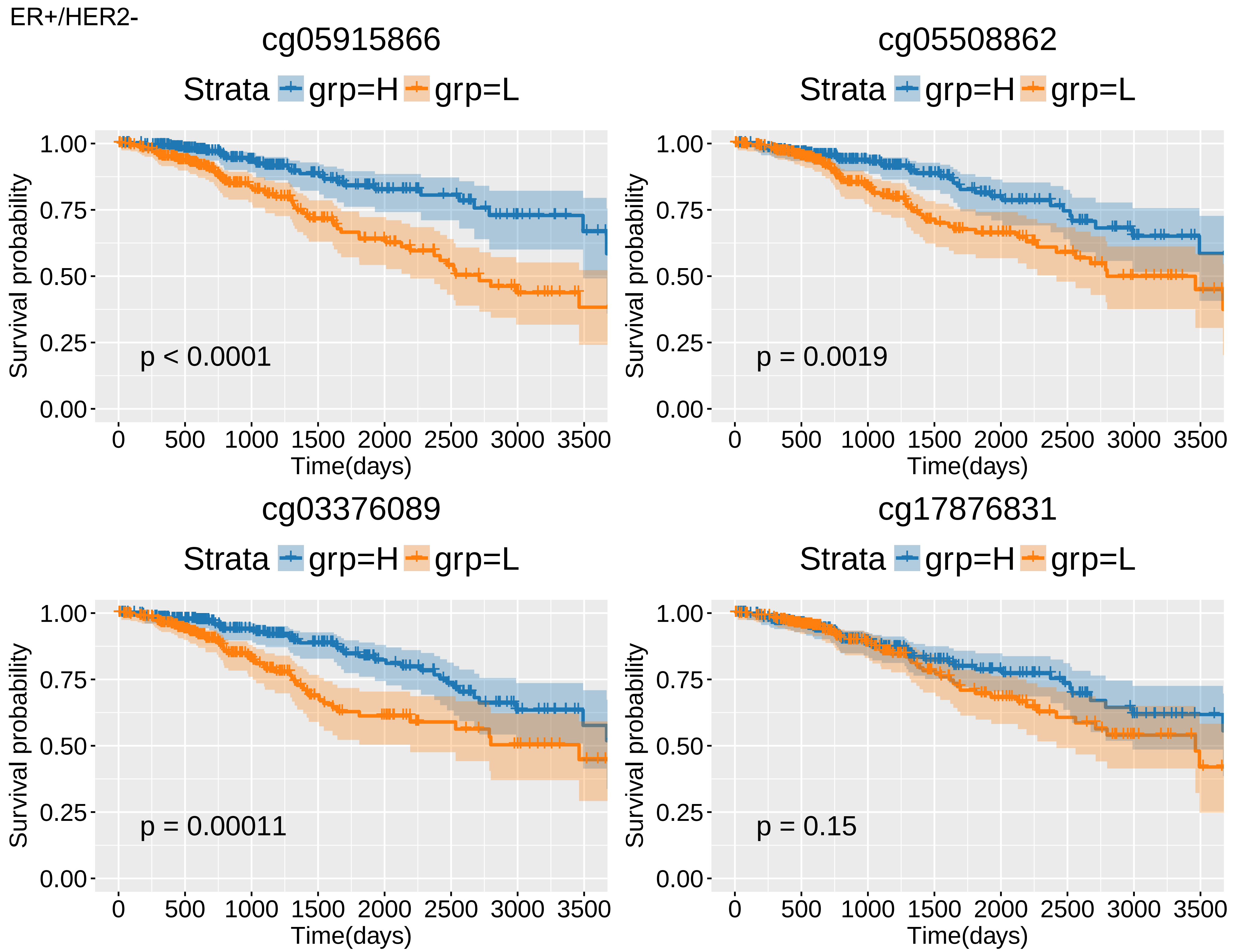
ER+/HER2-
cg05915866
Strata grp=H grp=L
p < 0.0001
cg05508862
Strata grp=H grp=L
p = 0.0019
cg03376089
Strata grp=H grp=L
p = 0.00011
cg17876831
Strata grp=H grp=L
p = 0.15
Survival probability
1.00
0.75
0.50
0.25
0.00
Time(days)
0
500
1000
1500
2000
2500
3000
3500

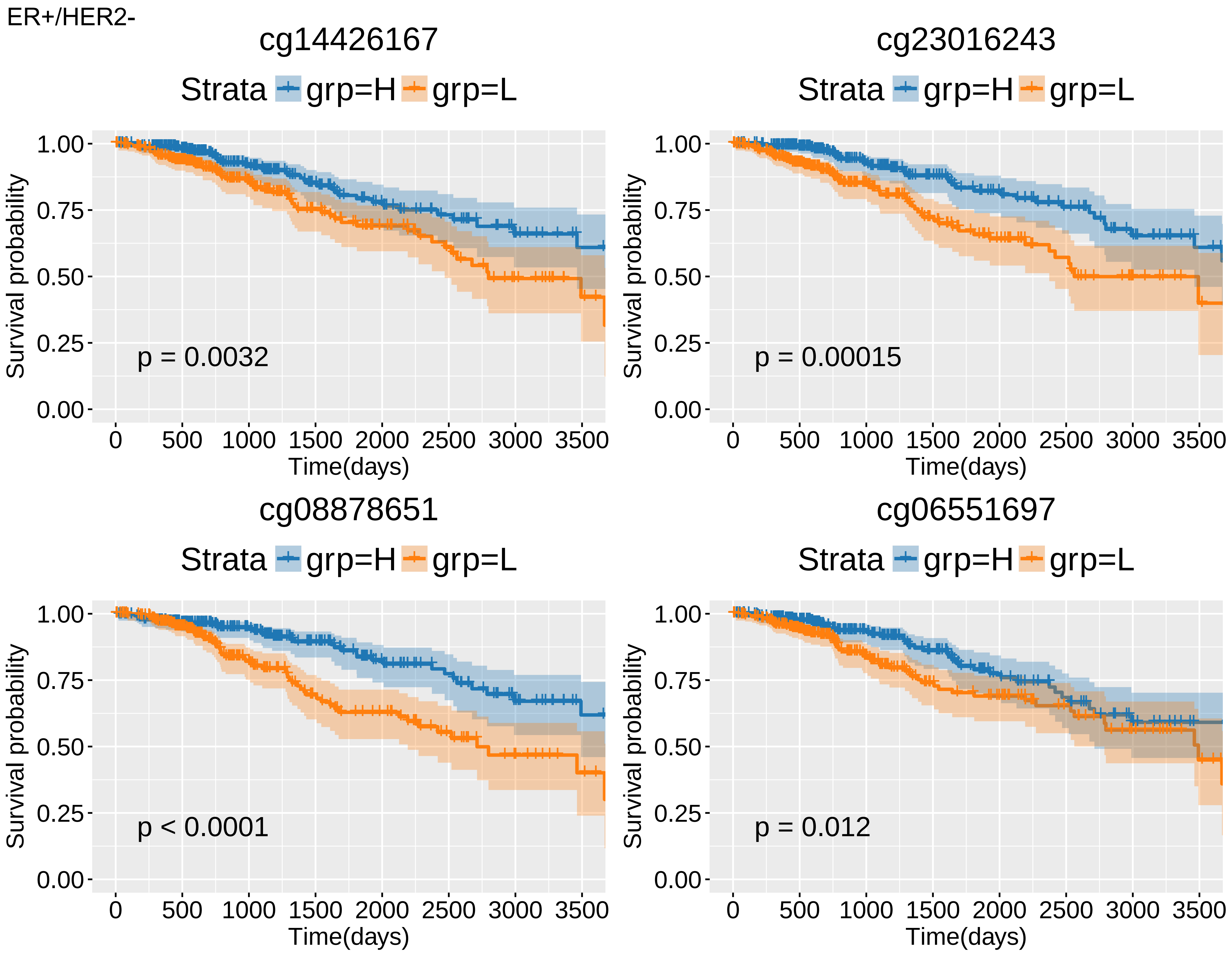
ER+/HER2-
cg14426167
Strata grp=H grp=L
p = 0.0032
cg23016243
Strata grp=H grp=L
p = 0.00015
cg08878651
Strata grp=H grp=L
p < 0.0001
cg06551697
Strata grp=H grp=L
p = 0.012
Survival probability
1.00
0.75
0.50
0.25
0.00
0
500
1000
1500
2000
2500
3000
3500
Time(days)

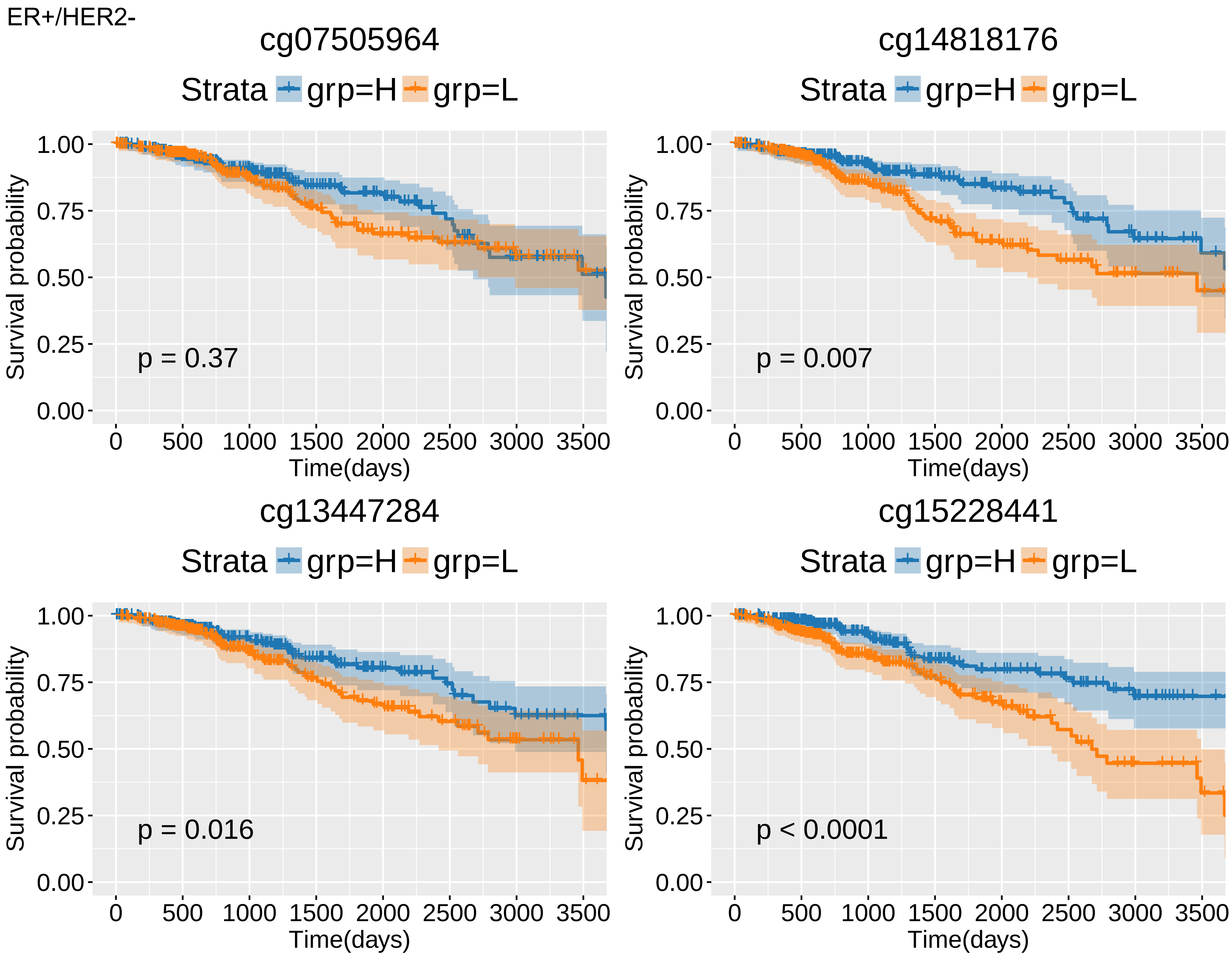
ER+/HER2-
cg07505964
Strata grp=H grp=L
Survival probability
1.00
0.75
0.50
0.25
0.00
p = 0.37
0 500 1000 1500 2000 2500 3000 3500
Time(days)
cg14818176
Strata grp=H grp=L
Survival probability
1.00
0.75
0.50
0.25
0.00
p = 0.007
0 500 1000 1500 2000 2500 3000 3500
Time(days)
cg13447284
Strata grp=H grp=L
Survival probability
1.00
0.75
0.50
0.25
0.00
p = 0.016
0 500 1000 1500 2000 2500 3000 3500
Time(days)
cg15228441
Strata grp=H grp=L
Survival probability
1.00
0.75
0.50
0.25
0.00
p < 0.0001
0 500 1000 1500 2000 2500 3000 3500
Time(days)

ER+/HER2-

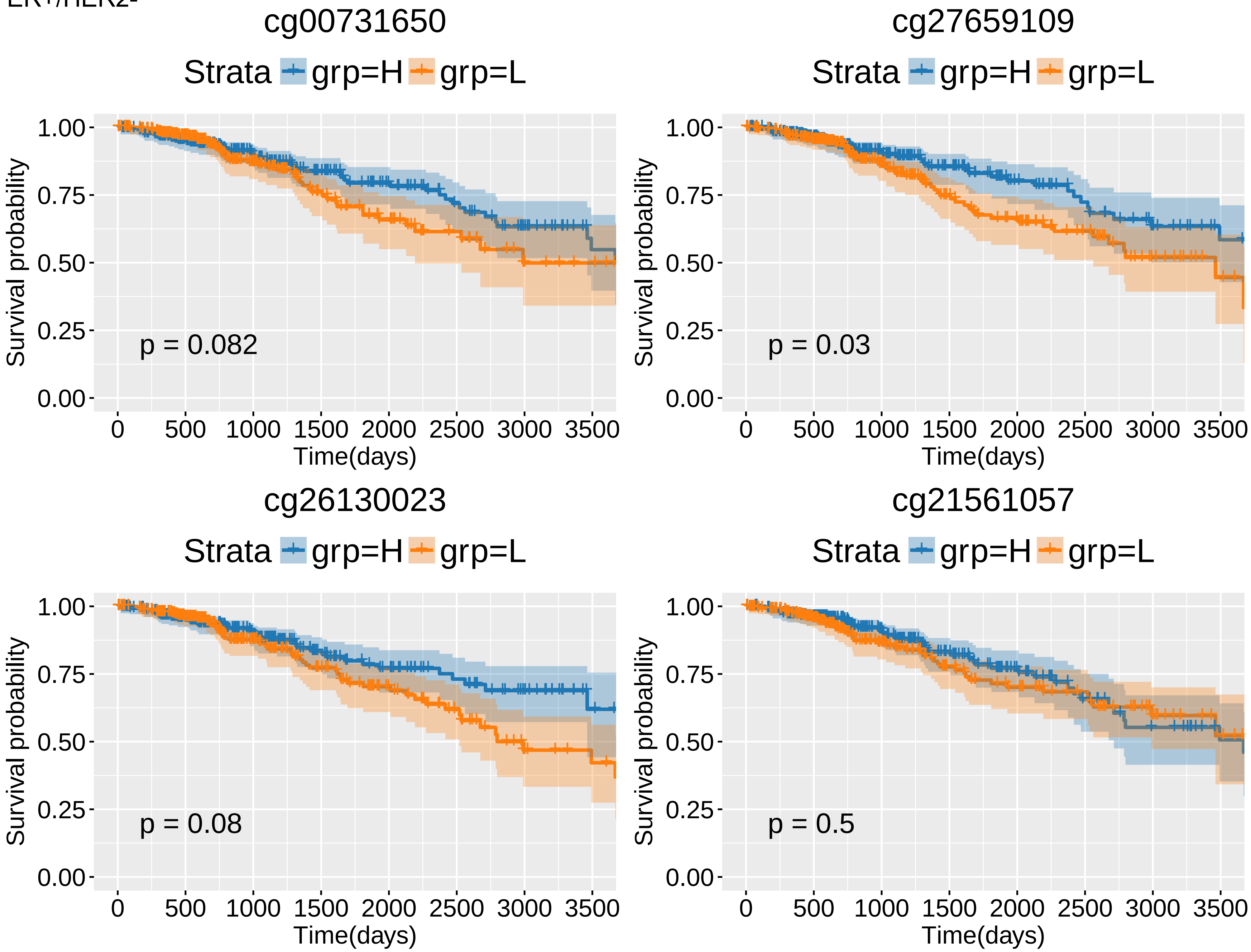

cg00731650
Strata grp=H grp=L
Survival probability
1.00
0.75
0.50
0.25
0.00
p = 0.082
0 500 1000 1500 2000 2500 3000 3500
Time(days)
cg27659109
Strata grp=H grp=L
Survival probability
p = 0.03
Time(days)
cg26130023
Strata grp=H grp=L
Survival probability
p = 0.08
Time(days)
cg21561057
Strata grp=H grp=L
Survival probability
p = 0.5
Time(days)

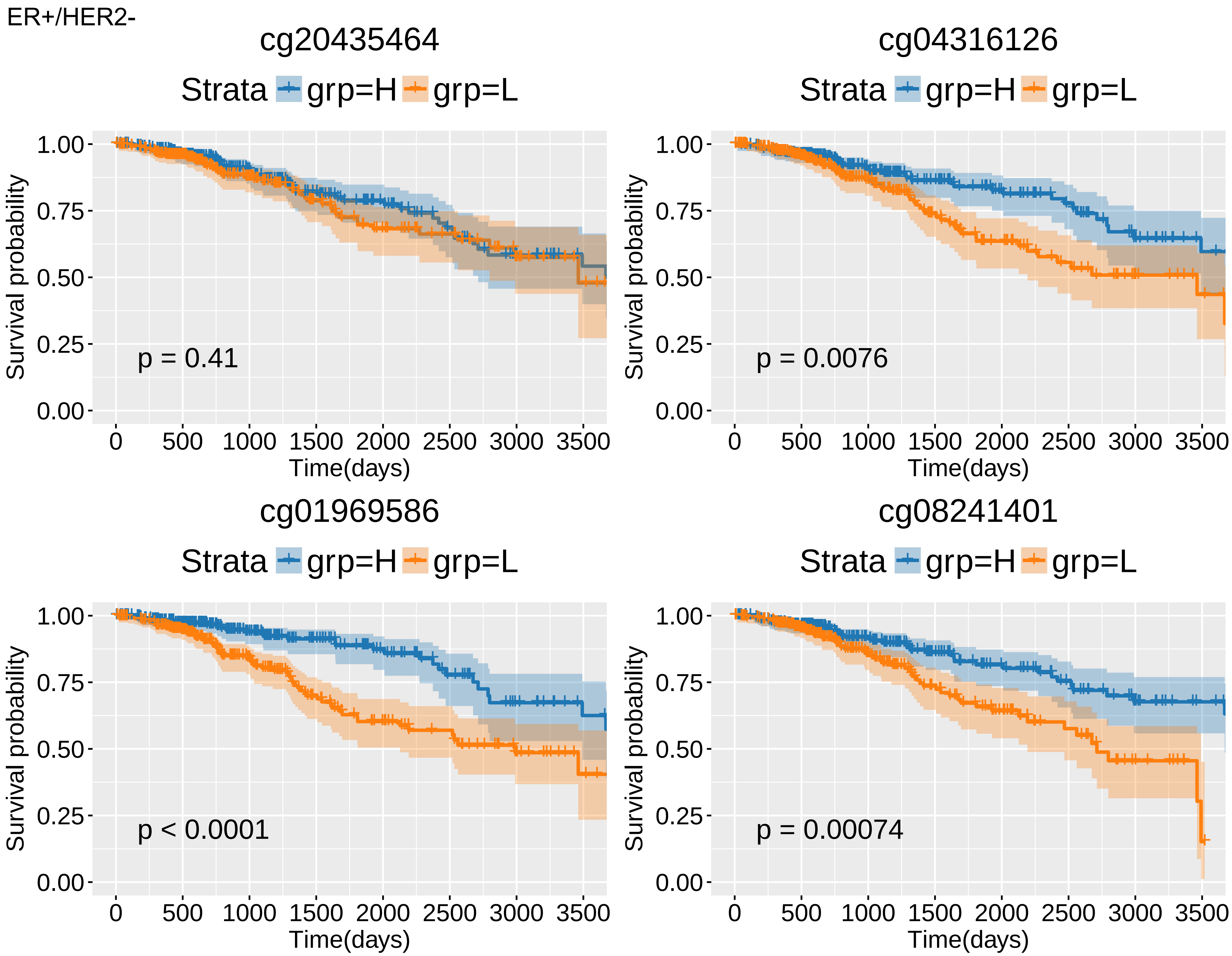
ER+/HER2-
cg20435464
Strata grp=H grp=L
Survival probability
1.00
0.75
0.50
0.25
0.00
p = 0.41
0 500 1000 1500 2000 2500 3000 3500
Time(days)
cg04316126
Strata grp=H grp=L
Survival probability
p = 0.0076
Time(days)
cg01969586
Strata grp=H grp=L
Survival probability
p < 0.0001
Time(days)
cg08241401
Strata grp=H grp=L
Survival probability
p = 0.00074
Time(days)

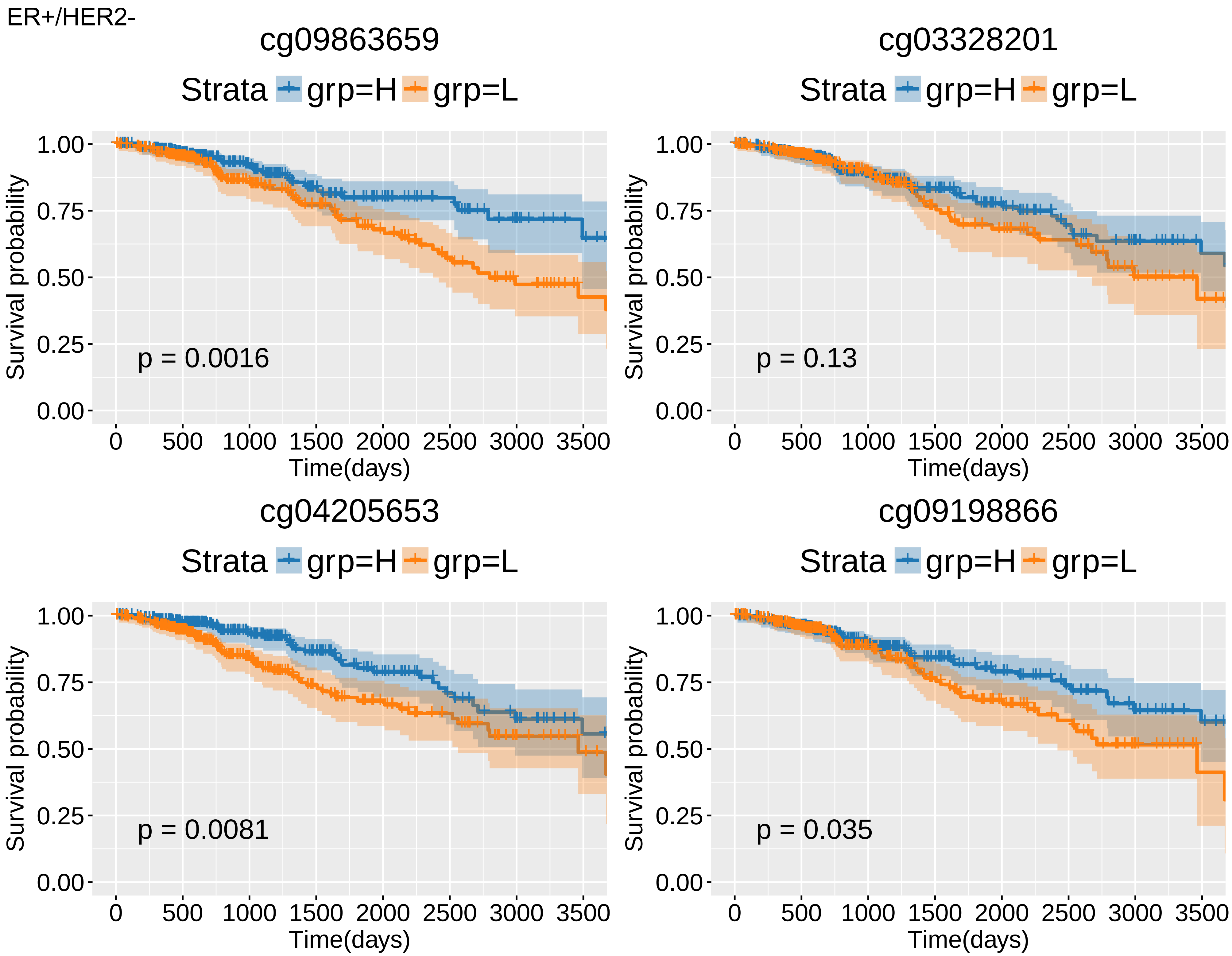
ER+/HER2-
cg09863659
Strata grp=H grp=L
p = 0.0016
cg03328201
Strata grp=H grp=L
p = 0.13
cg04205653
Strata grp=H grp=L
p = 0.0081
cg09198866
Strata grp=H grp=L
p = 0.035
Survival probability
1.00
0.75
0.50
0.25
0.00
Time(days)
0
500
1000
1500
2000
2500
3000
3500

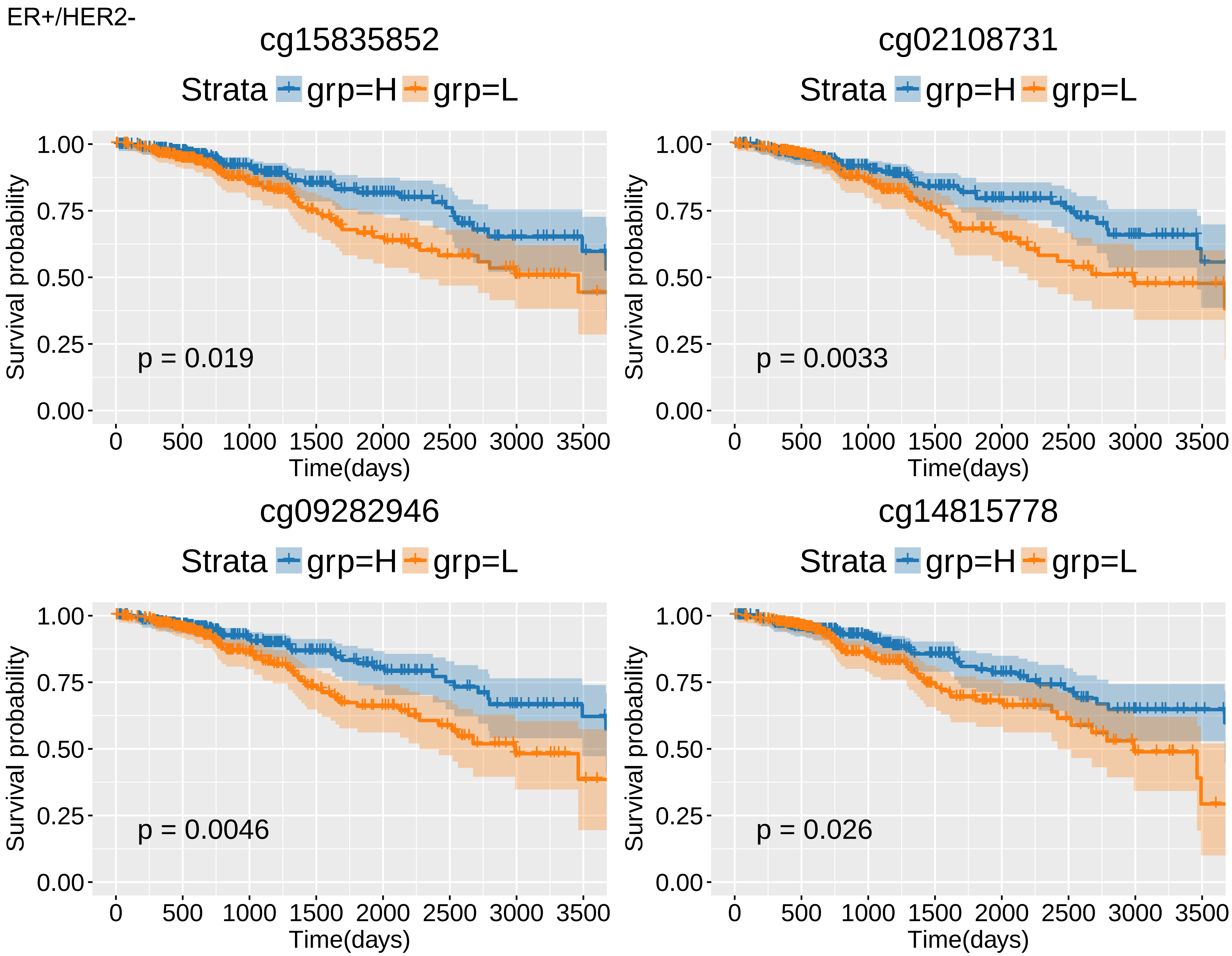
ER+/HER2-
cg15835852
Strata grp=H grp=L
p = 0.019
cg02108731
Strata grp=H grp=L
p = 0.0033
cg09282946
Strata grp=H grp=L
p = 0.0046
cg14815778
Strata grp=H grp=L
p = 0.026
Survival probability
Time(days)
1.00
0.75
0.50
0.25
0.00
0
500
1000
1500
2000
2500
3000
3500

TAM

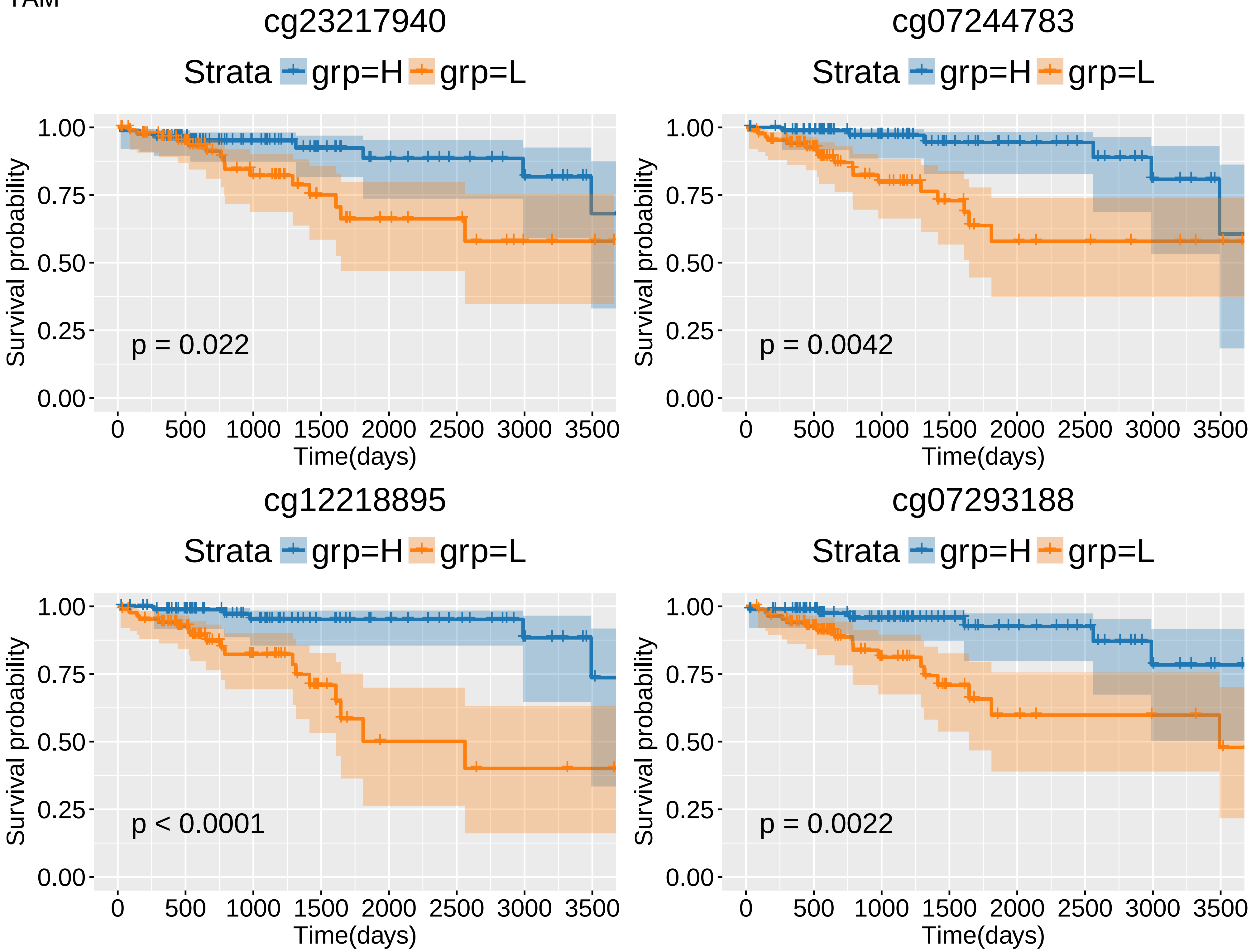
cg23217940
Strata grp=H grp=L
Survival probability
1.00
0.75
0.50
0.25
0.00
p = 0.022
0 500 1000 1500 2000 2500 3000 3500
Time(days)
cg07244783
Strata grp=H grp=L
Survival probability
p = 0.0042
Time(days)
cg12218895
Strata grp=H grp=L
Survival probability
p < 0.0001
Time(days)
cg07293188
Strata grp=H grp=L
Survival probability
p = 0.0022
Time(days)

TAM

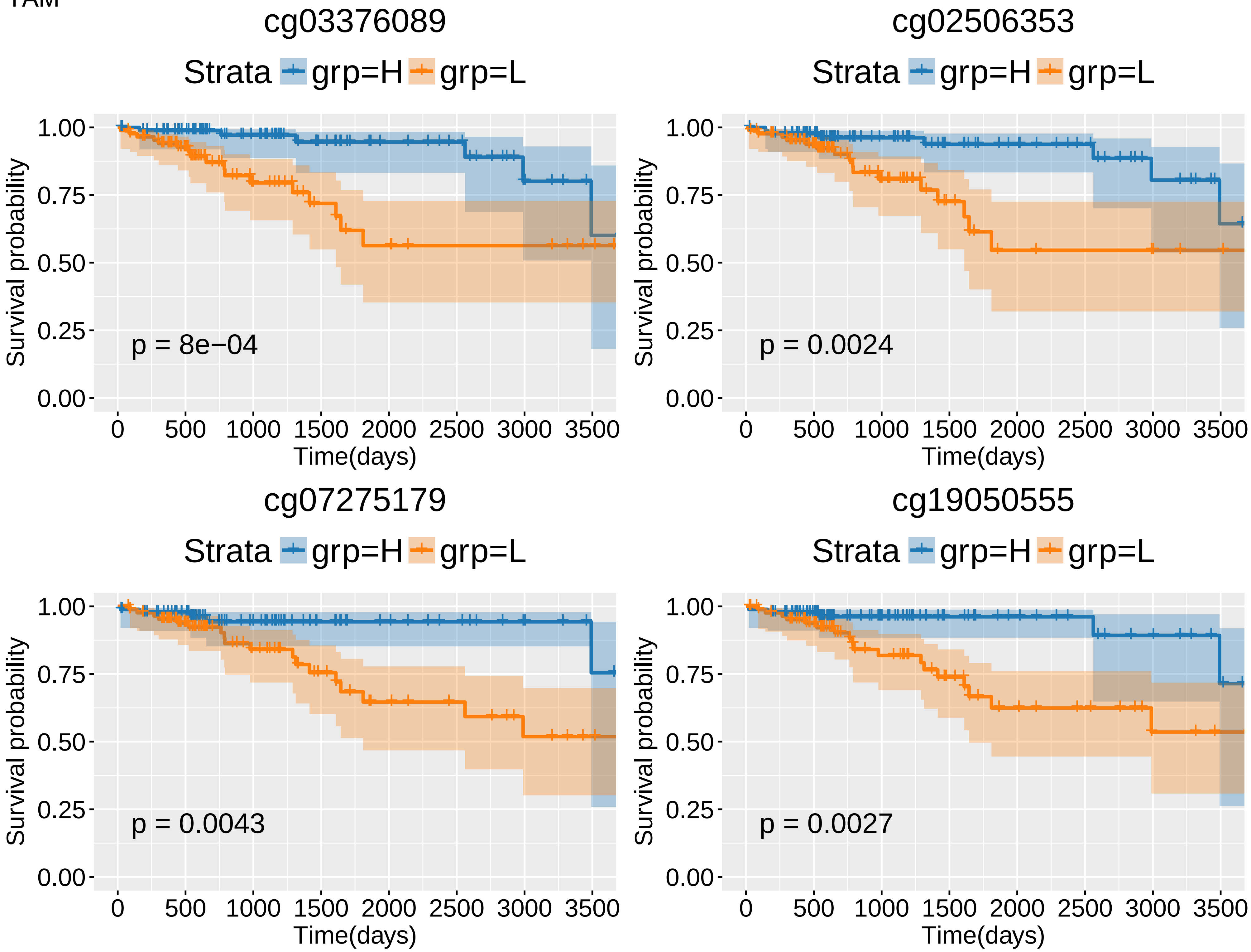
cg03376089
Strata grp=H grp=L
Survival probability
1.00
0.75
0.50
0.25
0.00
p = 8e−04
0 500 1000 1500 2000 2500 3000 3500
Time(days)
cg02506353
Strata grp=H grp=L
Survival probability
1.00
0.75
0.50
0.25
0.00
p = 0.0024
0 500 1000 1500 2000 2500 3000 3500
Time(days)
cg07275179
Strata grp=H grp=L
Survival probability
1.00
0.75
0.50
0.25
0.00
p = 0.0043
0 500 1000 1500 2000 2500 3000 3500
Time(days)
cg19050555
Strata grp=H grp=L
Survival probability
1.00
0.75
0.50
0.25
0.00
p = 0.0027
0 500 1000 1500 2000 2500 3000 3500
Time(days)

TAM

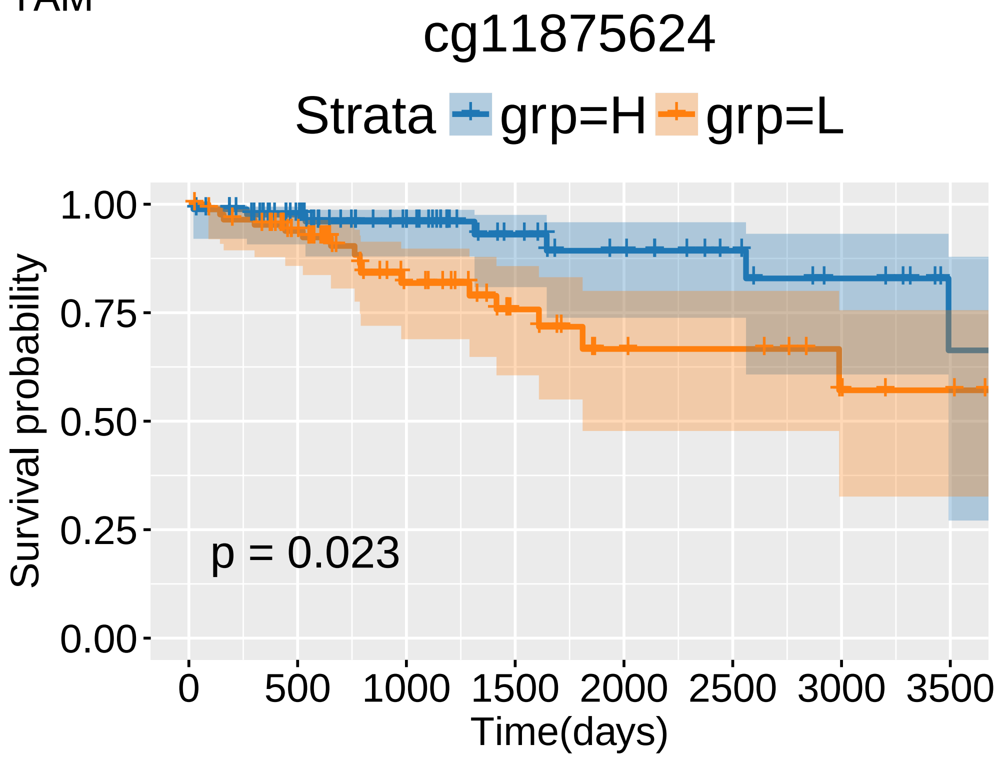
cg11875624
Strata
grp=H
grp=L
Survival probability
1.00
0.75
0.50
0.25
0.00
p = 0.023
0
500
1000
1500
2000
2500
3000
3500
Time(days)

AI

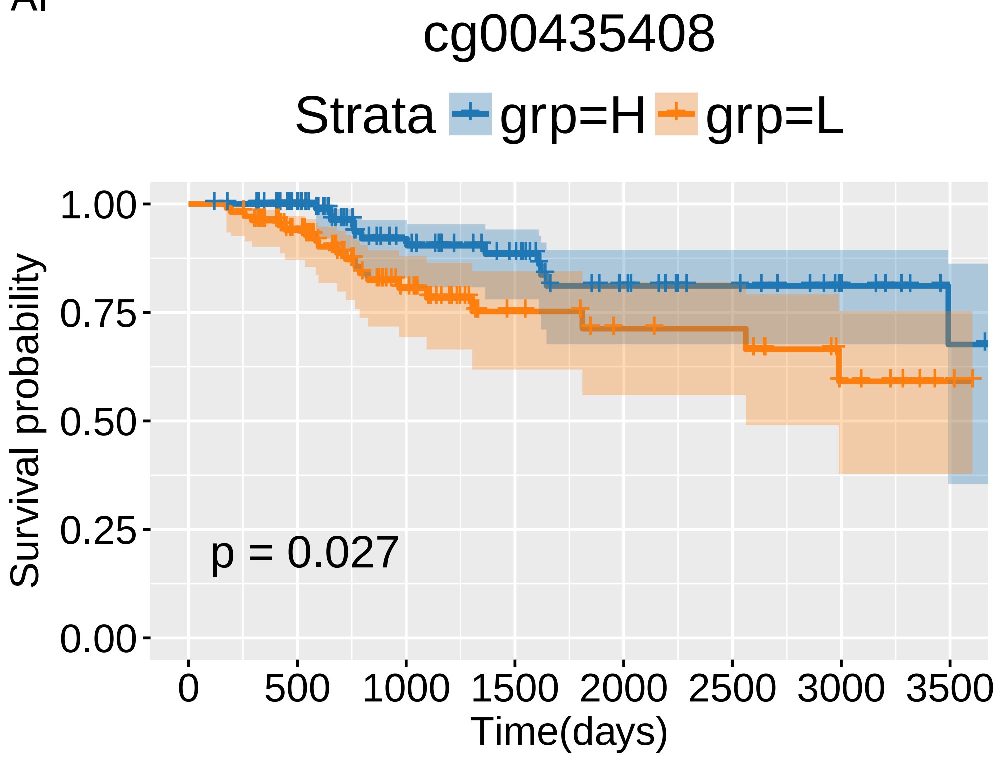
cg00435408
Strata grp=H grp=L
1.00
0.75
0.50
0.25
0.00
Survival probability
p = 0.027
0
500
1000
1500
2000
2500
3000
3500
Time(days)
