## Supplemental file 6 _ overlap for "Candidate methylation sites associated with endocrine therapy resistance in the TCGA ER+/HER2- breast cancer cohort"

### Slide 1
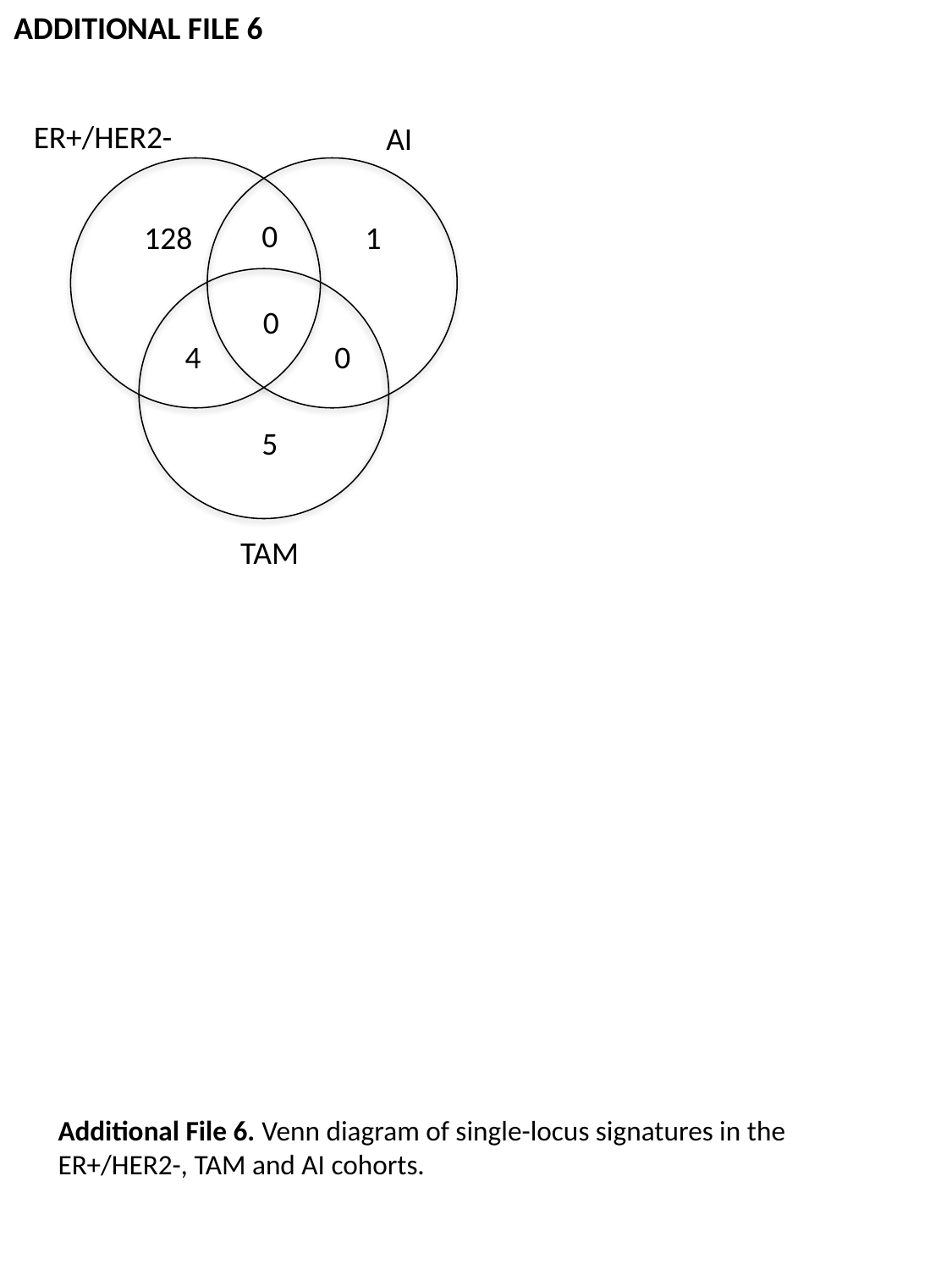

ADDITIONAL FILE 6
ER+/HER2-
AI
0
128
1
0
4
0
5
TAM
Additional File 6. Venn diagram of single-locus signatures in the ER+/HER2-, TAM and AI cohorts.
