## Supplemental file 9 _ Venn Diagrams for "Candidate methylation sites associated with endocrine therapy resistance in the TCGA ER+/HER2- breast cancer cohort"

### Slide 1
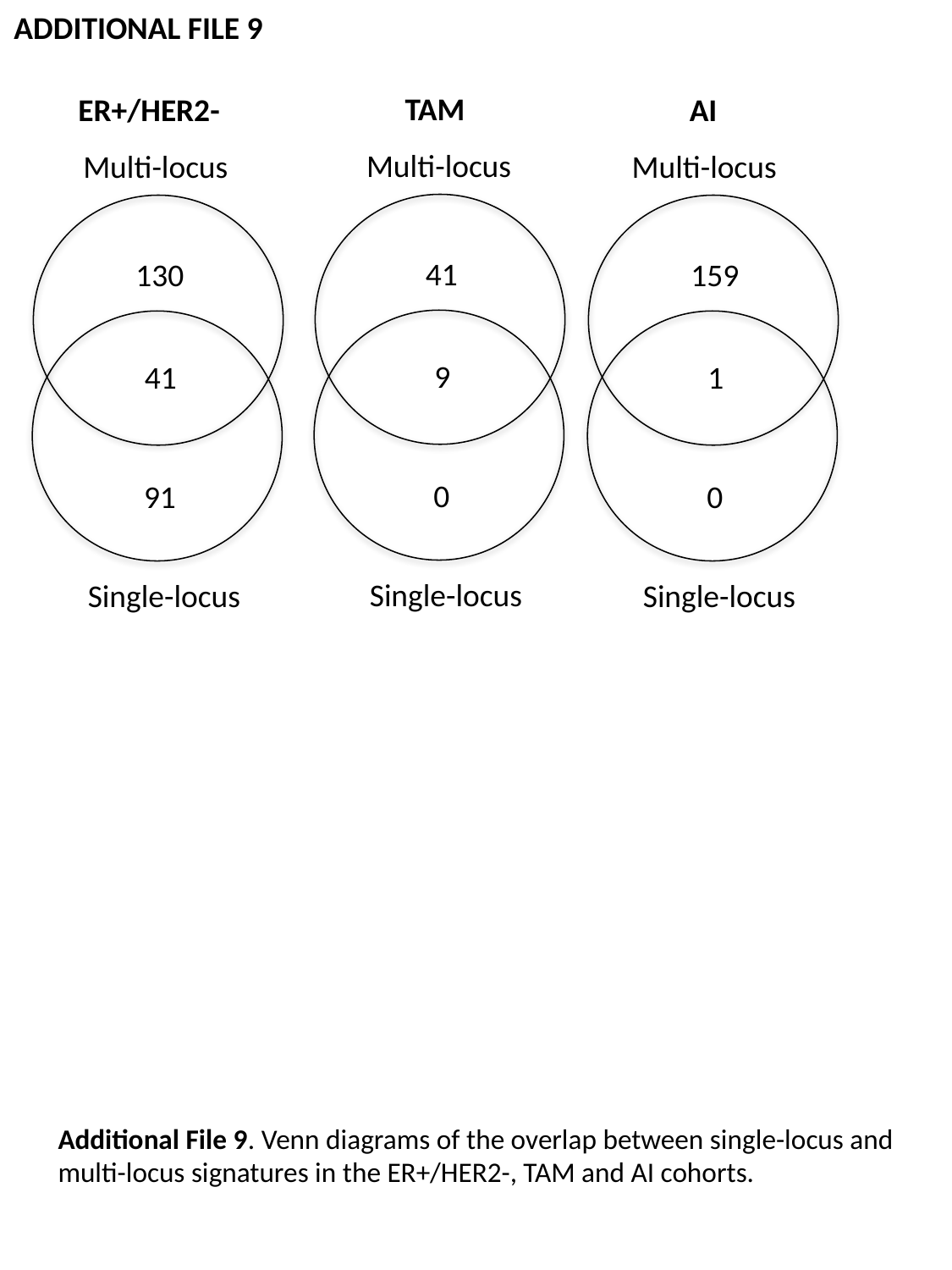

ADDITIONAL FILE 9
TAM
ER+/HER2-
AI
Multi-locus
Multi-locus
Multi-locus
41
130
159
9
41
1
0
91
0
Single-locus
Single-locus
Single-locus
Additional File 9. Venn diagrams of the overlap between single-locus and multi-locus signatures in the ER+/HER2-, TAM and AI cohorts.
